# Small Molecule-Directed RNA Modification via Proximity-Driven Catalysis

**DOI:** 10.64898/2026.05.31.729146

**Authors:** Song Chen, Tuan-Khoa Kha, Yiran Zhao, Junsong Guo, Baixi Chen, Ru-Yi Zhu

## Abstract

Selective chemical modification of RNA is essential for RNA functionalization, probing RNA structure-function relationships and developing RNA-targeted therapeutics. Existing chemical strategies often rely on guanine accessibility or multiple helper DNA strands, restricting their generality and biological applicability. Inspired by DNA-guided DMAP catalysis and small-molecule binding-induced crosslinking, we report a small molecule-directed, DMAP-catalyzed, proximity-driven strategy for site-selective RNA functionalization. By appending a catalytic DMAP moiety to RNA-binding ligands, 2′-OH groups are selectively acylated in the presence of azide-bearing acyl donors, enabling subsequent installation of bioorthogonal handles. This approach was validated across diverse RNAs, including Pepper and Clivia RNA aptamers, G-quadruplex Broccoli RNA, and endogenous FMN riboswitch RNA. For a 400-nt Pepper-7SK fusion, selective modification of the Pepper motif was achieved with minimal perturbation to the nucleus localization function of 7SK RNA. Optimized PEG-pentafluorophenyl (PFP) acyl donors provided enhanced reactivity and low background. The method operates catalytically, decouples ligand recognition from the labeling moiety, and enables selective enrichment of target RNAs, offering a versatile platform for RNA functionalization, ligand profiling, and potentially live-cell applications.

## Introduction

Selective chemical modification of RNA is essential for RNA functionalization, elucidating RNA structure-function relationships and identification of novel RNA-small molecule interactions.^1–2^ Chemical approaches have been developed that exploit complementary DNA strands as masking elements to achieve site-selective RNA modification using highly reactive small-molecule reagents. In these strategies, base-paired regions of RNA are sterically and electrostatically protected, while unpaired nucleotides remain selectively accessible.^3–5^ Accordingly, for short RNA substrates, hybridization of helper DNA to non-target regions-leaving the desired site in a loop or gap-has enabled site-selective 2′-OH acylation and guanine *O*^6^ alkylation.^5–7^ However, the efficiency and practicality of this approach diminish for long or highly structured RNAs as it relies on many helper DNA strands to cover the entire RNA sequence except for the intended reaction site. To overcome this limitation, we and the Kool group independently reported DNA-guided catalysis, in which 4-dimethylaminopyridine (DMAP) tethered to a guide DNA enables selective acyl or aryl transfer to 2’-OH at predetermined sites.^8–9^

An alternative approach to achieve selective RNA modification leverages ligand binding-induced proximity reactivity of highly reactive functional groups. In this strategy, RNA-binding small molecules are covalently tethered to reactive moieties, enabling covalent crosslinking to RNA (**Figure 1**, middle). Proximity-driven, small-molecule binding-induced covalent labeling is well established in the protein field and underpins activity-based protein profiling.^10–11^ Extending this concept to RNA, Disney, Schneekloth, Velema, and others developed diazirine-bearing RNA-binding small molecules that, upon photoactivation, form covalent adducts with their target RNAs. Incorporation of click handles enables subsequent enrichment and identification of bound RNAs, an approach termed Chem-CLIP (Chemical Cross-Linking and Isolation by Pull-down).^1, 12–17^ Despite its utility, the reliance on short-wavelength UV irradiation and low crosslink efficiency raise concerns regarding RNA damage and restrict applicability in biological settings.^18^

**Figure 1.**
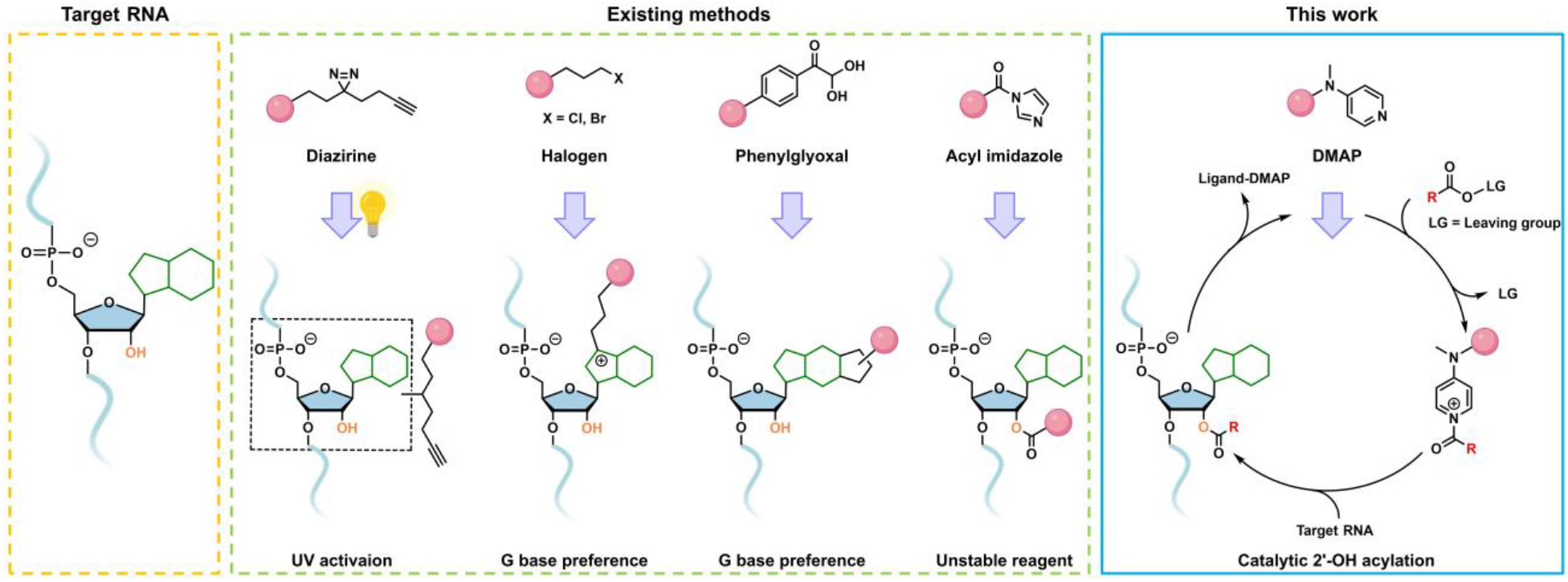
Representative strategies for covalent crosslinking of target RNAs and the ligand-tethered DMAP-based catalytic modification strategy (this work). Pink ball represents an RNA-binding small molecule.

Additional covalent RNA-targeting strategies have been reported by Disney, Micura, and others, including phenylglyoxal-derived and alkyl halide-based electrophiles.^2, 19–20^ Phenylglyoxal reagents selectively react with the nucleophilic *N*1 and *N*2 positions on the Watson-Crick face of unpaired guanine residues, forming cyclized adducts. In contrast, alkyl halides preferentially alkylate the *N*7 position of guanine bases proximal to ligand-binding sites. Although effective for capturing ligand-RNA interactions, their reactivity is strongly dependent on the presence and spatial positioning of guanine residues, thereby limiting their general applicability. Furthermore, the highly electrophilic feature of these probes poses significant challenges for their synthetic derivatization, restricting downstream applications.

Unlike aforementioned guanine-selective electrophiles, acylation of ubiquitous RNA 2′-OH groups by acyl imidazole reagents offers a more general approach for RNA modification.^21^ In particular, acyl imidazole warheads have been appended to RNA-binding small molecules to enable proximity-driven RNA modification for profiling RNA-small molecule interactions (**Figure 1**, middle).^22–24^ However, the high intrinsic reactivity of these electrophiles often compromises reagent stability and necessitates elevated concentrations to compete with hydrolysis and potential off-target reactions with proteins.

Inspired by the DNA-guided DMAP catalysis platform and small molecule binding-induced RNA crosslinking, we reasoned that RNA-binding small molecules if tethered with DMAP could offer a complementary strategy for selective RNA modification particularly suited for folded RNAs. Here, we report a small molecule-directed, DMAP-catalyzed proximity-driven acylation approach that exploits intrinsic RNA-ligand binding interactions to achieve site-selective RNA functionalization (**Figure 1**, right). We demonstrate the generality of this strategy across diverse RNA targets, including Pepper RNA aptamer, G-quadruplex (G4)-containing Broccoli RNA aptamer, and endogenous FMN riboswitch RNA. Our experimental results show labeling occurs at nucleotides proximal to the ligand-binding site, consistent with reported crystal structures and directly reports on ligand-RNA interactions. Consequently, it offers a useful strategy for probing RNA-ligand interactions, particularly for systems lacking high-resolution structural information. In the presence of azide-bearing acyl donors, this method enables the site-selective installation of bioorthogonal handles onto target RNAs, allowing subsequent derivatization. Notably, for a Pepper-7SK fusion RNA of approximately 400 nucleotides (nt), highly specific labeling of the full-length RNA is achieved by targeting the Pepper motif, with minimal perturbation of native RNA function. In this context, the short, ligand-recognizable Pepper sequence effectively acts as a molecular “Velcro,” enabling selective tagging of RNAs of interest. Finally, we further demonstrate that this proximity-based labeling platform enables enrichment of target RNA from total RNA via pull-down experiments, highlighting its potential as a small-molecule- or RNA-centric approach for identifying RNA-small-molecule interactions.

## Results and Discussion

### Design and evaluation of the HBC-DMAP catalyst for Pepper RNA

We selected the Pepper RNA aptamer as a model system to establish our proof of concept. Pepper RNA aptamers are a class of fluorogenic RNAs identified via SELEX (systematic evolution of ligands by exponential enrichment) that bind (4-((2-hydroxyrthyl)(methyl) amino)-benzylidene)-cyanophenyl-acetonitrile (HBC)-series fluorogens with high affinity. The resulting fluorophore-aptamer complexes exhibit tunable emission maxima across the cyan-to-red spectral range, depending on the specific fluorophore structure. Owing to these favorable photophysical properties, Pepper RNA aptamers have been widely employed for live-cell RNA imaging. In this study, we generated F30-scaffolded Pepper RNA by *in vitro* transcription (IVT) (**Figure. 2a**). HBC530 was chosen as the small-molecule ligand due to its exceptionally strong binding to Pepper RNA, with a dissociation constant of ~3.5 nM.^25^ This high-affinity interaction provides a robust platform for implementing affinity-guided, target-specific RNA modification.

**Figure 2.**
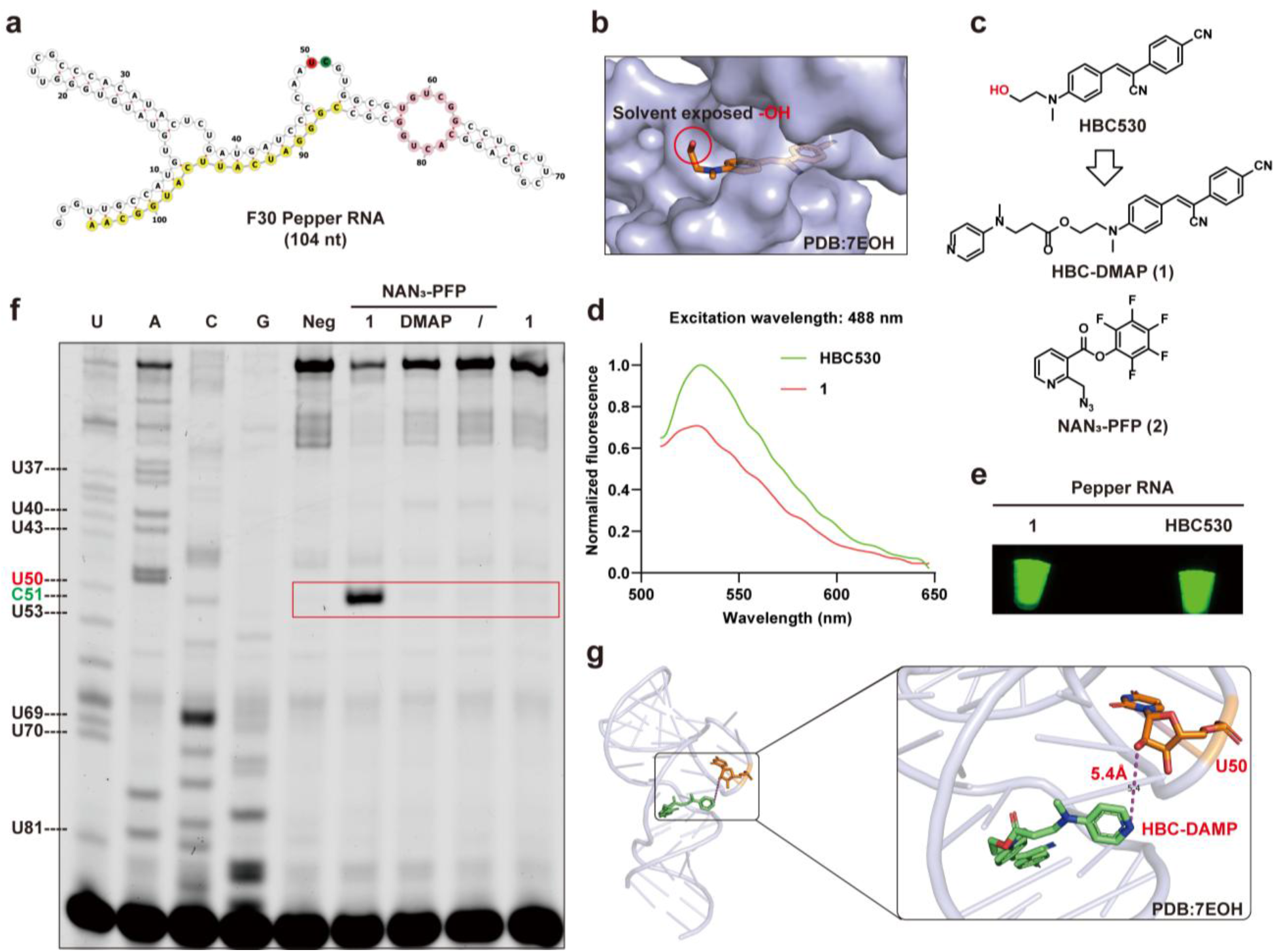
Design and evaluation of the HBC-DMAP catalyst for Pepper RNA modification. (**a**) Secondary structure of F30 Pepper RNA predicted by the RNAFOLD folding algorithm. The binding pocket (pink) for HBC530, primer binding site (yellow) for reverse transcriptase primer extension, the modification site (red) and RT-STOP site (green) are shown. (**b**) Crystal structure of the Pepper RNA with its ligand HBC530 (PDB:7EOH). (**c**) DMAP-tethered Pepper ligand **1** and acyl donor NAN_3_-PFP **2.** (**d**) Normalized fluorescence spectrum of Pepper RNA incubated with HBC530 or **1**. (**e**) Solution of Pepper RNA incubated with HBC530 or **1**, visualized by Typhoon fluorescence imaging. (**f**) Primer extension assay for the identification of modified bases by **1** with **2**. The reaction was conducted with 2 µM Pepper RNA, 10 µM HBC-DMAP or DMAP, 20 mM NAN_3_-PFP in 100 mM MOPS pH 8, 6 mM MgCl_2_, and 100 mM NaCl for 4 h under 37 °C, 8% DMSO. (**g**) Molecular docking study of **1** with Pepper RNA (PDB: 7EOH).

Guided by the reported co-crystal structure of HBC530 bound to the Pepper RNA aptamer, we noted that the solvent-exposed orientation of the hydroxyl group from HBC530 provides an ideal site for DMAP conjugation with minimal disruption to ligand-RNA binding (**Figure 2b**). Accordingly, we synthesized the DMAP-conjugated ligand HBC-DMAP (**1**) and employed NAN_3_-PFP (**2**) as the acyl donor, which in our previous studies demonstrated low background reactivity alongside high catalytic efficiency when paired with DNA-DMAP conjugates (**Figure 2c**).^13^ To assess whether **1** maintains binding affinity for the Pepper RNA, we measured fluorescence upon RNA binding, using HBC530 as a reference. Ligand **1** retained ~70% of the fluorescence intensity of HBC530, consistent with preservation of high-affinity binding (**Figure 2d** and **2e**).

We next examined RNA acylation by incubating F30-scaffolded Pepper RNA (2 μM) with **1** (10 μM) and **2** (20 mM) in MOPS buffer at 37 °C for 4 h. Modification sites were mapped using primer extension, where reverse transcription (RT) terminates at acylated nucleotides. The resulting truncated products were resolved by denaturing polyacrylamide gel electrophoresis (PAGE) and aligned with Sanger sequencing ladders generated from the corresponding ddNTP/dNTP mixtures.^4, 26^ A pronounced RT-stop band was observed only in the presence of both **1** and **2**, whereas no truncation occurred with **2** alone or with untargeted DMAP (**Figure 2f**). Alignment with the sequencing ladders identified the modified site as U50. Notably, U50 lies outside the HBC530 binding pocket (bases highlighted in pink in **Figure 2a**), in a distal loop of the Pepper RNA. Structural rearrangements upon ligand binding bring this loop into close spatial proximity to the ligand-binding site.^27^ Consistent with this model, molecular docking of **1** into the Pepper RNA structure showed occupancy of the HBC530 binding pocket with the DMAP moiety positioned toward the 2′-OH group of U50, supporting a proximity-driven, site-selective modification mechanism (**Figure 2g**).

To further verify that the modification occurs at the 2′-OH of U50, we employed the shortest reported Pepper RNA sequence (49 nt) and corresponding dU variant in which the 2′-OH at position U32 (corresponding to U50 in F30 Pepper RNA) was removed.^27^ Under identical reaction conditions, primer extension analysis revealed a clear RT-stop band at U32 only for the intact Pepper RNA (49 nt), whereas no such stop was observed for the dU-containing RNA, indicating the absence of modification at this position (**Figure S1a**). We further validated these findings by HPLC analysis. A distinct new peak corresponding to the modified product was observed exclusively in the Pepper RNA (49 nt) sample, while no comparable peak was detected in either the DMAP control group or the dU RNA sample (**Figure S1b**). Consistent with these results, MALDI-TOF mass spectrometry showed a mass shift corresponding to the modified RNA only in the presence of the intact U32 residue, whereas no such modified mass peak was detected for the dU RNA sample (**Figure S1c**).

Collectively, these results demonstrate that DMAP-conjugated small molecules, when paired with an efficient acyl donor, can achieve affinity-guided, site-selective chemical modification of target RNAs.

### SAR and kinetics studies on this RNA acylation reaction

We next investigated the structure-activity relationship (SAR) of HBC-DMAP conjugates. Extending the flexible linker between the HBC scaffold and the DMAP catalyst generated compounds **3** and **4**, while modification of the DMAP moiety yielded compound **5** (**Figure S2a**). Fluorescence measurements indicated that all three analogues retained ~50% of HBC530’s emission intensity, consistent with preserved binding affinity toward Pepper RNA (**Figure S2b**). Each compound was then incubated with Pepper RNA in the presence of **2**, and modification sites and efficiencies were assessed by primer extension (**Figure S2c**). All analogues modified the same site and displayed efficiencies comparable to compound **1** (**Figure S2d**). This uniformity likely reflects the preferential reactivity of U50, positioned near the binding pocket, combined with the flexible linkers in these HBC-DMAP analogues, resulting in minimal variation in site selectivity and overall acylation efficiency.

The azide group on acyl donor **2** enables conjugation of cyanine5-dibenzocyclooctyne (Cy5-DBCO) to **2**-modified RNA via click reaction, allowing the kinetics of acylation to be monitored by Cy5 fluorescence. To delineate factors governing reaction efficiency and specificity, we systematically varied the concentrations of ligand **1** and acyl donor **2**, as well as the reaction time. Increasing either component enhanced RNA labeling, indicating that the reaction rate is positively correlated with the availability of both ligand and acyl donor. Notably, **2** exhibited low background reactivity even at concentrations up to 20 mM, highlighting its favorable selectivity profile (**Figure S3a**). In contrast, the concentration of **1** required careful optimization: maximal site-selective modification with minimal nonspecific background was achieved at 10 μM (**Figure S3b**), whereas higher concentrations increased off-target acylation, likely due to the inherent catalytic activity of the DMAP moiety. Time-course studies showed progressive accumulation of RNA modification, reaching a plateau after ~6 h (**Figure S3c**). Prolonged incubation beyond this point did not improve labeling and caused a slight decrease, which we attribute to partial hydrolysis of some installed ester linkages under the reaction conditions.

Following the structure-activity and kinetic analyses, we assessed whether this approach could enable simultaneous multi-site modification on longer RNAs containing tandem Pepper repeats, which are often used to enhance fluorescence output.^25^ An *in vitro* transcribed RNA harboring two Pepper units was treated with ligand **1** or **5** in the presence of acyl donor **2**. Primer extension revealed one prominent modification site, while a second site near the 5′ end was poorly resolved, likely due to RT stalling (**Figure S4a/b**). Importantly, after DBCO-Cy5 click labeling, the two-Pepper RNA displayed nearly twice the Cy5 fluorescence of the single-Pepper RNA, consistent with concurrent modification at multiple sites (**Figure S4c**/**d**).

### Optimization of catalyst-acyl donor pairs for improved RNA modification efficiency under neutral conditions

Building on the proof-of-concept studies, we aimed to further optimize the catalytic system with two primary objectives: (i) to enhance the aqueous solubility of acyl donor **2**, which currently requires 8% DMSO, and (ii) to identify catalyst-acyl donor pairs that operate efficiently under neutral conditions, enabling potential intracellular applications. To this end, we designed, synthesized, and evaluated HBC derivatives incorporating pyridinium oxime (PyOx) or imidazole in combination with structurally diverse acyl donors featuring different acyl and leaving groups (**Figure 3a**).

**Figure 3.**
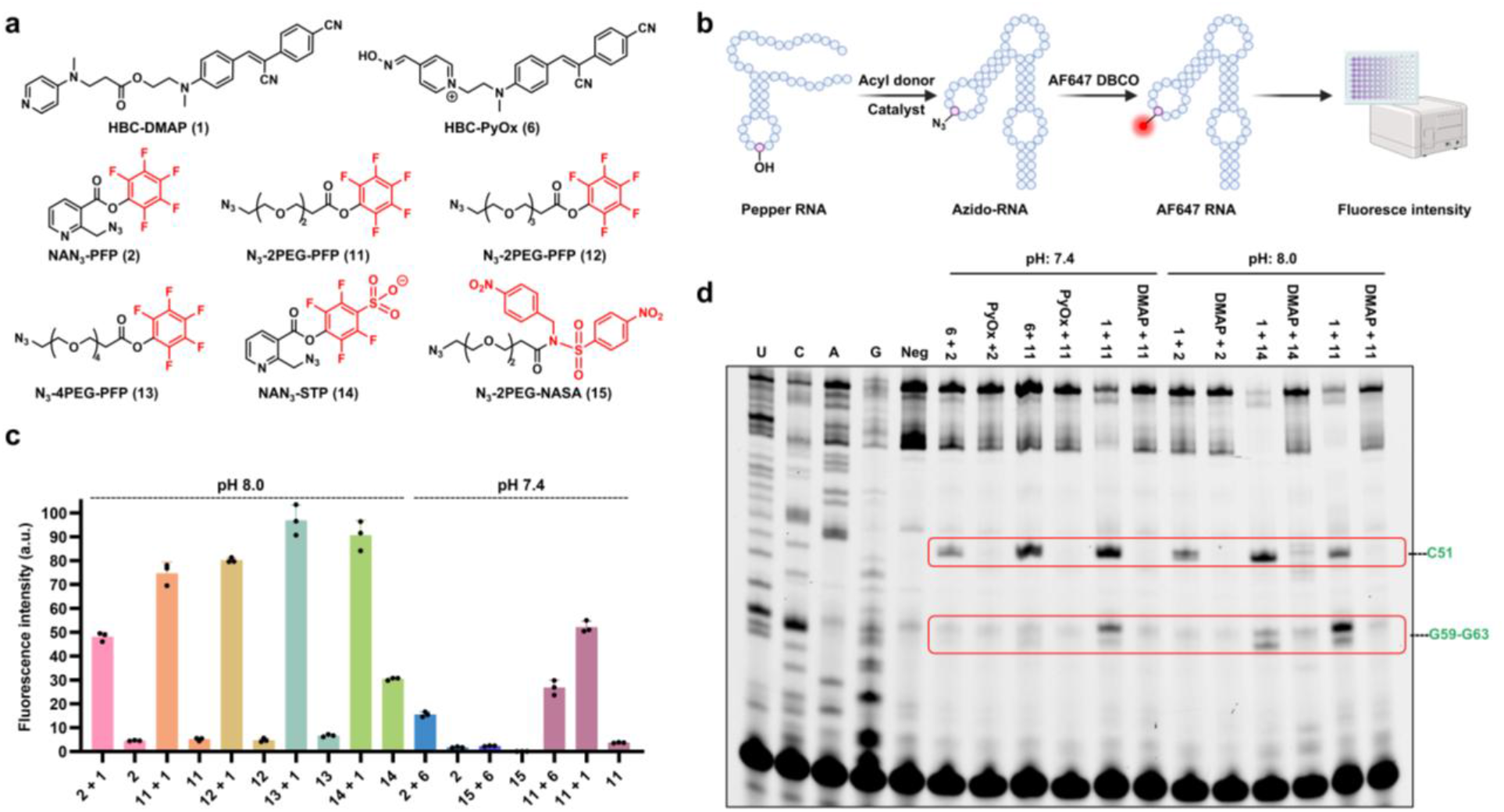
Screening different types of catalyst-acyl donor combinations. (**a**) Structures of Pyridinium oxime- and DMAP-tethered ligands and diverse acyl donors. (**b**) Illustration of AF647 installment onto azido Pepper. (**c**) Fluorescence-based evaluation of reaction performance of various combinations at neutral (pH 7.4) and basic (pH 8.0) conditions. Data are shown as mean values (*n* = 3 independent experiments). (**d**) Primer extension assay for the identification of the preferred catalytic combinations.

The design of PyOx catalysts was inspired by previous work from the Hamachi group, which demonstrated that ligand-tethered PyOx moieties can promote site-selective protein modification under mild conditions.^11^ Notably, unlike DMAP, these catalysts retain activity at neutral pH when paired with *N*-acyl-*N*-alkylsulfonamide (NASA) acyl donors. Guided by this precedent, we synthesized a series of HBC-PyOx conjugates with linkers of varying lengths and systematically evaluated their RNA-modifying activity via primer extension assays (**Figure S5a**). Among the tested candidates, HBC-PyOx **6**, featuring the shortest linker, exhibited the highest reactivity with acyl donor **2**, though still lower than that of **1** at pH 8 (**Figure S5c**). Minimal RNA modification was detected for HBC-PyOx **7**-**9** with longer linkers. Interestingly, fluorescence measurements upon binding Pepper RNA revealed nearly undetectable emission for **6**, likely arising from fluorescence quenching by the positively charged pyridinium moiety rather loss of binding affinity (**Figure S5b**).

We next evaluated imidazole as an alternative acyl transfer catalyst. Owing to its lower p*K*_a_ (~7.0) relative to DMAP (~ 9.7), imidazole was expected to exist in a greater proportion of its nonprotonated, catalytically competent form at neutral pH. Accordingly, HBC-imidazole conjugate **10** was synthesized and examined for RNA acylation at pH 7.4. Primer extension analysis revealed that its modification efficiency was inferior to that of HBC-PyOx **6** under identical conditions (**Figure S6**). Based on these results, **1** and **6** were selected for further investigation.

In parallel, we designed and evaluated a panel of structurally diverse acyl donors (**Figure 3a**). To address the limited aqueous solubility of acyl donor **2**, we synthesized a series of PEGylated PFP esters with increasing linker lengths (compounds **11**-**13**). In addition, a 4-sulfotetrafluorophenyl (STP)-activated ester (NAN_3_-STP **14**) was prepared, motivated by its broad application in protein modification owing to its high intrinsic reactivity and enhanced aqueous solubility.^28^ Guided by Hamachi’s work, we also synthesized a PEG-based NASA ester (compound **15**) to evaluate its compatibility with the HBC-PyOx catalyst.^11^

These acyl donors were systematically evaluated in combination with either **1** or **6** under basic (pH 8.0) or neutral (pH 7.4) conditions. Following modification, the RNA was subjected to click conjugation with Alexa Fluor 647-dibenzocyclooctyne (AF647-DBCO), and AF647 fluorescence intensity was used to assess the modification efficiency for each catalyst-donor pair (**Figure 3b**). Under basic conditions, the azide-PEG acyl donors (**11**-**13**) in combination with HBC-DMAP **1** consistently outperformed the original acyl donor **2**, providing ~1.5-, 1.6-, and 1.9-fold increases in labeling efficiency, respectively, while maintaining comparably low background reactivity. Acyl donor **14** also displayed strong activity (~1.8-fold enhancement); however, this increase was accompanied by substantially elevated background labeling, approximately fivefold higher than that observed for **2**. Under neutral conditions, the pairing of **6** with **11** afforded ~2-fold higher activity relative to **2**, although both systems remained less efficient than **1** under basic conditions. In contrast, acyl donor **15** exhibited minimal activity with **6**, likely due to its limited aqueous solubility. Notably, the combination of **11** with **1** emerged as the most effective system under neutral, DMSO-free conditions, delivering ~1.2-fold higher activity than the **1**-**2** pair even at pH 8.0, albeit reaching only ~80% of its maximal efficiency observed at pH 8.0 for the **1**-**11** pair (**Figure 3c**).

The fluorescence-based trends were further validated by primer extension assays, which showed good overall agreement (**Figure 3d**). Unexpectedly, reactions employing **11** and **1** generated an additional prominent RT-stop band. Alignment with sequencing ladders assigned this stop to nucleotides located directly within the HBC530 binding pocket of the Pepper RNA (**Figure 2a**). This finding offers a mechanistic rationale for the enhanced performance of the PEG-azide acyl donors: improved aqueous solubility increases the effective concentration of the reactive species, while reduced steric demand and greater conformational flexibility relative to **2** facilitate more efficient catalytic acylation within the ligand-binding pocket. Collectively, these results identify the pairing of **1** with the N_3_-PEG-PFP acyl donor as the most effective system examined, delivering consistently high activity under both basic and neutral conditions with low background reactivity.

### Dual colored fluorescent RNA aptamer and FRET

A fundamental limitation of fluorescent RNA aptamers arises from the background emission of their unbound small-molecule ligands. Because these environmentally sensitive fluorogens can become fluorescent in viscous intracellular environments, generating false-positive signals. The development of orthogonal, multicolor RNA aptamer platforms is therefore highly desirable. To address this challenge, we introduce a strategy based on acyl-installed azide handles that enable subsequent click conjugation with diverse fluorophores. This modular approach facilitates the straightforward construction of dual-color RNAs while minimizing background interference through orthogonal fluorescence readouts (**Figure 4a**).

**Figure 4.**
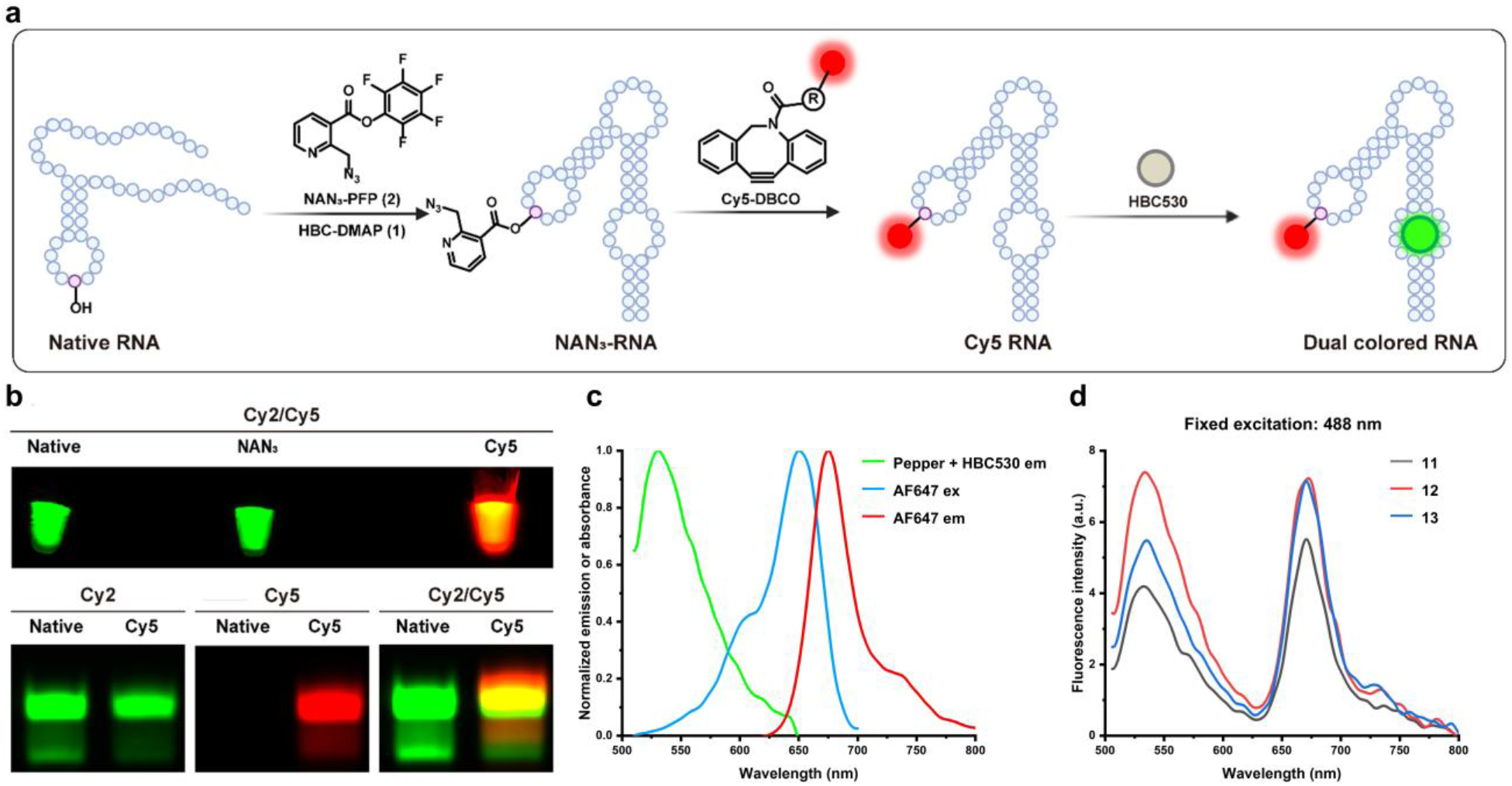
Fluorescence labeling of target RNA and FRET phenomenon. (**a**) Illustration of Cy5 installment onto NAN_3_ modified Pepper to realize dual color. (**b**) Solution of HBC530 incubate with native Pepper, NAN_3_-Pepper and Cy5-labeled Pepper via NAN_3_-PFP **2**, visualized by Typhoon fluorescence imaging (top) and the fluorescence image of 8% denaturing PAGE analysis of the modified RNA (bottom). (**c**) Spectral overlap between the emission of HBC530 and the excitation of AF647. (**d**) The FRET phenomenon of different acyl donor involved modification, AF647 emission was observed under 488 nm excitation, despite 488 nm being outside the excitation range of AF647.

We first performed click conjugation of Cy5-DBCO to acylated Pepper RNA and subsequently imaged the modified RNA following incubation with its cognate ligand HBC530. Fluorescence imaging of RNA-HBC530 complexes, together with denaturing PAGE analysis stained with HBC530, showed that unmodified Pepper RNA and acylated RNA lacking click conjugation exhibited only green fluorescence. In contrast, Cy5-labeled Pepper RNA displayed strong colocalized signals in both the Cy2 and Cy5 channels, appearing yellow in the merged images (**Figure 4a** and **4b**). The modified construct exhibited robust Cy5 emission, accompanied by partial attenuation of the intrinsic green fluorescence (**Figure S7b/c**). We attribute this effect to two factors: (i) the relatively rigid scaffold of acyl donor **2** may partially disrupt proper RNA folding, and (ii) partial Förster resonance energy transfer (FRET), with the Pepper-HBC complex serving as the donor and Cy5 as the acceptor, thereby reducing donor emission (**Figure S7b**). We next examined azide-PEG acyl donors (**11**-**13**), which afford higher modification efficiencies. Following modification, the modified RNA was reacted with AF647-DBCO. Upon excitation at 488 nm, efficient FRET to AF647 was observed across this donor series (**Figure 4c** and **4d**). Notably, AF647 emission arises from energy transfer after formation of the Pepper-HBC complex rather than from direct AF647 excitation, indicating that FRET-based detection effectively suppresses background signals. Moreover, the FRET readout provides a sensitive means to probe RNA structure and RNA-ligand interactions, offering particular utility for systems lacking high-resolution structural information.

### Investigate the general applicability of the strategy on other RNAs

We next sought to extend the optimized catalytic combination to the modification of multiple target RNAs. First, we applied it to another fluorescent RNA aptamer, Clivia, using its ligand NBSI600 (**Figure S8a/b**).^29^ The co-crystal structure showed that the ligand’s hydroxyl group is solvent-exposed, providing a suitable site for tethering the DMAP moiety.^30^ After synthesizing the DMAP-tethered compound NBSI-DMAP (**16**), fluorescence measurements revealed that it exhibited even higher fluorescence upon binding Clivia RNA than NBSI600, indicating strong RNA binding affinity (**Figure S8c**). Ligand **16** was then incubated with Clivia RNA in the presence of acyl donor **11**, and the modified RNA was reacted with Cy5-DBCO. Compared to DMAP alone, **16** produced a 4.5-fold increase in Cy5 fluorescence (**Figure S8d**). Primer extension analysis further confirmed that the modification sites were located within the previously reported NBSI binding pocket, supporting the proximity-driven, affinity-guided RNA modification (**Figure S8e**).

Having established that our RNA modification strategy efficiently targets RNAs lacking G-quadruplex (G4) structures, we next evaluated its applicability to G4-containing RNAs using the Broccoli RNA aptamer as a model system. The initially reported ligand DFHBI was subsequently improved to DFHBI-1T, which exhibits enhanced fluorescence intensity and reduced background.^31^ Guided by the structural evolution from DFHBI to DFHBI-1T, we installed the DMAP moiety at the same position to preserve binding affinity, affording DFHBI-DMAP (**17**) (**Figure 5b**). Fluorescence analysis revealed that **17** elicited even greater emission upon binding Broccoli RNA than DFHBI, consistent with strong affinity (**Figure 5c**). Incubation of **17** with Broccoli RNA in the presence of acyl donor **11** or **2**, followed by primer extension analysis, confirmed selective modification at G60 (**Figure 5d**). Docking studies further revealed that the DMAP warhead of **17** resides in close spatial proximity to the 2′-OH of G60 (~7.4 Å), thereby enabling site-specific acylation (**Figure 5a** and **5e**).

**Figure 5.**
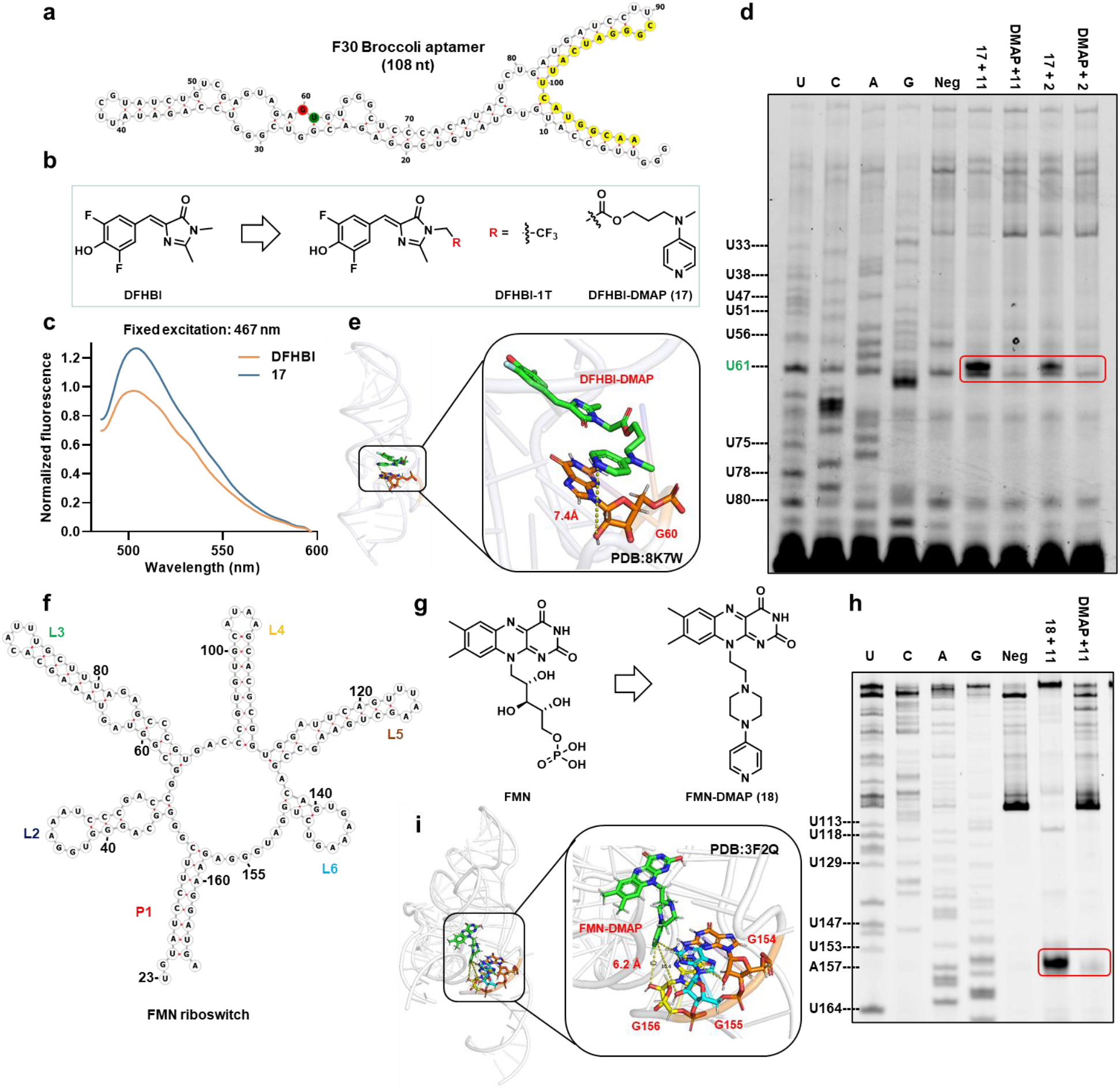
Extension of the affinity-based target RNA modification strategy to G-quadruplex-containing Broccoli RNA and endogenous FMN riboswitch RNA. (**a**) Secondary structure of F30 Broccoli RNA predicted by the RNAFOLD folding algorithm. Primer binding site (yellow) for reverse transcriptase primer extension, the modification site (red) and RT-STOP site (green) are shown. (**b**) Broccoli aptamer ligand (DFHBI and DFHBI-1T) and DMAP-tethered ligand (compound **17**). (**c**) The fluorescence test indicates DFHBI-DMAP **17** maintains good binding affinity with Broccoli RNA. (**d**) Primer extension assay for the identification of modified bases by DFHBI-DMAP **17** with N_3_-2PEG-PFP **11** or NAN_3_-PFP **2.**(**e**) Predicted molecular docking study of DFHBI-DMAP **17** with Broccoli RNA (PDB: 8K7W). (**f**) Secondary structure of the FMN riboswitch from *Bacillus subtilis*, with loop regions annotated in color. (**g**) FMN riboswitch ligand (FMN) and DMAP-tethered ligand (compound **18**). (**h**) Primer extension assay for the identification of modified bases by FMN-DMAP **18** with N_3_-2PEG-PFP **11**. (**i**) Predicted molecular docking study of FMN-DMAP **18** with FMN riboswitch (PDB: 3F2Q).

To further demonstrate the applicability of our strategy to endogenous RNAs, we selected the flavin mononucleotide (FMN) riboswitch RNA-one of the most widely distributed classes of bacterial regulatory RNA elements-as a model target (**Figure 5f**).^32^ FMN serves as its endogenous small-molecule ligand. Previous studies have shown that covalent targeting of FMN RNA can be achieved by installing functional groups on the phosphate side chain of FMN.^22^ Inspired by this strategy, we appended the DMAP-catalytic warhead at the same position to generate FMN-DMAP (**18**) (**Figure 5g**). Incubation of **18** (15 μM) with FMN riboswitch RNA (2 μM) in the presence of acyl donor **11 (**10 mM) at 37 °C for 6 h, followed by primer extension analysis, revealed a pronounced RT-stop signal within the A153-G155 region, consistent with efficient modification at G154-G156 (**Figure 5h**). In contrast, no discernible RT-stop bands were detected in the DMAP control channel.

Previous studies have established that the phosphate side chain of FMN is critical for maintaining high binding affinity.^33^ Accordingly, native FMN is expected to bind FMN riboswitch RNA more tightly than the FMN-DMAP conjugate **18**. On this basis, we performed a competition experiment in which FMN (15 μM) was preincubated with FMN riboswitch RNA for 30 min prior to the addition of **18** and acyl donor **11**. Primer extension analysis showed that the RT-stop signals at G154-G156 were almost completely abolished upon FMN addition, indicating that the observed modification arises from proximity-driven acylation mediated by FMN-DMAP and that FMN-DMAP occupies the same ligand-binding pocket as FMN (**Figure S9**). Molecular docking studies further corroborated this conclusion. Although minor differences in binding conformation were observed, FMN-DMAP occupied the same binding pocket as FMN, with its catalytic moiety positioned in close proximity to the 2′-OH groups of G154-G156, consistent with the experimentally identified modification sites (**Figure 5i**). Taken together, these results demonstrate that the ligand-DMAP (catalyst) and azide-PEG PFP ester (acyl donor) pairs are effective across artificial RNAs lacking or containing G4 structures as well as endogenous RNA targets, underscoring the generality of this affinity-guided, proximity-driven RNA modification strategy.

### Covalent labeling of target RNA by fusing a recognition sequence

Probe-labeled oligonucleotides are widely used in imaging, diagnostics, nanotechnology, and biomedical applications.^34–35^ Conventionally, modified nucleic acids are generated by incorporating functionalized nucleosides during solid-phase synthesis via phosphoramidite chemistry. However, this strategy is generally not applicable to long RNAs, typically exceeding 100 nt in length.^36^ To overcome this limitation, an effective alternative is the use of short, targetable RNA tags fused to the RNA of interest, enabling indirect modification of the target RNA through selective labeling of the tag sequence. One well-established example is the RNA-TAG strategy, which exploits *E*. coli tRNA guanine transglycosylase (TGT) to specifically recognize a 17-nt hairpin RNA sequence (ECYA1) within a fusion construct and catalyze guanosine transglycosylation with functionalized pre-queuosine 1 (PreQ1) analogues.^5^ While highly selective, this approach relies on a specialized enzyme (TGT) that is not broadly present in cells, necessitating exogenous enzyme addition and thereby limiting its applicability.

Here, we demonstrate a distinct enzyme-free tagging strategy in which the Pepper RNA is introduced as a novel RNA tag fused to a target RNA, thereby enabling selective chemical modification of the tag sequence and indirect labeling of the target RNA. To validate this concept, U6 and 7SK small nuclear RNAs (snRNAs) were selected as model targets, as both are widely employed in confocal imaging owing to their well-defined nuclear localization. Pepper-U6 and Pepper-7SK fusion RNAs were generated by IVT.^25^ The Pepper sequence (~50 nt) comprises only a small fraction of the full-length RNA construct, minimizing potential perturbation to the native function of the target RNAs (**Figure S10a/b**). To further assess the extent to which our modification strategy perturbs RNA function, we compared the fluorescence intensities of several modified and native RNAs used in this study. As shown in **Figure S7**, installation of the FRET pair reduced the fluorescence intensity of Cy5-labeled Pepper RNA (**Figure S7a-c**). In contrast, introducing only an azido-bearing group caused minimal changes, with all RNAs retaining comparable fluorescence after modification (**Figure S7d-g**). These results indicate that our strategy enables functionalization of target RNAs with minimal perturbation to their intrinsic properties. Next, each fusion RNA was incubated with **1** in the presence of **11** at pH 8.0, and successful modification was confirmed by primer extension analysis (**Figure 6a** and **S10c**). The weaker modification bands observed for Pepper-U6 likely arise from frequent intrinsic RT-stops induced by Pepper-U6’s secondary or tertiary structures; nevertheless, the identified modification sites are consistent with previous results, supporting efficient and site-selective tagging.

**Figure 6.**
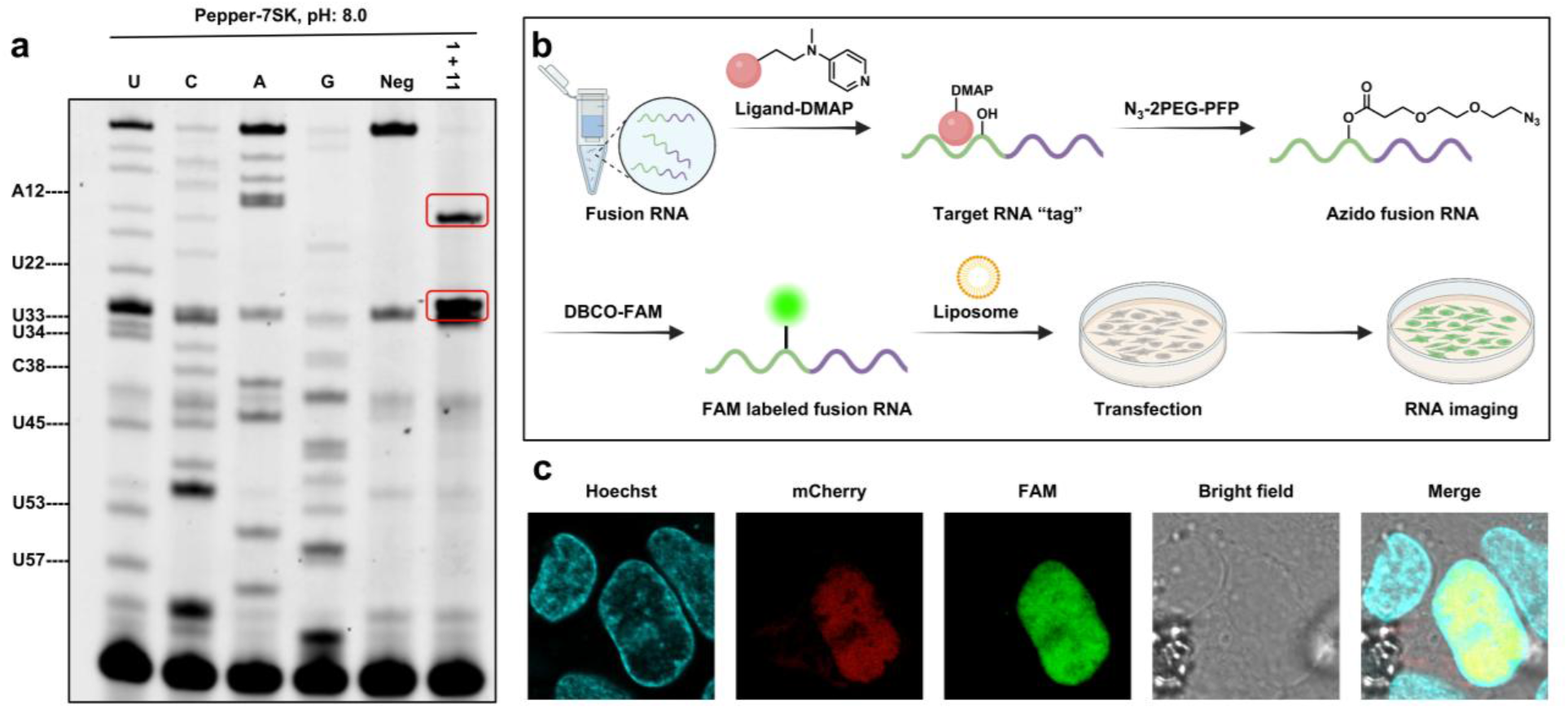
Affinity-guided RNA labeling via fusion of a Pepper RNA tag to the target RNA. (**a**) Primer extension assay for the identification of modified bases of 7SK-Pepper fused RNA. The reaction was conducted with 2 µM Pepper-7SK RNA, 10 µM compound **1**, 10 mM **11** in100mM MOPS pH 8, 6mM MgCl_2_,100mM NaCl for 4 h. (**b**) Schematic illustration showing selective recognition and modification of the Pepper sequence in the fusion RNA by HBC-DMAP, with minimal impact on the nuclear localization of 7SK RNA after modification. (**c**) Images of U2OS cells transfected with FAM-labeled Pepper-7SK RNA and SC35-mCherry plasmid. Blue: Nuclei stained by Hoechst 33342; red: translated mCherry-SC35 protein; green: FAM-labeled RNA.

The modified Pepper-7SK fusion RNA was subsequently evaluated to determine whether the introduced modification affects RNA function. 7SK RNA was chosen as a model target because of its well-defined nuclear localization, providing a robust and reproducible benchmark for evaluating RNA imaging and labeling strategies by confocal microscopy. The resulting azide-containing Pepper-7SK fusion RNA was conjugated to fluorescein (FAM)-DBCO, followed by purification using an RNA cleanup and concentration column to afford FAM-labeled Pepper-7SK RNA. To assess whether the modified Pepper-7SK RNA retained the native nuclear localization of 7SK RNA, the FAM-labeled fusion RNA was transfected into U2OS cells together with an SC35-mCherry plasmid to evaluate nuclear colocalization (**Figure 6b**).^37^ Confocal imaging revealed that the modified Pepper-7SK RNA preserved robust nuclear localization and exhibited clear colocalization with SC35-positive nuclear speckles (**Figure 6c**). In addition, we performed in vitro fluorescence intensity comparisons using a microplate reader, which revealed that the fluorescence intensity of FAM-labeled Pepper-7SK RNA is nearly an order of magnitude higher than that of the same concentration of unlabeled Pepper-7SK RNA in the presence of HBC530 (**Figure S10d**). Consistent with this observation, cellular imaging experiments comparing unmodified Pepper-7SK RNA stained with HBC530 to covalently FAM-labeled Pepper-7SK RNA after transfection showed that, under identical imaging conditions, the unmodified Pepper-7SK RNA produced negligible signal and was nearly undetectable (**Figure S10e**).

Collectively, these results demonstrate that selective chemical modification of a Pepper RNA tag enables efficient and functional labeling of a fused target RNA, validating this strategy as a versatile and enzyme-independent approach for indirect labeling of target RNA.

### Affinity-guided RNA modification facilitates RNA ligand profiling

Given the central roles of RNA in regulating gene expression and genome organization, it is not surprising that RNA dysregulation is implicated in numerous human diseases.^38^ Accordingly, strategies that enable selective RNA targeting hold considerable therapeutic promise. The RNA-targeted modification platform described herein operates through a proximity-driven mechanism, wherein the catalyst-bearing ligand must first engage its cognate RNA with defined affinity. This binding event positions the DMAP-tethered ligand within the RNA binding pocket, thereby promoting selective acylation of proximal 2′-OH groups. Such an affinity-guided mode of action renders this approach particularly well suited for RNA ligand profiling and target engagement studies.

To assess this capability, the Pepper RNA-previously validated and readily amenable to chemical modification-was employed as a model system (**Figure 7a**). HEK293T cells were transfected with a plasmid encoding a circular Pepper-RhoBAST fusion RNA.^39^ Total RNA was subsequently extracted and incubated *in vitro* with the acyl donor **11** and ligand **1** at pH 7.4, a near-physiological condition selected to facilitate potential intracellular translation of the method. Following selective modification of the target RNA, the installed azide groups on the Pepper RNA were conjugated to desthiobiotin-DBCO via click chemistry. The resulting RNA mixture was incubated with streptavidin-coated magnetic beads, subjected to extensive washing, and then released by tris(2-carboxyethyl) phosphine (TCEP)-mediated cleavage of disulfide linkages and blocked by iodoacetamide (IAA).

**Figure 7.**
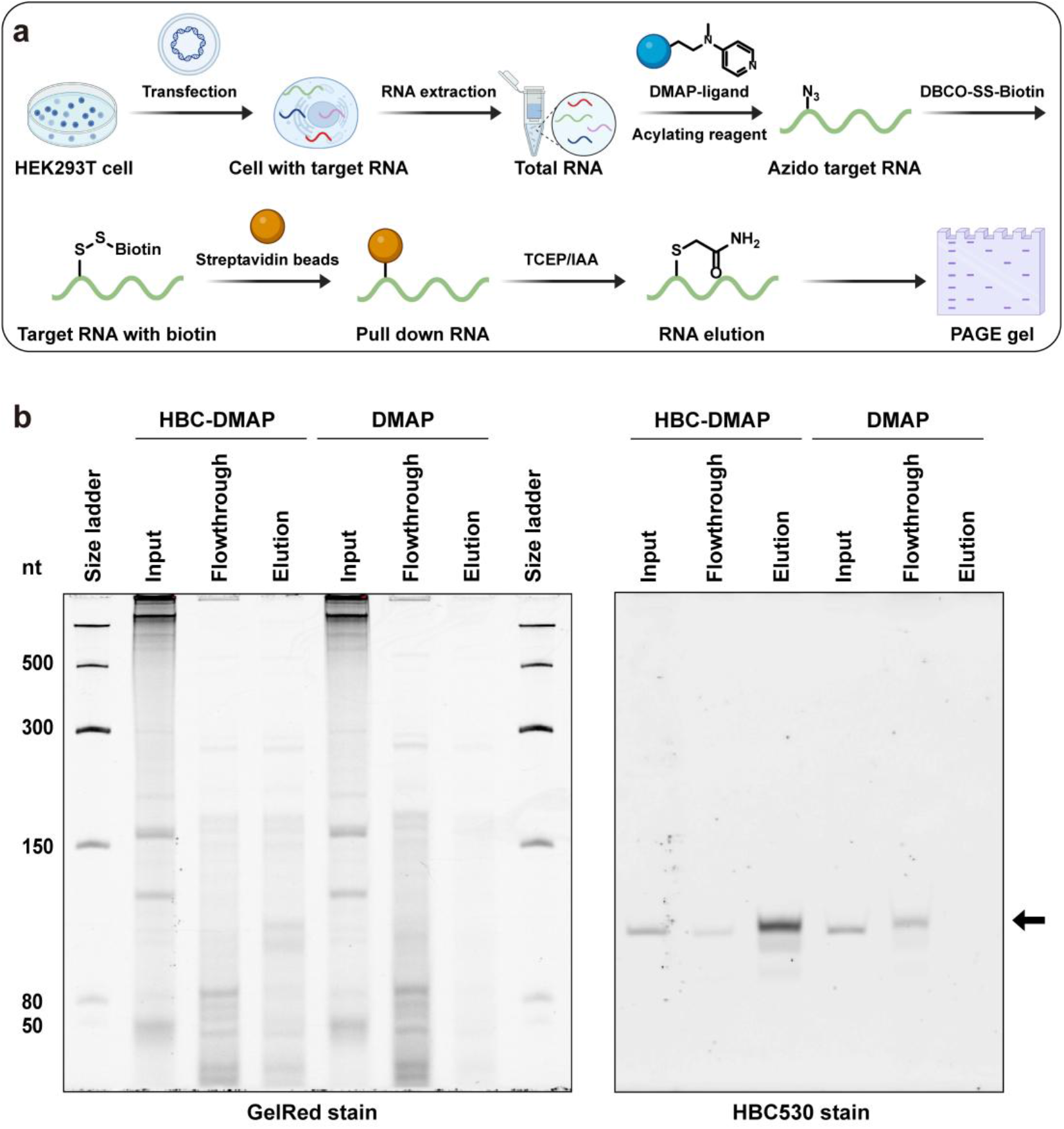
Affinity-based target RNA modification facilitates RNA ligand profiling. (**a**) Schematic presentation of RNA pulldown to enrich target RNA. (**b**) Aliquots (2%) of input and flowthrough and the elution fraction (50%) were separated on a 10% TBE-urea-PAGE gel. Pepper-RhoBAST circular RNA (139 nt) was visualized by stain with HBC530 (right panel). Total RNA was stained with GelRed (left panel).

Target RNA pull-down was assessed by 10% denaturing PAGE, followed by sequential staining with HBC530 and GelRed (**Figure 7b**). Pepper RNA was readily visualized as the same band in the eluate for both HBC530 and GelRed staining, whereas only faint signals were detected in the input and flow-through fractions. Quantitative fluorescence analysis revealed a pull-down efficiency of ~35% of total Pepper RNA, with the HBC-DMAP ligand exhibiting approximately fivefold enrichment relative to DMAP alone. These findings further confirm that the observed enrichment is affinity-driven. Collectively, the data demonstrate that the bifunctional HBC ligand enables substantial and selective enrichment of the target RNA over total RNA in streptavidin-mediated pull-down experiments, highlighting the potential of this approach for RNA ligand profiling in complex cellular settings.

## Conclusions

In this work, we report an affinity-guided strategy for selective RNA modification that leverages the specific recognition between small-molecule ligands and their target RNAs. By appending a catalytic DMAP moiety at a judiciously chosen position on an RNA-binding ligand, site-selective acylation of 2′-OH groups was achieved in the presence of appropriate acyl donors. Successful modification was confirmed through both fluorescence-based assays and primer extension analysis. Systematic screening and optimization of acyl donors identified PEG-PFP ester derivatives, which combine enhanced solubility and reactivity with minimal background labeling. The generality of this approach was further demonstrated across diverse RNA structural contexts, establishing its broad applicability for affinity-guided RNA functionalization. The major differences between prior ligand-directed RNA crosslinking approaches and our modification strategy are twofold: (i) whereas previous methods rely on stoichiometric labeling, our approach operates catalytically, allowing potentially lower ligand concentrations; and (ii) in our method, the ligand and the labeling moiety are decoupled, in contrast to integrated conjugates that are often synthetically challenging and difficult to further derivatize.

Importantly, the labeling occurs at nucleotides proximal to the ligand-binding site, the site-selective modification provides direct information about ligand-RNA interactions. This feature is particularly valuable for RNA-ligand complexes that lack high-resolution structural data. Furthermore, this entirely chemical, enzyme-independent strategy was successfully applied to fusion constructs in which Pepper RNA tags were appended to 7SK target RNA. The modified fusion RNAs preserved the native functional properties of 7SK, enabling accurate intracellular localization. Finally, the combination of HBC-DMAP **1** with the N_3_-2PEG-PFP **11** acyl donor enabled selective enrichment of the Pepper-tagged RNA from total RNA under near-physiological conditions, highlighting the utility of this approach for RNA ligand profiling. Looking forward, this methodology offers a promising platform for direct applications in live-cell RNA targeting and functional studies.

## Acknowledgements

This research was supported by the National University of Singapore and the Ministry of Education, Singapore, under Academic Research Fund Tier 2 MOE-T2EP10223-0002 (R.-Y.Z.), Tier 2 MOE-T2EP30223-0004 (R.-Y.Z.), Tier 2 MOE-T2EP10125-0008 (R.-Y.Z.), and Tier 3 MOET32023-0003. Figures are created with BioRender. U2OS cell line was a gift from Professor Jiangbo Wei and Zhanghui Han. We also acknowledge the facilities, and the scientific and technical assistance of the Chemical, Molecular and Materials Analysis Centre, Department of Chemistry, and Confocal Microscopy Laboratory at Center for Bioimaging Sciences, Department of Biological Sciences, National University of Singapore.

## Notes

### Competing Interest Statement

The authors have declared no competing interest.

