## supporting information for "Small Molecule-Directed RNA Modification via Proximity-Driven Catalysis"

### 1. General information

#### Supplementary Table 1. List of nucleic acids used in this study

DNAs and RNAs were purchased from Integrated DNA Technologies (IDT).

| Sequence name | Sequence (from 5' to 3') | Reference |
|---|---|---|
| DNA template for F30 Pepper RNA | TTGCCATGAATGATCCCGGCGCCAGTGCCTGCCG AAGCAGGCCGACACGCCACGATTGGGGATCATC AGAGTATGTGGGCGAACCCACATACACATGGCA ACCCTATAGTGAGTCGTATTA | 1 |
| DNA template for F30 2x Pepper RNA | TTGCCATGAATGATCCCGGCGCCAGTGCCTCTCC GGCGCCAGTGCCTCTCCGAAGAGAGGCCGACAC GCCACGATTGGGAGAGGCCGACACGCCACGATT GGGGATCATCAGAGTATGTGGGCGAACCCACAT ACACATGGCAACCCTATAGTGAGTCGTATTA | 1 |
| DNA template for F30 Broccoli RNA | TTGCCATGAATGATCCCGAAGGATCATCAGAGT ATGTGGGAGCCCACACTCTACTCGACAGATACG AATATCTGGACCCGACCGTCTCCCACATACACAT GGCAACCCTATAGTGAGTCGTATTA | 2 |
| DNA template for F30 Clivia RNA | TTGCCATGAATGATCCCGAAGGATCATCAGAGT ATGTGGGAGGAAGTGTCCGCCTTTCGGCGTGTTT ACAATCTTCCTCCCACATACACATGGCAACCCTA TAGTGAGTCGTATTA | 3 |
| DNA template for FMN riboswitch RNA | GCGGAATTCTAATACGACTCACTATAGGGCCTTC GGGCCAATGTATCCTTCGGGGCAGGGTGGAAAT CCCGACCGGCGGTAGTAAAGCACATTTGCTTTAG AGCCCGTGACCCGTGTGCATAAGCACGCGGTGG ATTCAGTTTAAGCTGAAGCCGACAGTGAAAGTCT GGATGGGAGAAGGATGATCGATCCGGTTCGCCG GATCCAAATCGGGCTTCGGTCCGGTTC | 4 |
| DNA template for Pepper-7SK RNA | GAAAGGCAGACTGCCACATGCAGCGCCTCATTT GGATGTGTCTGCAGTCTTGGAAGCTTGACTACCC TACGTTCTCCTACAAATGGACCTTGAGAGCTTGT TTGGAGGTTCTAGCAGGGGAGCGCAGCTACTCG TATACCCTTGACCGAAGACCGGTCCTCCTCTATC GGGGATGGTCGTCCTCTTCGACCGAGCGCGCAG CTTCGGGAGGGACGCACATGGAGCGGTGAGGGA GGAAGGGGACACCCGCCTAGCCAGCCAGATCAG CCGAATCAACCCTGGCGATCAATGGGGTGACAG ATGTCGCAGCCAGATCGCCTCACATCCGTCGACC GGTACCTACGGCGCCAGTGCCTGCCGAAGCAGG | 1 |

| | | |
|---|---|---|
| | CCGACACGCCACGATTGGTAGGTACCGCCCTATA GTGAGTCGTATTA | |
| DNA template for Pepper-U6 RNA | CAAAATATGGAACGCTTCACGAATTTGCGTGTCA TCCTTGCGCAGGGGCCATGCTAATCTTCTCTGTA TCGTTCCAATTTTAGTATATGTGCTGCCGAAGCG AGCACGTCGACCGGTACCTACGGCGCCAGTGCC TGCCGAAGCAGGCCGACACGCCACGATTGGTAG GTACCGCCCTATAG TGAGTCGTATTA | 1 |
| Pepper RNA (49 nt) | rGrGrCrGrCrArCrUrGrGrCrGrCrUrGrCrGrCrCrUrUrCr GrGrGrCrGrCrCrArArUrCrGrUrArGrCrGrUrGrUrCrGr GrCrGrCrC | 5 |
| dU Pepper RNA (49 nt) | rGrGrCrGrCrArCrUrGrGrCrGrCrUrGrCrGrCrCrUrUrCr GrGrGrCrGrCrCrArA/ideoxyU/rCrGrUrArGrCrGrUrGr UrCrGrGrCrGrCrC | / |
| F30 Pepper RNA RT primer | /56-FAM/ TTGCCATGAATGATCCCG | / |
| F30 2x Pepper RNA RT primer | /56-FAM/ TTGCCATGAATGATCCCG | / |
| F30 Broccoli RNA RT primer | /56-FAM/ TTGCCATGAATGATCCCG | / |
| F30 Clivia RNA RT primer | /56-FAM/ TTGCCATGAATGATCCCG | / |
| FMN riboswitch RNA RT primer | /56-FAM/ GAACCGGACCGAAGCCCG | 4 |
| Pepper-7SK RNA RT primer | /56-FAM/ AGATCGCCTCACATCCGTCG | / |
| Pepper-U6 RNA RT primer | /56-FAM/ ATGTGCTGCCGAAGCGAG | / |
| Pepper RNA (49 nt) Primer | /56-FAM/ GGCGCCGACACG | / |

The plasmid was purchased directly from Addgene, amplified following the recommended protocol, and purified by FastPure Plasmid Mini Kit (vazyme, catalog: DC201-01) for experimental use.

| **Plasmid name** | **Addgene catalog** |
|---|---|
| pRRL-SRSF2-WT-mCherry | 84020 |
| pAV-U6+27-Tornado-Pepper-RhoBAST | 196343 |

**Supplementary Table 2. List of all reagents and materials**

| Reagents | Source | Catalog number |
|---|---|---|
| **Chemicals** | | |
| MOPS | Sigma-Aldrich | #M1254 |
| $MgCl_2$ | BLDpharm | #BD136984 |
| NaCl | GCE Laboratory Chemicals | #E9440 |
| Urea | Sigma-Aldrich | #U5128 |
| Cy5-DBCO | MedChemExpress | #HY-D1068 |
| FAM-DBCO | LumiProbe | #151F0 |
| AF647-DBCO | LumiProbe | #1G8F0 |
| HEPES | Sigma-Aldrich | #54457 |
| TCEP | BLDpharm | BD155793 |
| Iodoacetamide | BLDpharm | BD134751 |
| DBCO-S-S-PEG3-biotin | MedChemExpress | # HY-140128 |
| **Reagents, Buffers and Enzymes** | | |
| Biotechnology water | 1st BASE | #BUF-1180 |
| SuperScript™ III Reverse Transcriptase | Thermo Scientific | #18080044 |
| RNaseOUT™ Recombinant Ribonuclease Inhibitor | Thermo Scientific | #10777019 |
| dNTP set (100 mM) | Thermo Scientific | #R0181 |
| Dideoxynucleotide triphosphate set | Roche | #03732738001 |
| Tri-color 6x DNA loading dye | 1st BASE | #BIO-1560 |
| Acrylamide solution (40%) | 1st BASE | #HC2040 |
| 10x TBE buffer | Vivantis | #PB1040 |
| Ammonium persulfate | Thermo Scientific | #17874 |
| TEMED | Thermo Scientific | #HC2006 |
| 10x PBS buffer | 1st BASE | #BUF-2040 |
| Dulbecco's Modified Eagle Medium (DMEM) | Gibco | #11995065 |
| Fetal bovine serum (FBS) | Gibco | #A5209402 |
| Trizol™ LS reagent | Thermo Scientific | #10296028 |
| SYBR™ Safe DNA gel stain | Thermo Scientific | #S33102 |
| Opti-MEM™ | Thermo Scientific | #31985070 |
| Lipofectamine™ transfection reagent | Thermo Scientific | #LMRNA001 |
| Hoechst 33342 | Thermo Scientific | #H1399 |
| GelRed | Biotium | #41003 |
| **Commercial kits** | | |
| RNA clean-up and concentrator-5 column | Zymo Research | #R1016 |

| | | |
|---|---|---|
| HiScribe™ T7 Quick High Yield RNA Synthesis Kit | New England BioLabs | #E2050S |
| FastPure Plasmid Mini Kit | Vazyme | #DC201-01 |
| Dynabeads™ Streptavidin Magnetic Beads | Thermo Scientific | #65001 |
| **Softwares** | | |
| MestReNova | Mestrelab | N/A |
| ImageJ | NIH | N/A |
| Origin2021b | OriginLab | N/A |
| BioRender | Biorender.com | N/A |
| PyMol | Schrödinger, Inc | N/A |
| Discovery Studio | BIOVIA | N/A |

#### 2. Supplementary figures

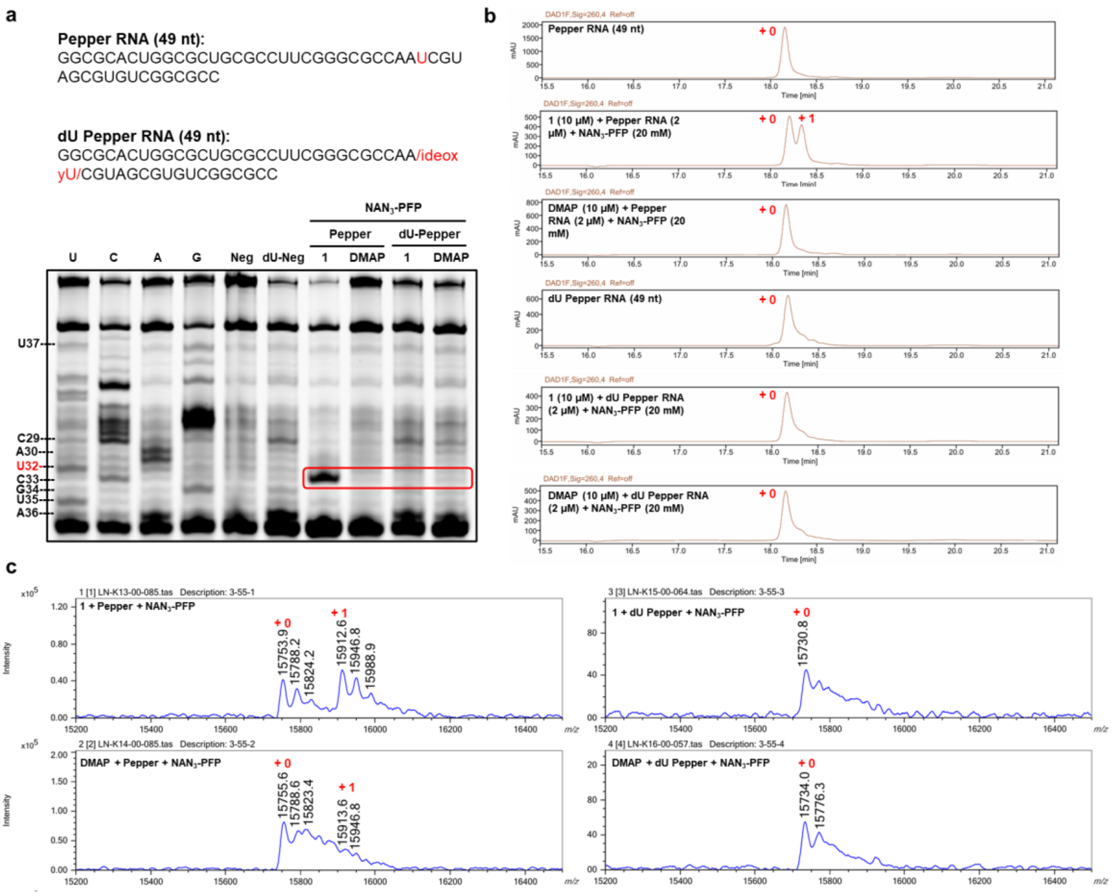


**Figure S1.** Verification of NAN3-PFP **2** mediated modification of target Pepper RNA. (**a**) Primer extension assay used to verify modification sites introduced by acylating reagent NAN3-PFP **2**. Pepper RNA (49 nt) and dU Pepper RNA (49 nt) were used to confirm whether the 2′-OH of U32 (corresponding to U50 in F30 Pepper RNA) is the modification site. (**b**) HPLC analysis of reactions. (**c**) MALDI-TOF mass spectra of Pepper RNA modified by $NAN_3$-PFP **2** in the presence of HBC-DMAP or DMAP. All the reactions were conducted with 2 μM Pepper RNA (49 nt) or dU Pepper RNA (49 nt), 10 μM HBC-DMAP or DMAP, 20 mM $NAN_3$-PFP in 100 mM MOPS pH 8, 6 mM $MgCl_2$, and 100 mM NaCl for 4 h under 37 °C, 8% DMSO.

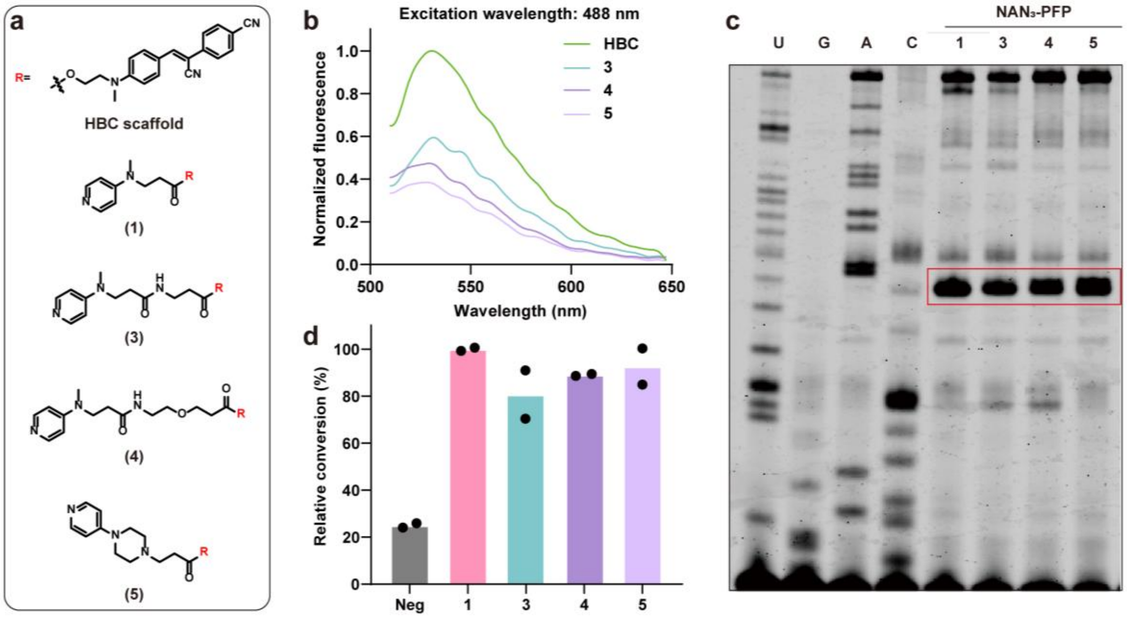


**Figure S2.** Design and evaluation of the analogues of HBC-DMAP **1** with different linkers. (**a**) The structure of DMAP-tethered ligands with different linkers. (**b**) The fluorescence test indicates these ligand-DMAP still maintain good binding affinity with Pepper RNA. (**c**) Primer extension assay for the identification of modified bases by acylating reagents with $NAN_3$-PFP **2**. (**d**) Relative modification efficiency at U50 of Pepper RNA. Data are shown as mean values ($n$ = 2 independent experiments).

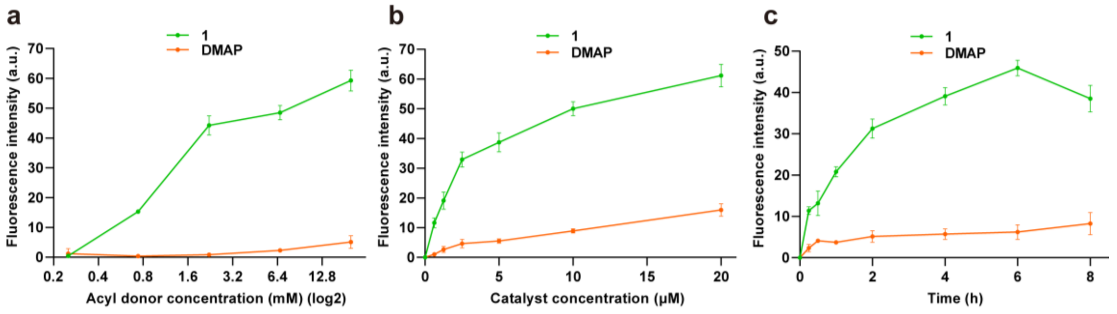


**Figure S3.** Kinetics study of this RNA acylation reaction**.** (**a-c**) The effects of acyl donor concentration (**a**), catalyst concentration (**b**), and incubation time (**c**) on the reaction were investigated by monitoring the Cy5 fluorescence (Excitation: 646 nm, Emission: 664 nm). (**a**) Acyl donor concentrations were varied by threefold serial dilutions starting from 20 mM (five concentrations in total); samples were incubated for 4 h., **1** (10 μM), RNA (2 μM). (**b**) **1** concentrations were varied by twofold serial dilutions starting from 20 μM (six concentrations in total); samples were incubated for 4 h., **2** (20 mM), RNA (2 μM). (**c**) Reactions with varying incubation times: 8 h, 6 h, 4 h, 2 h, 1 h, 30 min, and 15 min, **1** (10 μM), **2** (20 mM), RNA (2 μM). Data represent mean ± s.e.m., *n* = 3 independent experiments (for **a-c**).

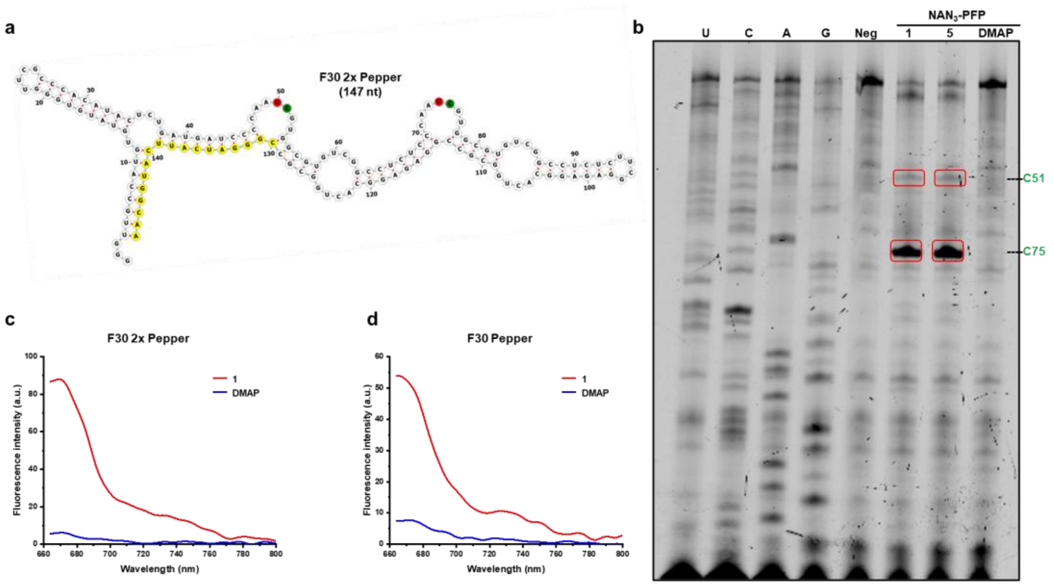


**Figure S4.** Simultaneous modification of RNA containing two Pepper units (F30 2x Pepper)**.** (**a**) Secondary structure of F30 Pepper RNA predicted by the RNAFOLD folding algorithm. The primer binding site (yellow) for reverse transcriptase primer extension, the modification site (red) and RT-STOP site (green) are shown. (**b**) Primer extension assay for the identification of modified bases by **1** with **2** or **5** with **2**. (**c-d**) Cy5 Fluorescence spectrum of 0.2 μM modified F30 2x Pepper or F30 Pepper RNA via NAN3-PFP **2**.

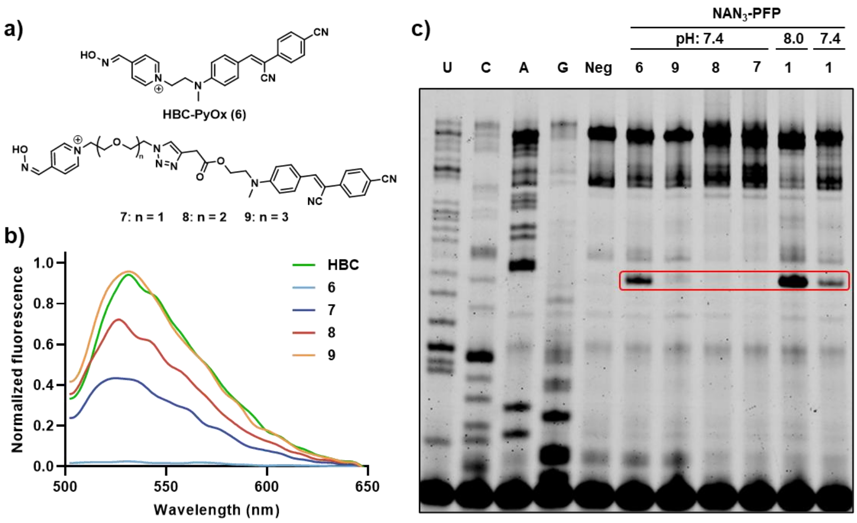

**Figure S5.** Design and evaluation of the analogues of HBC-PyOx (pyridinium oxime) with different linkers. (**a**) The structure of PyOx-tethered ligands with different linkers. (**b**) Fluorescence test to determine the binding affinity between these compounds with Pepper RNA. (**c**) Primer extension assay for the identification of modification efficiency.

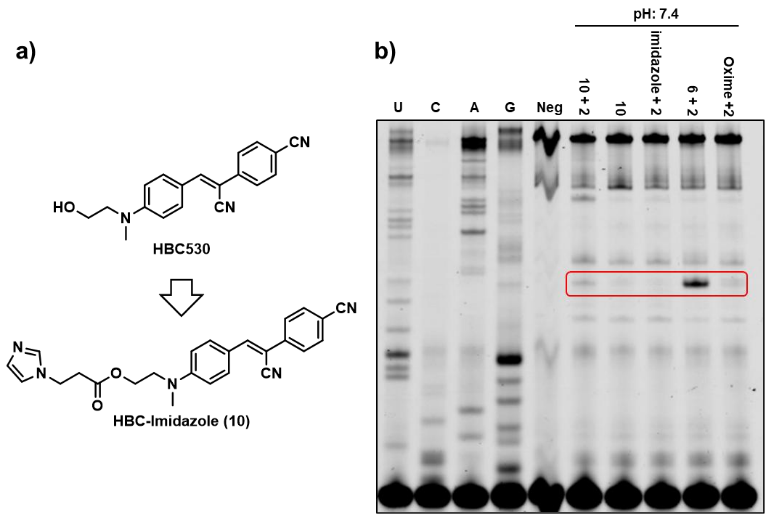


**Figure S6.** Design and evaluation of HBC-Imidazole. (**a**) The structure of Imidazole-tethered ligands (compound **10**). (**b**) Primer extension assay for the identification of modification efficiency under neutral condition.

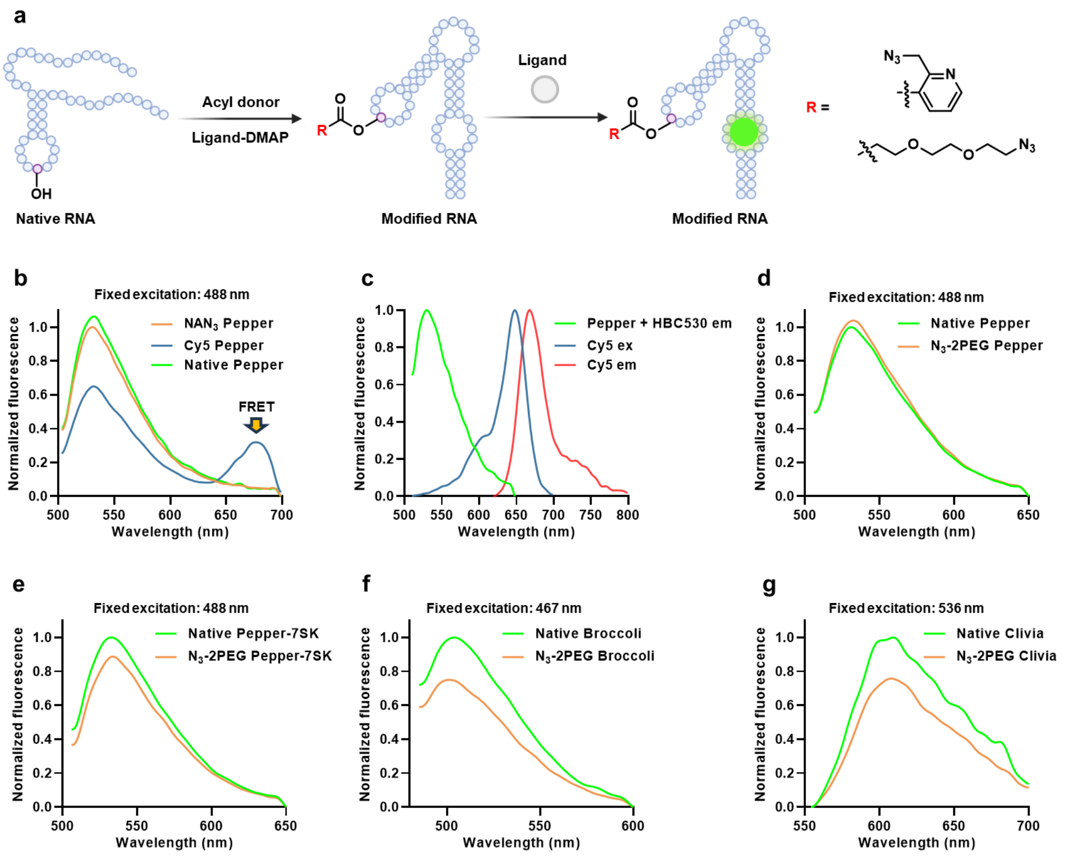

**Figure S7.** Effect of Modification Groups on the Intrinsic Fluorescence of Different RNAs. (**a**) RNA reacts with corresponding ligand-DMAP and acyl donors to get the modified RNA. Then compare the fluorescence intensity between modified RNA and native RNA incubated with their ligand. (**b-c**) NAN$_3$ Pepper still maintains similar fluoresce intensity when compared with native Pepper. But Cy5 Pepper will lose the intensity due to the FRET. (**d-g**) N$_3$-2PEG modified RNA compared the fluorescence intensity with native RNA when incubated with their ligands. (**d**) Pepper (**e**) Pepper-7SK (**f**) Broccoli (**g**) Clivia. All experiments were measured under the following conditions: RNA (0.2 μM) was heated in pH 7.4 HEPES buffer (40 mM HEPES, 5 mM $MgCl_2$, 100 mM KCl) at 65℃ for 5 min, snap-cooled on ice for 10 min, followed by addition of ligand (2 μM) and incubation at room temperature for 0.5 h. The solution was gently mixed, transferred to black 384-well plate, fluorescence was measured by Microplate Reader (Pepper/Pepper-7SK: Ex = 488 nm, Broccoli: Ex = 467 nm, Clivia: Ex = 536 nm).

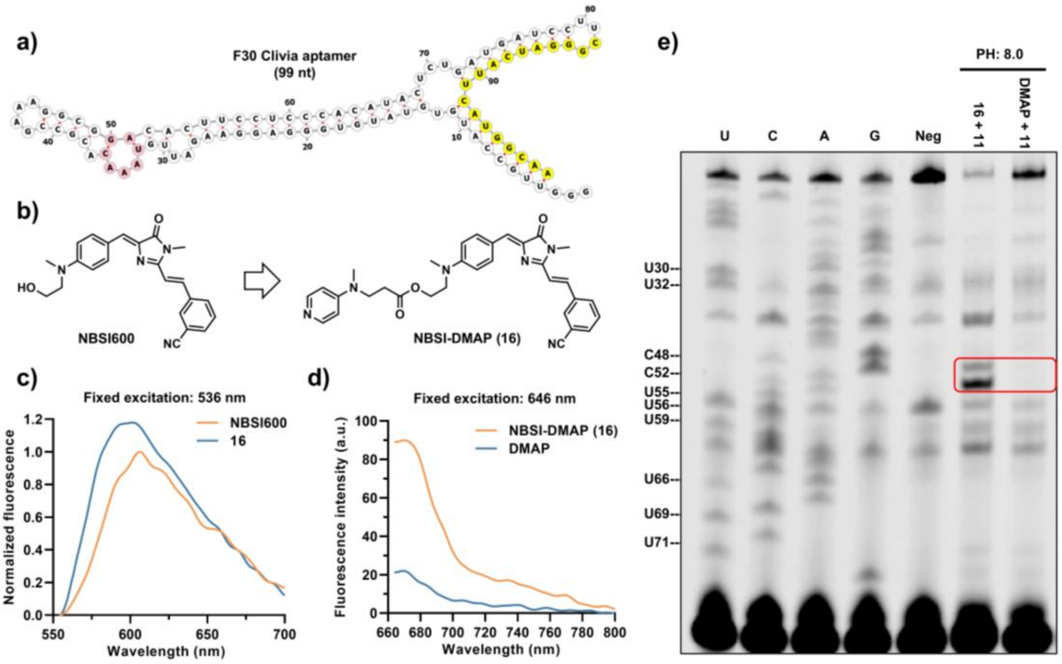


**Figure S8.** Design and evaluation of NBSI-DMAP for Clivia aptamer. (**a**) Secondary structure of F30 Clivia RNA predicted by the RNAFOLD folding algorithm. Primer binding site (yellow) for reverse transcriptase primer extension, the binding pocket (pink) between NBSI with Clivia RNA (PDB: 8HZE) are shown. (**b**) Clivia aptamer ligand (NBSI600) and DMAP-tethered ligand (compound **16**). (**c**) The fluorescence test indicates NBSI-DMAP **16** still maintains good binding affinity with Clivia RNA. (**d**) Cy5 Fluorescence spectrum of 0.2 μM modified Clivia RNA via N$_3$-2PEG-PFP **11**. (**e**) Primer extension assay for the identification of modified sites by NBSI-DMAP **16** with N$_3$-2PEG-PFP **11**.

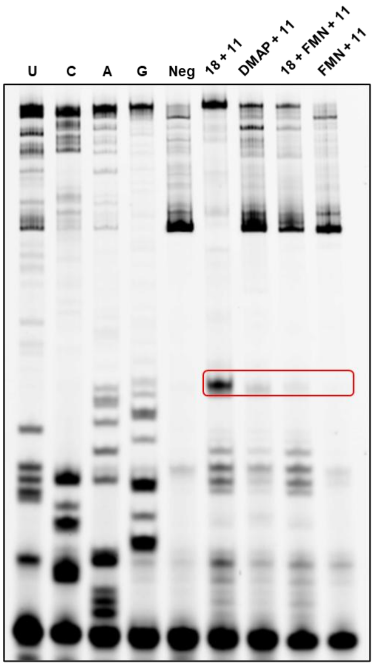


**Figure S9.** FMN competition experiment. FMN (15 μM) was first preincubated with FMN riboswitch RNA (2 μM) in 100mM MOPS pH 8, 6mM $MgCl_2$, 100mM NaCl for 0.5 h at 37 °C. Subsequently, FMN-DMAP (15 μM) and acyl donor **11** (10 mM) were added, and the mixture was incubated at 37 °C for 6 h prior to primer extension analysis.

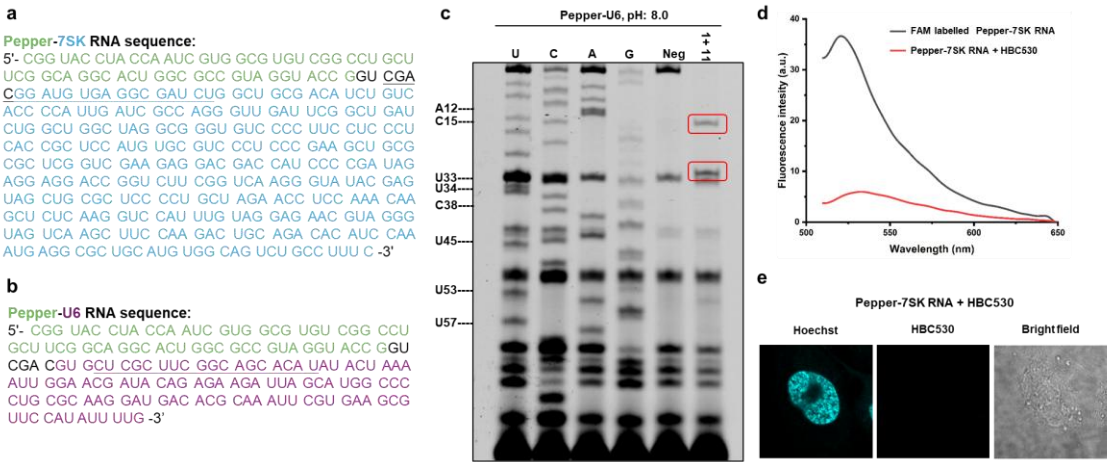


**Figure S10.** Affinity-guided RNA labeling via fusion of a Pepper RNA tag to the target RNA. (**a**) The sequence of Pepper-7SK fusion RNA. Primer binding site (underlined) for reverse transcriptase primer extension, the Pepper sequence (green) and 7SK sequence (blue) are shown. (**b**). The sequence of Pepper-U6 fusion RNA. Primer binding site (underlined) for reverse transcriptase primer extension, the Pepper sequence (green) and U6 sequence (purple) are shown. (**c**) Primer extension assay for identification of modification efficiency. (**d**) Fluorescence spectrum of 0.2 μM Pepper-7SK RNA incubated with 2.0 μM

HBC530 in HEPES buffer (40 mM HEPES, 5 mM $MgCl_2$, 100 mM KCl) for 30 minutes or 0.2 μM FAM labelled Pepper-7SK RNA. Fixed excitation wavelength 488 nm. (**e**) Images of U2OS cells transfected with unmodified Pepper-7SK RNA.

#### Biochemical experiments

**Instrumentation**: Nucleic acid concentration was measured by NanoDrop Lite Plus Spectrophotometers (Thermo Scientific). The A260 value was measured two times to take the average, and the concentration was calculated based on Beer-Lambert law, where the extinction coefficients of nucleic acid were calculated by the IDT Oligoanalyzer tool. MALDI-TOF spectra was acquired by JMS-S3000 SpiralTOF (JEOL), MS Tornado Analysis were used to process the data. Gel images were visualized by Typhoon 9410 fluorescence gel scanner (GE Amersham) or ChemiDoc Imaging System (Bio-Rad). Fluorescence spectra and fluorescence intensity were measured by Varioskan LUX Multimode Microplate Reader (Thermo Scientific) with black, non-binding 384-well plate. Confocal laser scanning microscopy imaging was performed on a FLUOVIEW FV3000 (Olympus).

**In vitro transcription of target RNA**: F30 Pepper RNA, F30 Broccoli RNA, F30 Clivia RNA, FMN riboswitch RNA, Pepper-7SK fusion RNA and Pepper-U6 fusion RNA were synthesized by *in vitro* transcription using HiScribe$^{TM}$ T7 Quick High Yield RNA Synthesis Kit, following the manufacturer's protocol. The reaction was incubated at 37℃ for 16 h. Subsequently, native PAGE of appropriate concentration was used according to the RNA length and run at 200 V for separation. Gel fragment containing desired RNA was excised and extracted in elution buffer (500 mM $NH_4OAc$ and 1 M EDTA pH 8.0) at 4℃ overnight. The solution was then filtered to remove the gel and purified with RNA clean and concentrator-5, following the manufacturer's protocol. RNA concentration was measured by Nanodrop as described above.

**Preparation of acylating reagents:** Acylating reagents are sensitive to moisture and stored in neat form at -20℃ after synthesis. To maintain the quality of acylating reagents, stock solutions in DMSO ($NAN_3$-PFP **2**, $N_3$-2PEG-NASA **15**) or water were freshly prepared before adding to reaction solution.

**Fluorescence spectrum of ligand and target fluorescent RNA:** Fluorescence spectra of the ligand-DMAP catalyst with the target RNA, or the ligand with the modified target RNA, were measured under the following conditions: RNA (0.2 μM) was heated in pH 7.4 HEPES buffer (40 mM HEPES, 5 mM $MgCl_2$, 100 mM KCl) at 65℃ for 5 min, snap-cooled on ice for 10 min, followed by addition of ligand-DMAP catalyst (2 μM) or ligand (2 μM) and incubation at room temperature for 0.5 h. The solution was gently mixed, transferred to black 384-well plate, fluorescence was measured by Microplate Reader (F30 Pepper or Pepper-7SK: Ex = 488 nm, F30 Broccoli: Ex = 467 nm, F30 Clivia: Ex = 536 nm, FAM: Ex = 488 nm, Cy5: Ex = 646 nm, Em: 671 nm, AF647: Ex = 655 nm, Em: 680 nm).

**Target RNA acylation by ligand-DMAP catalyst:** Except for the kinetics studies and FMN riboswitch RNA modification, for which the reaction conditions are described in detail in the figure legends or main-text, all other reactions were conducted under the following conditions. RNA (2 μM) was heated in pH 8.0 MOPS buffer (100 mM MOPS, 6 mM $MgCl_2$, 100 mM KCl) or pH 7.4 HEPES buffer (40 mM HEPES, 5 mM $MgCl_2$, 100 mM KCl), to 65℃ for 5 min and immediately snap cooling on ice for at least 10 min. To the annealed solutions were added ligand-DMAP (10 μM) catalyst and incubate under room temperature for 10 min. Freshly prepared stocks of acylating reagents in DMSO/water were added to the reaction mixture to a final concentration of 20 mM (10mM for Pepper-7SK fusion RNA). The reactions were incubated for 4 h at 37℃, and subsequently purified with RNA clean and concentrator-5, following the manufacturer's protocol. For investigation of reactivity with different ligand-DMAP conjugates, parallel reactions of the RNA in presence of DMAP were measured. The RNA was dissolved in nuclease-free water and stored at -80℃ for further analysis. HPLC analysis was performed using a C18 reverse-phase column (Phenomenex Kinetex EVO C18, 10 × 250 mm) at 55 °C with a flow rate of 0.8 mL $min^{-1}$. The gradient was 0-8 min, 0-50% B; 8-10 min, 50-70% B; 10-35 min, 70-80% B. Solvent A was 100 mM TEAA (pH 7.2), and solvent B was methanol. Detection was at 260 nm.

**MALDI-TOF analysis:** The spectra were recorded in the linear negative mode. The matrix includes 8 parts of 3-HPA solution (50 mg/mL 3-HPA in $H_2O$ and acetonitrile with 1:1 ratio) and 1 part of 50 mg/mL ammonium citrate dibasic aqueous solution. 1 μL of matrix and 1 μL of nucleic acid sample were added into the plate, dried under air flow for 10 min and inserted to MALDI TOF spectrometer for data acquisition. The conversion of acylation reaction was quantified based on the relative areas under the peak.

**Primer extension assay to identify modification site in target RNA:** Except for the FMN riboswitch RNA modification, for which the reaction conditions are same as the previous report,[4] all other reactions were conducted under the following conditions. 4 pmol RNA was mixed with 6 pmol RT Primer and 0.5 μL dNTP mix (10 mM each, for sequencing lane, ddNTP:dNTP = 8:1), and incubated for 5 min at 65℃, then immediately chilled on ice for 2 min. Then 2 μL 5x First-Strand Buffer, 1 μL 0.1 M DTT, 0.5 μL RNaseOUT and 0.25 μL Super Script III (200 U/ μL) were added to the final volume of 10 μL. For analysis of F30 Broccoli RNA, potassium ion component in First-Strand buffer was replaced with lithium ion to minimize the influence of G4 structure. The reaction was incubated with the following program: 25℃ for 10 min, 52℃ for 50 min, and 55℃ for 50 min. After the reaction, 7.25 mg urea and 2 μL 6x DNA loading dye (0.4% Orange G, 0.03% Bromophenol blue, 0.03% Xylene cyanol PF) was added and the mixture was denatured at 95℃ for 3 min. cDNA were analyzed by denaturing appropriate concentration of polyacrylamide gel 1x TBE buffer, constant voltage of 200 V. The cDNA gel was visualized by Typhoon scanner with Cy2 channel.

**Fluorescence labeling of azido target RNA:** To a solution of azido RNA (1 μM) in 1x PBS buffer was added fluorophore-DBCO (100 μM) with the final volume of 50 μL. The solution was incubated at 37℃ for 3 h. The RNA was then purified with RNA clean and concentrator-5, following the manufacturer's protocol. The fluorescence spectrum of RNA was recorded at 0.2 μM concentration in 1x HEPES buffer (40 mM HEPES, 5 mM $MgCl_2$, 100 mM KCl). To measure the fluorescence intensity, the solution was gently mixed, transferred to black 384-well plate, fluorescence was measured by

Microplate Reader (Cy5: Ex/Em = 646/671 nm, AF647: Ex/Em = 655/680 nm). The identity of Cy5 labelled dual colour F30 Pepper RNA was analysed by denaturing 8% polyacrylamide gel in 1x TBE buffer, 200 V, ~1 h. The denaturing PAGE was washed by 1x PBS buffer for one time to dilute the urea then stained by 10 uM HBC530 in 1x HEPES buffer (40 mM HEPES, 5 mM $MgCl_2$, 100 mM KCl). The RNA gel was visualized by Typhoon scanner with Cy5 and Cy2 channel.

**Confocal microscopy of FAM-labelled Pepper-7SK in living cells:** U2OS cells were seeded onto μ-slide 8-well-chambered coverslip (ibidi) at a density of 1.0 x $10^4$ cells and incubated overnight with 200 μL of DMEM supplemented with 10% FBS overnight at 37℃ and 5% $CO_2$ overnight. Transfection of pRRL-SRSF2-WT-mCherry plasmid was performed using Lipofectamine3000 according to the manufacturer's instructions. After 24 h, the medium was removed, cells were transfected with 150 ng FAM-labelled Pepper-7SK RNA by Lipofectamine 3000. After 16 h, the medium was removed, cells were washed 3 times with PBS and stained with 10 μg/mL Hoechst 33342 for 20 min at room temperature. After that, cells were washed 3 times with PBS, replaced with fresh culture medium and imaged with confocal microscope. For unmodified Pepper-7SK transfection, cells were transfected with 150 ng Pepper-7SK RNA by Lipofectamine 3000. After 16 h, the medium was removed, cells were washed 3 times with PBS and stained with 10 μg/mL Hoechst 33342 and 2 μM HBC530 in DMEM for 20 min at room temperature. After that, cells were imaged with confocal microscope equipped with 60x oil lens. Excitation wavelengths were set at 405 nm for Hoechst 33342, 488 nm for FAM-labelled Pepper-7SK or HBC530 and 561 nm for SRSF2-mCherry. Imaging parameters were kept constant throughout the experiment.

**Molecular Docking Study:** Molecular docking studies were performed using BIOVIA Discovery Studio 2019 (Dassault Systèmes BIOVIA, San Diego, CA, USA). The co-crystal structures of Pepper (PDB: 7EOH), Broccoli (PDB: 8K7W) and FMN riboswitch (PDB: 3F2Q) were selected and processed by the RNA Preparation Wizard including water deletion and addition of missing hydrogen atoms. The RNA structure was performing energy minimization using the CHARMm force field. The binding site was defined according to the ligand position, and a region with a radius of 15 Å was used to define the docking pocket. Docking simulations were carried out using the LibDock module with default parameters. The generated docking poses were ranked based on the LibDock score, and the top-scoring complexes were visually inspected to select the most reasonable binding mode.

**Streptavidin pull-down experiments:** Total RNA was isolated from HEK293T cells after 60 hours of transfection with pAV-U6+27-Tornado-Pepper-RhoBAST using Trizol Reagent according to the manufacturer's instructions, dissolved in 40 mM HEPES, 100 mM KCl, 5 mM $MgCl_2$, pH 7.4, heated to 90℃ for 2 min and subsequently transferred to room temperature for 10 min to allow refolding. 40 μg of total RNA were incubated with 10 μM HBC-DMAP **1** or DMAP for 10 min. To the solution was added 2 μL stock of 250 mM $N_3$-2PEG-PFP **11** in water to the final volume of 50 μL. The solution was incubated at 37℃ for 4 h. After that, RNA was purified by ethanol precipitation by adding 0.1x (v:v) of 3 M NaOAc (pH 5.2) and 3.75x (v:v) of absolute EtOH. The mixture was incubated at -80℃ overnight, and the pellet was obtained by centrifuging (21,000 RCF) for 1 h at 4℃. The supernatant was removed, and the pellet was washed 2 times with 75% EtOH and centrifuged (21,000 RCF) for 10 min at 4℃. The supernatant was removed, and the pellet was dried in air for 0.5 h. The RNA was dissolved in nuclease-free water and stored at -80℃ for further analysis. The concentration was determined by Nanodrop as described above. For biotinylation, 20 μL of 2.5 mg/ml DBCO-S-S-$PEG_3$-biotin was added and incubated overnight at room temperature in a total volume of 60 μL with gentle agitation. RNA was purified by ethanol precipitation as above, dissolved in 100 μL of water. Then, the purified RNA was added to100 μL of Streptavidin Magnetic Beads (slurry (150 μL volume of beads) equilibrated in 2× B&W buffer (10 mM Tris-HCl, pH 7.5, 1 mM EDTA, 2 M NaCl) according to the manufacturer's instructions and incubated at room temperature for 2 h with gentle agitation followed by two washes with 1× B&W buffer and RNase-free water. Elution was carried out by incubation of the beads by incubation with 200 μL

Releasing Buffer (200 mM TCEP and 600 mM $K_2CO_3$) at 37℃ for 30 min. An equivalent of 200 μL iodoacetamide (400 mM) was added to the sample, which was incubated at 37℃ for an additional 30 min. Collect the elution and washed the beads by 100 μL RNase-free water. The flowthrough and elution solution were purified with RNA clean and concentrator-5, following the manufacturer's protocol. Input, flowthrough (2% each) and eluate (50%) were separated on 8% TBE-urea polyacrylamide gel. The denaturing PAGE was washed by 1x PBS buffer for one time to dilute the urea then stained by 10 μM HBC530 in 1x HEPES buffer (40 mM HEPES, 5 mM $MgCl_2$, 100 mM KCl). The RNA gel was visualized by Typhoon scanner with Cy2 channel. Subsequently, the gel was stained with GelRed and imaged under UV light by ChemiDoc Imaging System (Bio-Rad).

#### 3. Chemical synthesis of organic molecules

**General synthesis procedures:**

Chemicals were purchased from commercial suppliers and used without further purification unless stated otherwise. All reagents were weighed and handled in air, while anhydrous solvents were handled under $N_2$ atmosphere. Analytical TLC was performed on ready-to-use plates with silica gel 60 (Merck, F254), with ultraviolet light or KMnO4 as means of visualization. Flash column chromatography was performed over Fisher Scientific silica gel (grade 60, 230-400 mesh). Preparative TLC was performed on silica gel Xinnuo HSGF254 preparative TLC plates. Preparative reversed-phase HPLC was conducted with Agilent 1200 DAD HPLC system using Phenomenex C-18 column of 250×10 mm at a flow rate of 3 mL/min with the detector wavelength of 210 and 254 nm. Fraction containing purified compounds from HPLC were evaporated to remove acetonitrile, then the remaining aqueous solutions were freezed at -80℃, and lyophilized in Labconco™ FreeZone™ 2.5 L Freeze Dryer. NMR spectra were acquired on Bruker 500 MHz spectrometer (500 MHz for $^1$H NMR 126 MHz for $^{13}$C NMR) or Bruker 600 MHz spectrometer (600 MHz for $^1$H NMR 150 MHz for $^{13}$C NMR) spectrometer and chemical shift reported in parts per million (δ) relative to internal standard TMS (0 ppm). Calibration was performed with reference to the deuterium NMR solvent (CDCl3: 7.26 [chloroform], CD3OD: 3.31 [methanol], DMSO-d6: 2.50 [dimethyl sulphoxide]) and carbon NMR solvent (CDCl3: 77.00 [chloroform], CD3OD: 49.00 [methanol], DMSO-d6: 39.52 [dimethyl sulphoxide]). The relative number of protons were integrated, and the coupling constants were unified as Hertz (Hz). Multiplicity (s, singlet; d, doublet; t, triplet; q, quartet; p, pentet; m, multiplet) were reported. Mass spectra were measured on a LCQ Fleet LCMS (Thermo Scientific). High-resolution mass spectrometric data was obtained using 6546 LC-QTOF (Agilent).

**Synthesis of HBC-DMAP (1):**

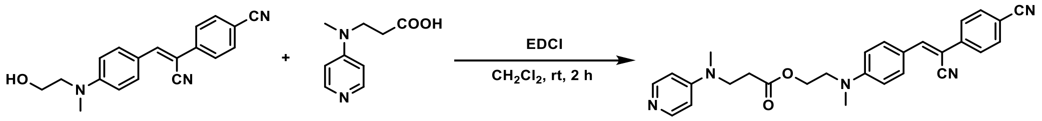


HBC530 and intermediate 1a were prepared according to previous work[1, 6].

To a solution of the **HBC530** (1.0 equiv.), **1a** (1 equiv.) in $CH_2Cl_2$ was added EDCI (1.5 equiv.). The solution was stirred at rt for 2 h, then the solution was then evaporated to remove solvent. The crude mixture was then purified by preparative thin-layer chromatography (DCM:MeOH = 50:1, 1% TEA) to afford the desired product, **1** (**HBC-DMAP**), as a yellow solid (55%).

$^1$H NMR (500 MHz, DMSO) δ 8.20 (d, $J$ = 7.3 Hz, 2H), 8.01 (s, 1H), 7.94 – 7.89 (m, 4H), 7.88 – 7.84 (m, 2H), 7.10 – 6.90 (m, 2H), 6.88 (d, $J$ = 9.2 Hz, 2H), 4.25 (t, $J$ = 5.5 Hz, 2H), 3.74 (m, 4H), 3.08 (s, 3H), 3.03 (s, 3H), 2.60 (t, $J$ = 7.1 Hz, 2H).

$^{13}$C NMR (126 MHz, DMSO) δ 171.28, 158.74, 158.47, 157.04, 151.80, 145.85, 139.97, 133.42, 132.35, 126.01, 121.08, 119.21, 119.18, 112.16, 110.31, 107.67, 101.00, 62.06, 50.22, 47.59, 38.75, 38.47, 31.29.

ESI-MS $[M+H]^+$ calcd for $C_{28}H_{28}N_5O_2$: 466.2238, found: 466.2245.

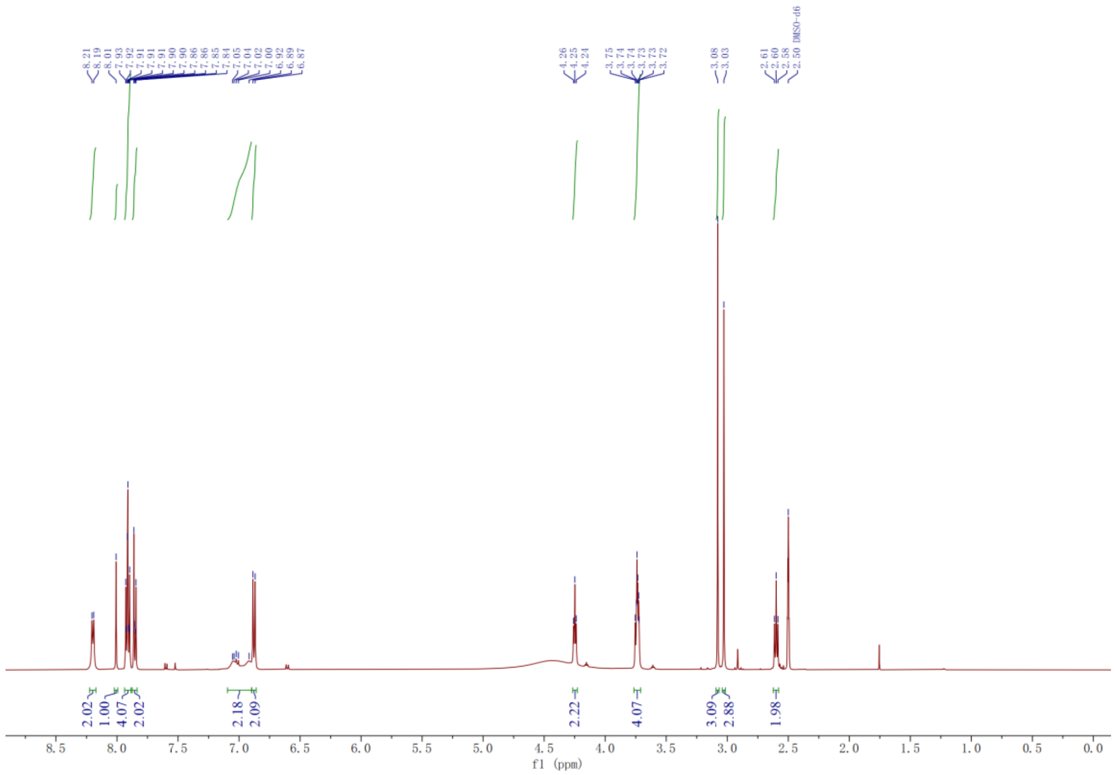

DMSO-d6
f1 (ppm)


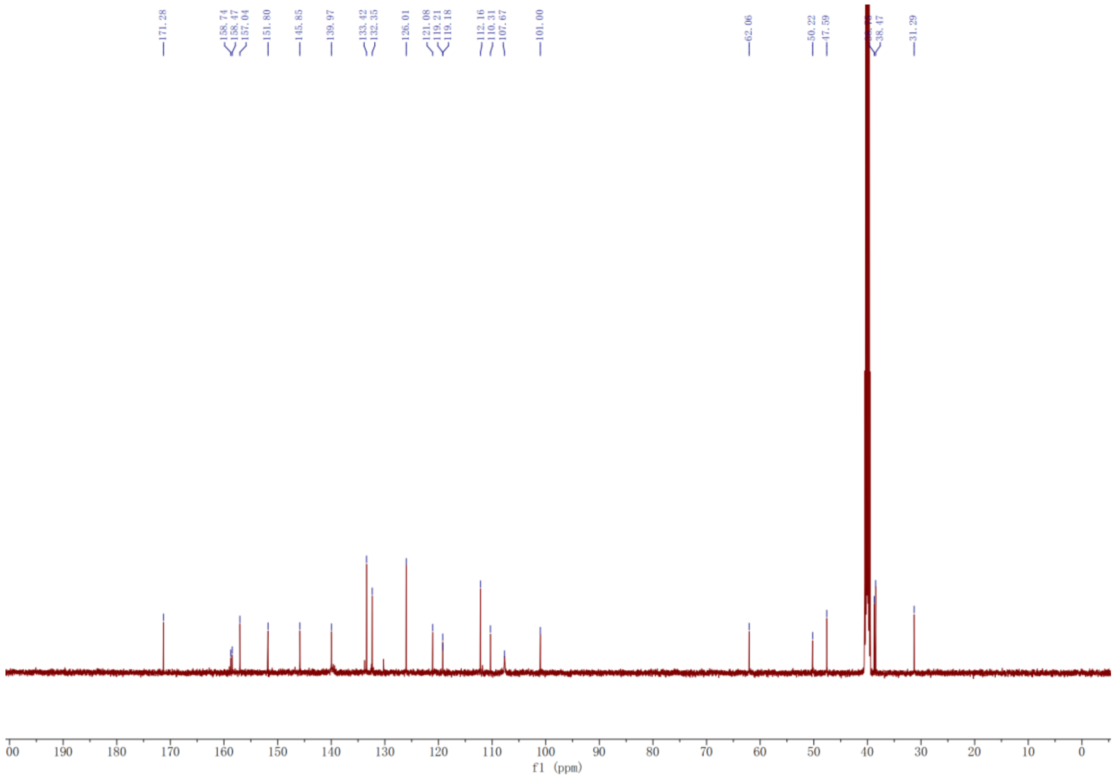

f1 (ppm)

**Synthesis of NAN$_3$-PFP (2):**

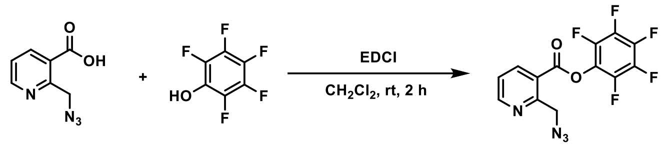


2a 2b 2 (NAN$_3$-PFP)

**2** were prepared according to previous work[7]. Light yellow oil (70%).

$^1$H NMR (500 MHz, $CDCl_3$) δ 8.90 (dd, $J$ = 4.8, 1.8 Hz, 1H), 8.54 (dd, $J$ = 8.0, 1.8 Hz, 1H), 7.50 (dd, $J$ = 7.9, 4.8 Hz, 1H), 4.90 (s, 2H).

$^{13}$C NMR (126 MHz, $CDCl_3$) δ 161.33, 158.19, 153.84, 139.78, 123.18, 121.89, 54.13.

$^{19}$F NMR (471 MHz, $CDCl_3$) δ -152.21 – -152.26 (m), -156.71 (t, $J$ = 21.7 Hz), -161.58 (dd, $J$ = 21.8, 17.4 Hz).

ESI-MS $[M+H]^+$ calcd for $C_{13}H_6F_5N_4O_2$: 345.0405, found: 345.0410.

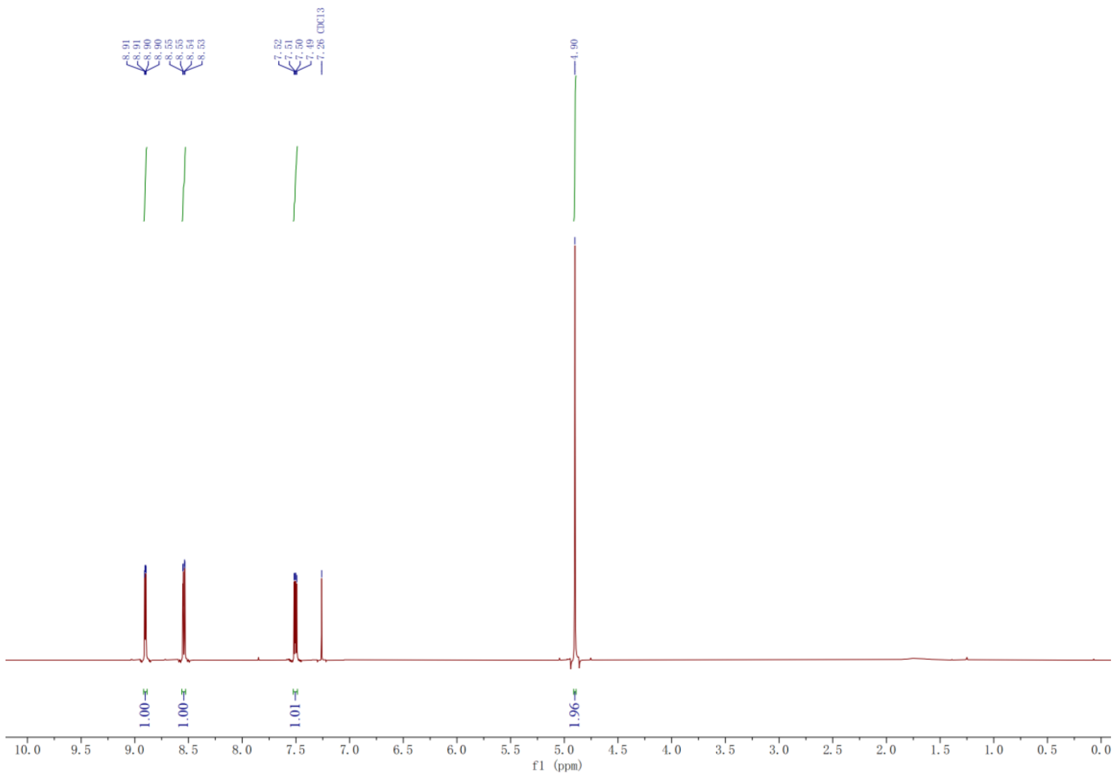

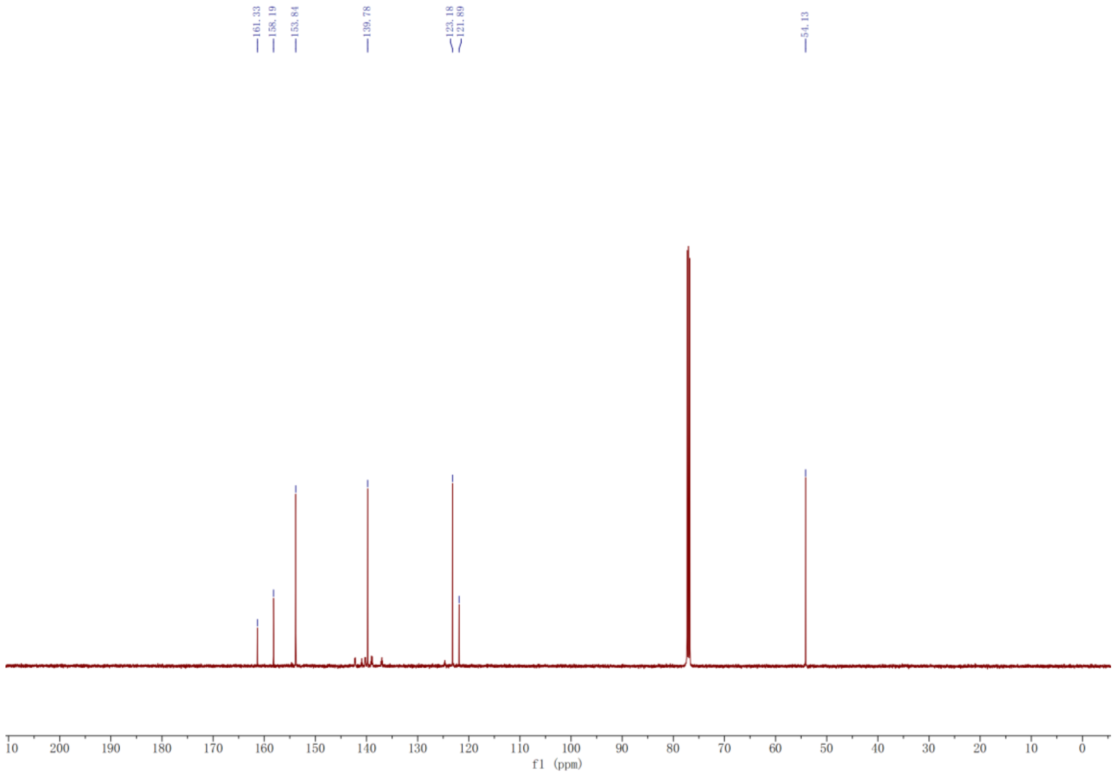
161.33
158.19
153.84
139.78
123.18
121.89
54.13
f1 (ppm)

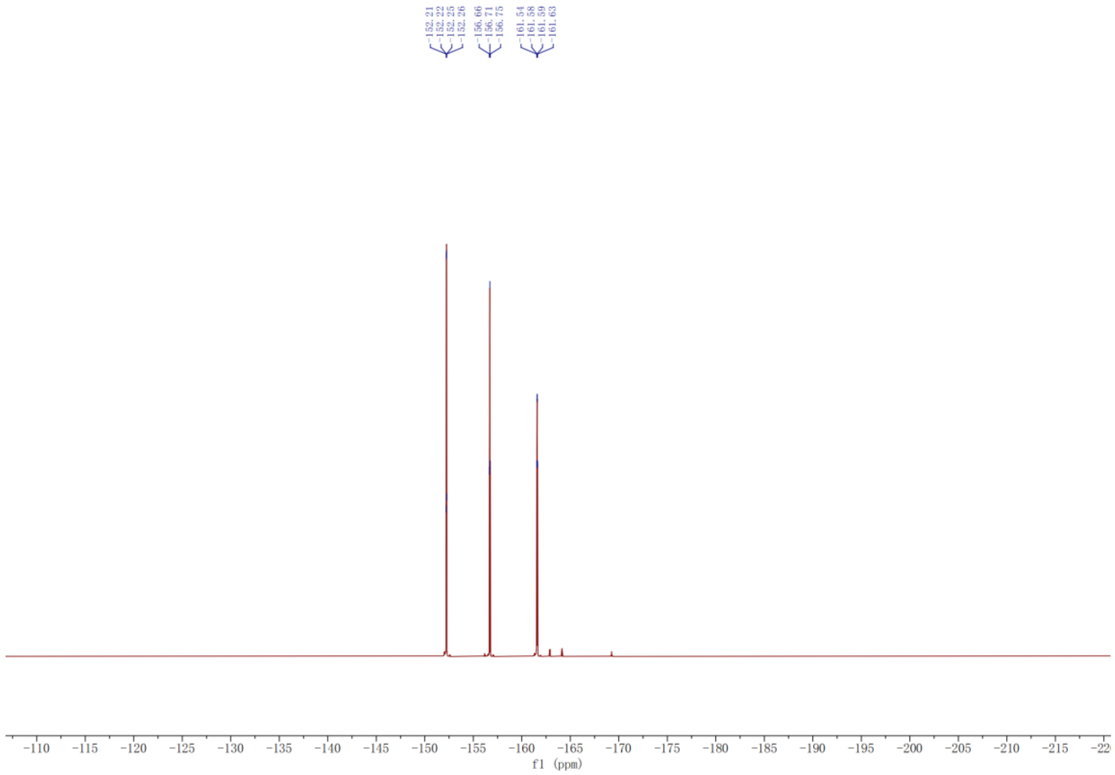
-152.21
-152.22
-152.25
-152.26
-156.66
-156.71
-156.75
-161.54
-161.58
-161.59
-161.63
f1 (ppm)

**Synthesis of HBC-DMAP analogue (3):**

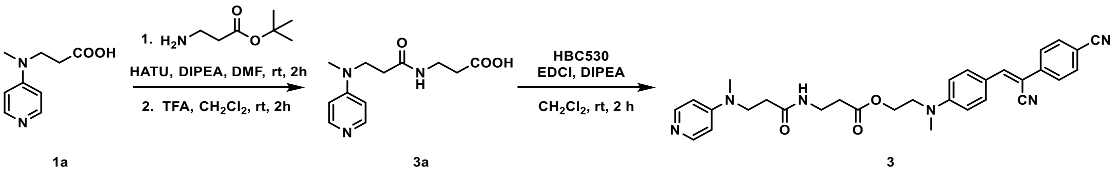


To a solution of 3-(methyl(pyridin-4-yl) amino) propanoic acid (**1a**, 200 mg, 1,10 mmol) and *tert*-butyl 3-aminopropanoate (160 mg, 1.10mmol) in DMF (5 mL) was added HATU (630 mg, 1.65 mmol) and DIPEA (400 μL, 2.20 mmol). The solution was stirred at room temperature for 2 h, then purified by preparative thin-layer chromatography (DCM:MeOH = 20:1) to afford *tert*-butyl 3-(3-(methyl(pyridin-4-yl) amino) propanamido) propanoate. Next, a solution of *tert*-butyl 3-(3-(methyl(pyridin-4-yl) amino) propanamido) propanoate in 1:1 TFA/DCM was stirred at rt for 2 h. The solvent was evaporated under reduced pressure to give intermediate **3a** that was used in the following reactions without further purification (120 mg, 43%).

To a solution of **3a** (120 mg, 0.48 mmol) and **HBC530** (145 mg, 0.48 mmol) in DCM (5 mL) was added EDCI (110 mg, 0.72 mmol) and DIPEA (340 μL, 2.40 mmol). The solution was stirred at room temperature for 2 h, then purified by preparative thin-layer chromatography (DCM:MeOH = 10:1) to afford compound **3** as a yellow solid (103 mg, 40%).

$^1$H NMR (600 MHz, DMSO) δ 8.21 (d, $J$ = 7.1 Hz, 2H), 8.12 (t, $J$ = 5.7 Hz, 1H), 8.03 (s, 1H), 7.93 (d, $J$ = 8.7 Hz, 4H), 7.09 – 7.01 (m, 2H), 6.92 – 6.87 (m, 2H), 4.23 (t, $J$ = 5.6 Hz, 2H), 3.76 (t, $J$ = 6.8 Hz, 2H), 3.73 (t, $J$ = 5.7 Hz, 2H), 3.22 (q, $J$ = 6.5 Hz, 2H), 3.10 (s, 3H), 3.05 (s, 3H), 2.38 (dt, $J$ = 13.3, 6.8 Hz, 4H).

$^{13}$C NMR (151 MHz, DMSO) δ 171.70, 170.28, 156.99, 151.70, 145.90, 139.98, 133.43, 132.39, 130.26, 125.99, 121.09, 119.24, 119.18, 112.13, 110.30, 107.60, 100.92, 61.77, 50.31, 48.68, 40.53, 38.97, 38.79, 38.55, 35.09, 34.11, 33.01.

ESI-MS $[M+H]^+$ calcd for $C_{31}H_{33}N_6O_3$: 537.2609, found: 537.2610.

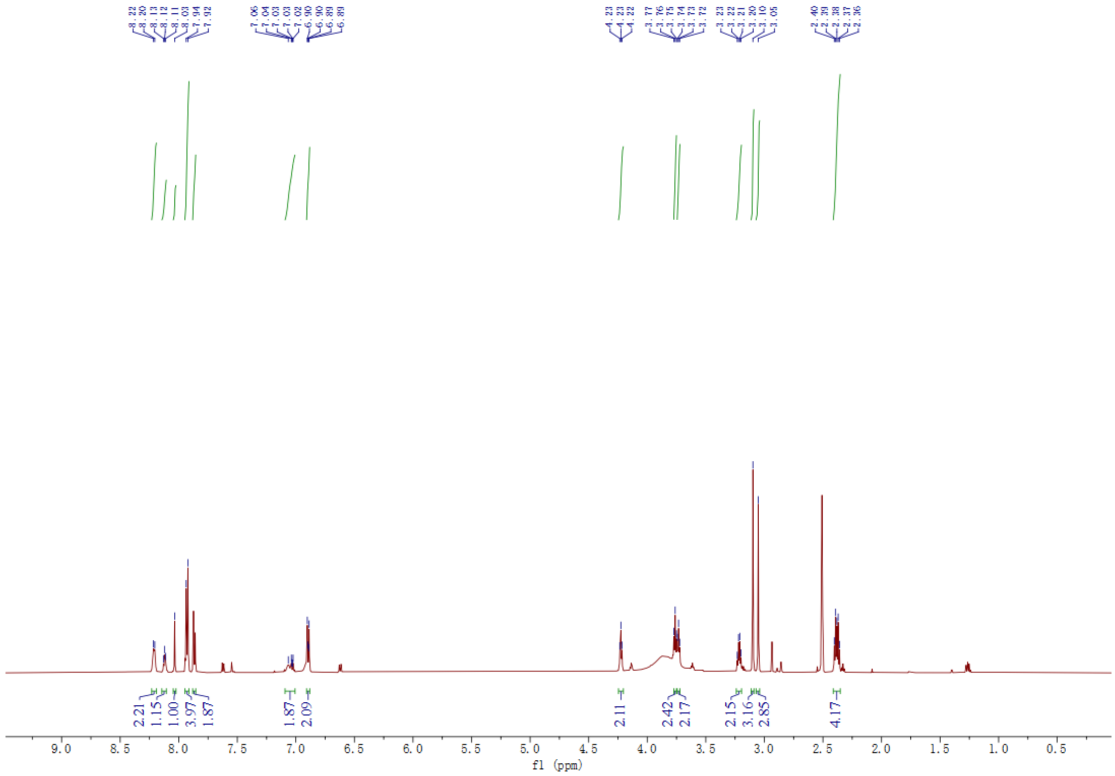

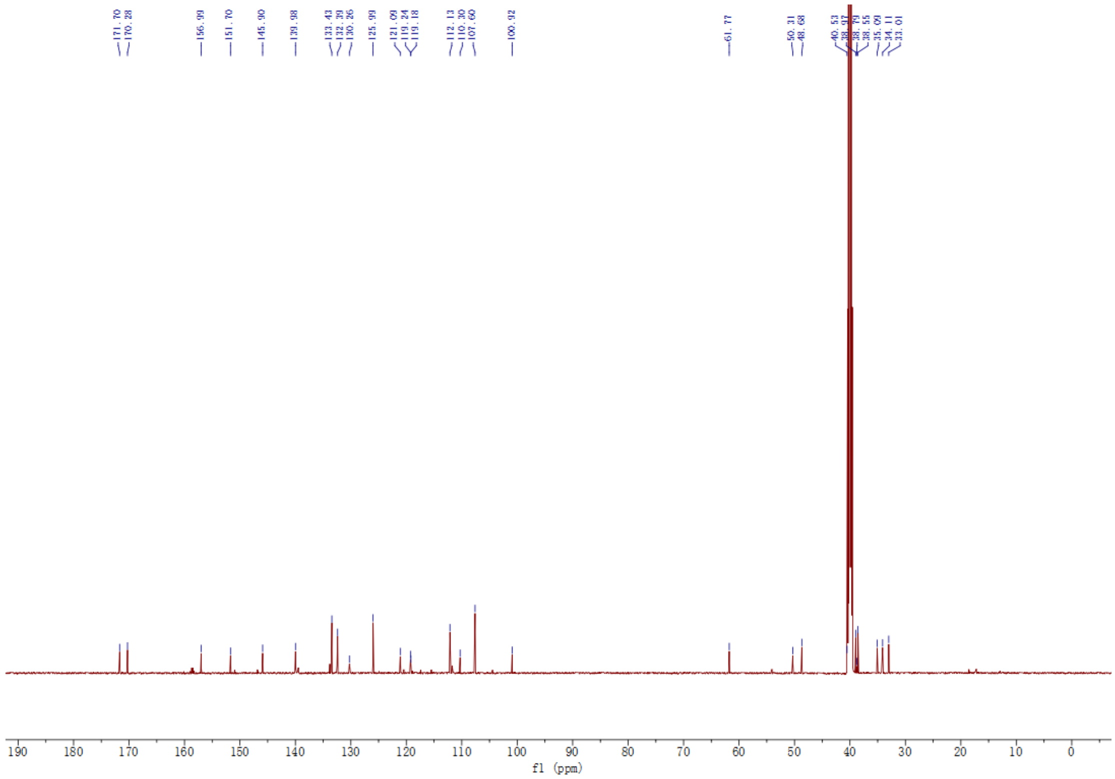

**Synthesis of HBC-DMAP analogue (4):**

The synthetic procedure for compound **4** was identical to that for compound **3**, except that tert-butyl 3-aminopropanoate was replaced with tert-butyl 3-(2-aminoethoxy) propanoate. Yellow solid (46%).

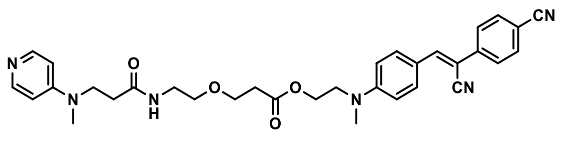


**4**

$^{1}$H NMR (500 MHz, DMSO) δ 8.09 (d, *J* = 6.0 Hz, 2H), 8.02 (s, 1H), 7.99 (m, 1H), 7.95 – 7.89 (m, 4H), 7.86 (d, *J* = 8.3 Hz, 2H), 6.88 (d, *J* = 8.9 Hz, 2H), 6.64 (d, *J* = 5.9 Hz, 2H), 4.23 (t, *J* = 5.5 Hz, 2H), 3.73 (t, *J* = 5.6 Hz, 2H), 3.60 (m, 2H), 3.53 (t, *J* = 6.2 Hz, 2H), 3.30 (t, *J* = 5.9 Hz, 2H), 3.13 (q, *J* = 5.5 Hz, 2H), 3.04 (s, 3H), 2.91 (s, 3H), 2.47 (t, *J* = 6.2 Hz, 2H), 2.31 (t, *J* = 6.8 Hz, 2H).

$^{13}$C NMR (151 MHz, DMSO) δ 171.51, 170.25, 157.00, 151.74, 145.89, 140.00, 133.42, 132.38, 130.26, 126.00, 121.07, 119.24, 112.15, 111.76, 110.29, 107.61, 100.91, 69.19, 66.03, 61.78, 50.30, 48.73, 40.53, 38.90, 38.55, 34.98, 33.04.

ESI-MS $[M+H]^+$ calcd for $C_{33}H_{37}N_6O_4$: 581.2871, found: 581.2879.

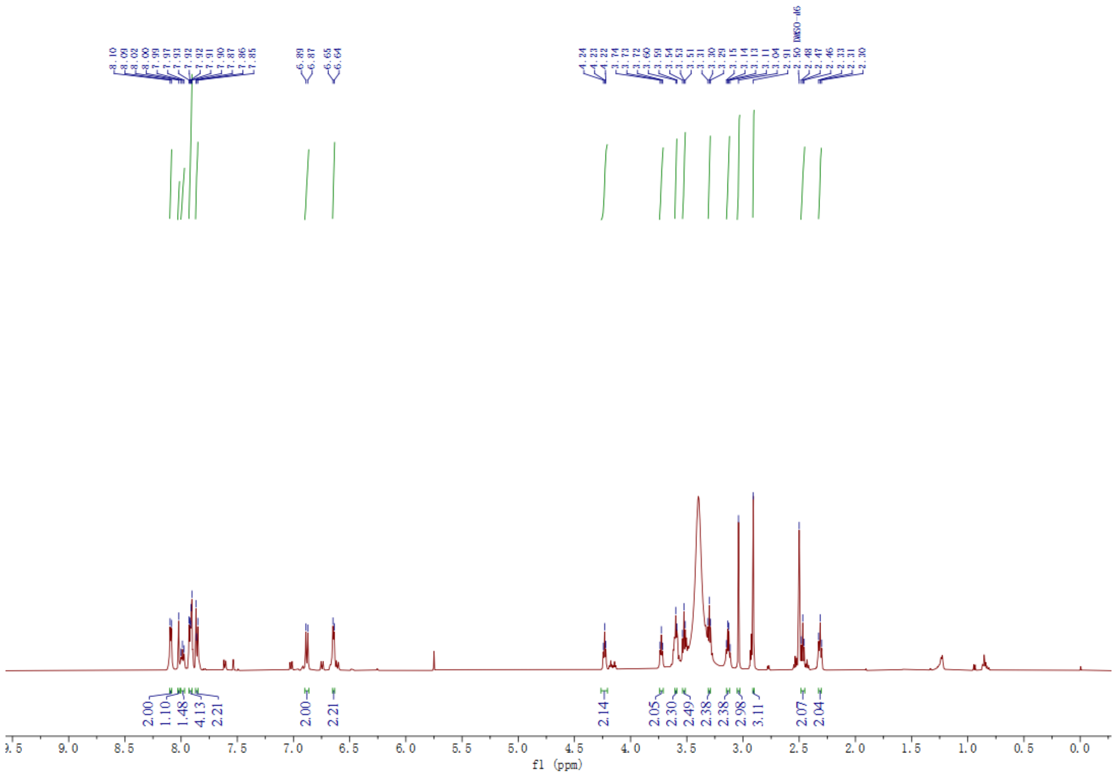

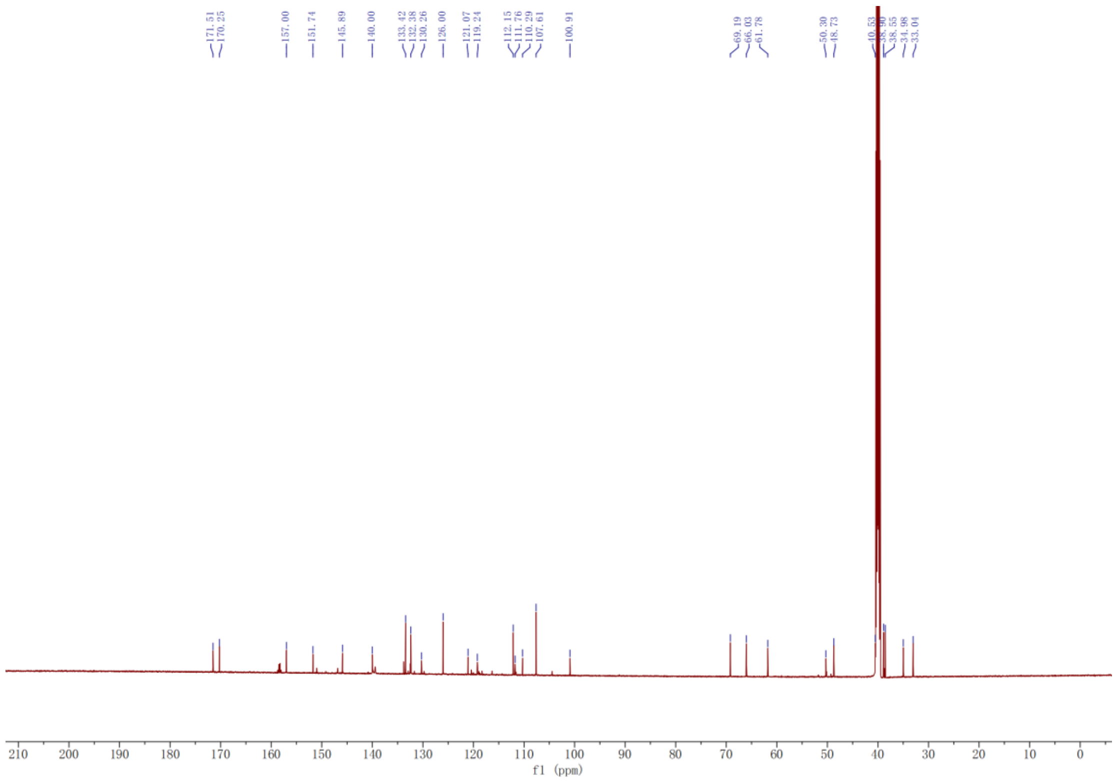

**Synthesis of HBC-DMAP analogue (5):**

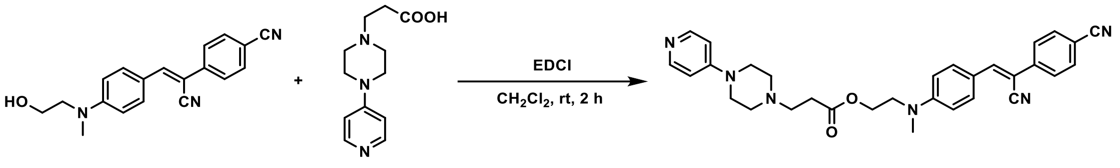


Intermediate 5a were prepared according to previous work[7].

**5** was obtained by the same procedures as **1**. Yellow solid (75%).

$^{1}$H NMR (500 MHz, DMSO) δ 8.36 (d, *J* = 7.6 Hz, 2H), 8.04 (s, 1H), 7.93 (dd, *J* = 8.8, 7.1 Hz, 4H), 7.89 – 7.83 (m, 2H), 7.28 (d, *J* = 7.5 Hz, 2H), 6.91 (d, *J* = 9.1 Hz, 2H), 4.29 (t, *J* = 5.6 Hz, 2H), 3.76 (t, *J* = 5.6 Hz, 2H), 3.32 (t, *J* = 7.5 Hz, 2H), 3.07 (s, 3H), 2.84 (t, *J* = 7.3 Hz, 2H).

$^{13}$C NMR (126 MHz, DMSO) δ 170.53, 158.94, 158.67, 157.23, 151.71, 145.89, 140.72, 139.95, 133.42, 132.41, 125.99, 121.13, 119.23, 112.16, 108.56, 101.02, 62.43, 51.42, 50.83, 50.23, 43.43, 38.93, 29.08.

ESI-MS $[M+H]^+$ calcd for $C_{31}H_{33}N_6O_2$: 521.2660, found: 521.2673.

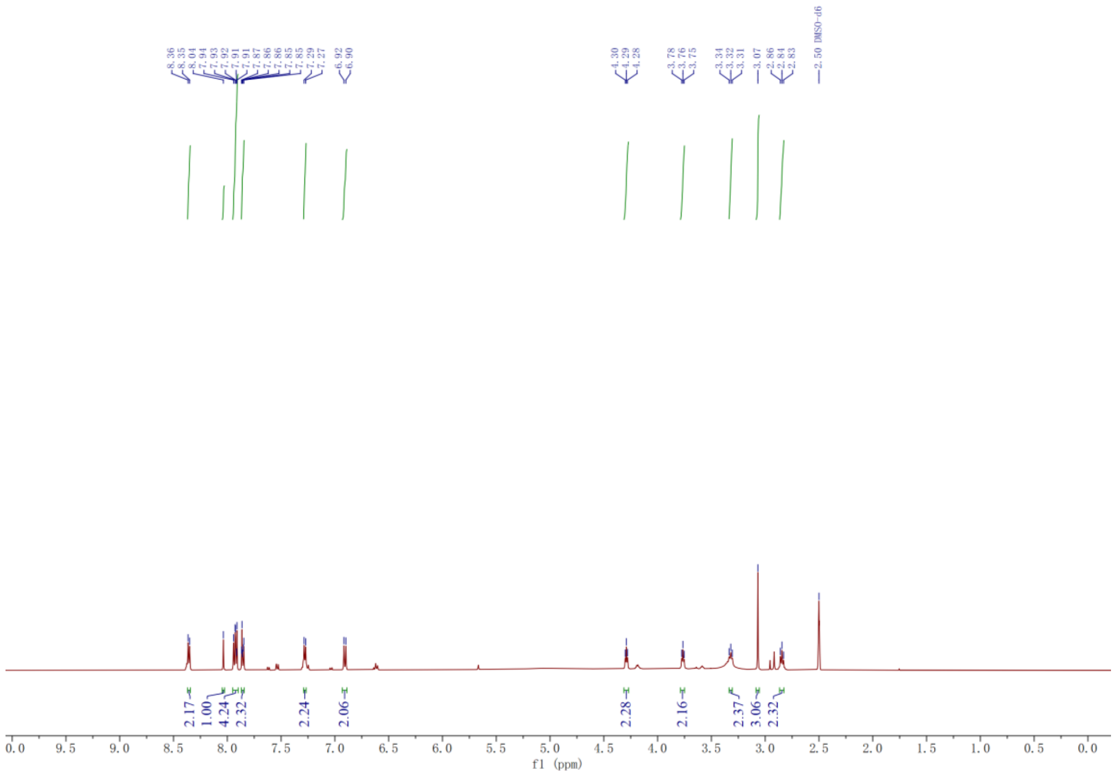
8.36
8.35
8.04
7.94
7.93
7.92
7.91
7.91
7.87
7.86
7.86
7.85
7.29
7.27
6.92
6.90
4.30
4.29
4.28
3.78
3.76
3.75
3.34
3.32
3.31
3.07
2.86
2.84
2.83
2.50 DMSO-d6
2.17
1.00
4.24
2.32
2.24
2.06
2.28
2.16
2.37
3.06
2.32
f1 (ppm)

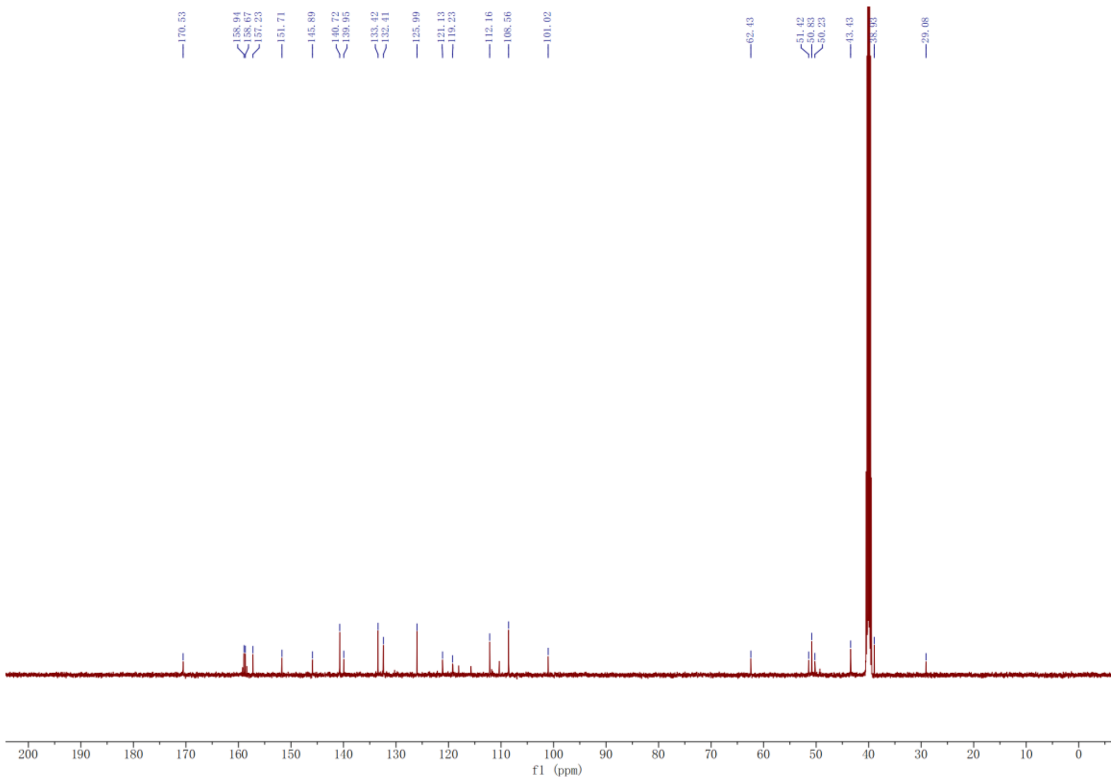
170.53
158.94
158.67
157.23
151.71
145.89
140.72
139.95
133.42
132.41
125.99
121.13
119.23
112.16
108.56
101.02
62.43
51.42
50.83
50.23
43.43
38.93
29.08
f1 (ppm)

**Synthesis of HBC-PyOx (6):**

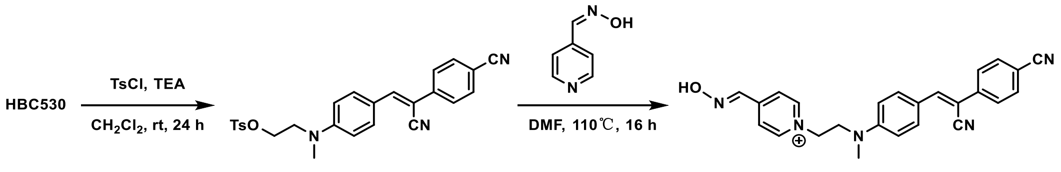


To a cooled stirring solution of HBC530 (120 mg, 0.40 mmol) and TEA (50 mg, 0,44 mmol) in 10 ml anhydrous $CH_2Cl_2$ at 0 °C, *para*-toluenesulfonyl chloride (TsCl, 100 mg, 0,40 mmol) in 2 ml $CH_2Cl_2$ was added dropwise. After addition, the mixture was warmed to room temperature and stirred for 24 h. The reaction mixture was washed with a saturated solution of $NaHCO_3$. The organic phase was dried over anhydrous $Na_2SO_4$. After filtration and removal of solvent under reduced pressure, the crude product was used directly without any further purification. After dissolution in 5 ml DMF, then pyridine-4-aldoxime (50 mg, 0.40 mmol) was added. The resulting mixture was stirred at 110 °C for 16 h. The solvent was removed then purified by preparative reversed-phase HPLC (0% to 100% ACN in 0.1% TFA water over 35 min). The product-containing fractions were dried by lyophilisation to afford **HBC-PyOx** (**6**) as a yellow solid (50 mg, 30%).

$^1$H NMR (500 MHz, MeOD) δ 8.85 (d, *J* = 6.5 Hz, 2H), 8.26 (s, 1H), 8.13 (d, *J* = 6.9 Hz, 2H), 7.86 (s, 1H), 7.83 (d, *J* = 8.1 Hz, 3H), 7.79 (s, 1H), 7.77 (d, *J* = 3.5 Hz, 2H), 6.75 (d, *J* = 9.1 Hz, 2H), 4.13 (t, *J* = 5.7 Hz, 2H), 3.05 (s, 3H), 2.66 (s, 2H).

$^{13}$C NMR (126 MHz, MeOD) δ 150.89, 150.30, 145.21, 144.68, 144.09, 139.80, 132.51, 131.73, 125.67, 123.97, 122.40, 118.00, 111.40, 110.97, 103.11, 58.57, 51.85, 39.03, 37.23.

ESI-MS $[M]^+$ calcd for $C_{25}H_{22}N_5O$: 408.1819, found: 408.1821.

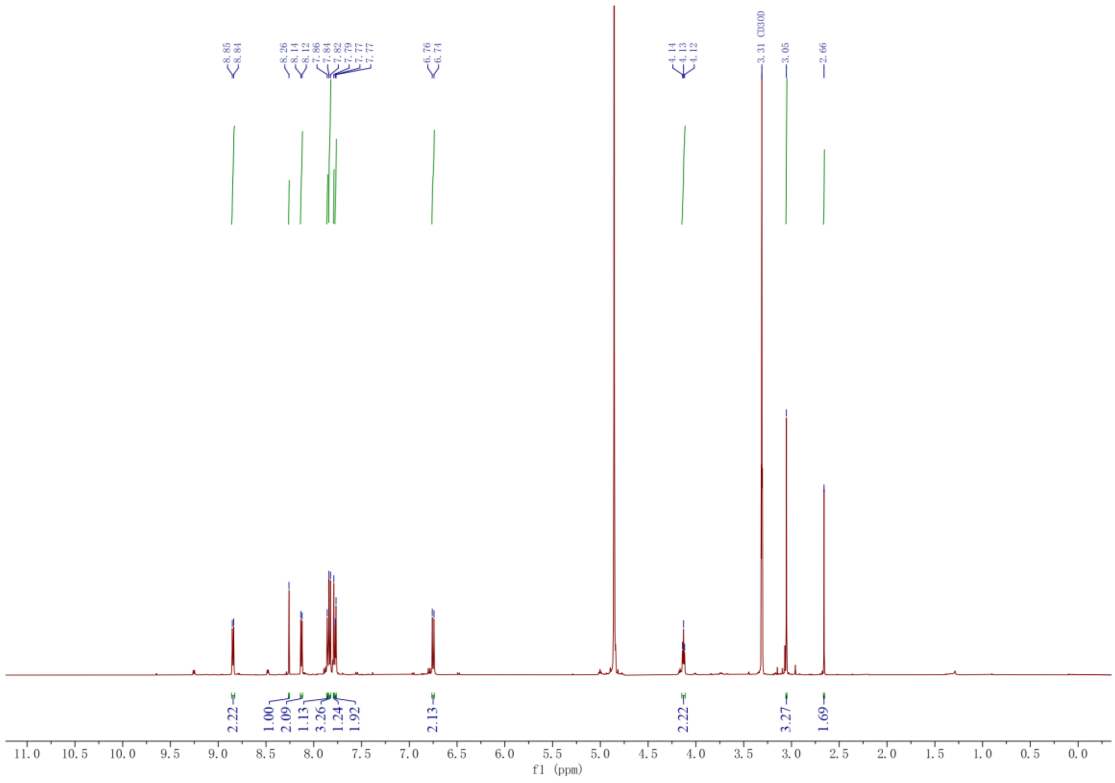
8.85
8.84
8.26
8.14
8.12
7.86
7.84
7.82
7.79
7.77
7.77
6.76
6.74
4.14
4.13
4.12
3.31 CD3OD
3.05
2.66
2.22
1.00
2.09
1.13
3.26
1.24
1.92
2.13
2.22
3.27
1.69
11.0 10.5 10.0 9.5 9.0 8.5 8.0 7.5 7.0 6.5 6.0 5.5 5.0 4.5 4.0 3.5 3.0 2.5 2.0 1.5 1.0 0.5 0.0
f1 (ppm)

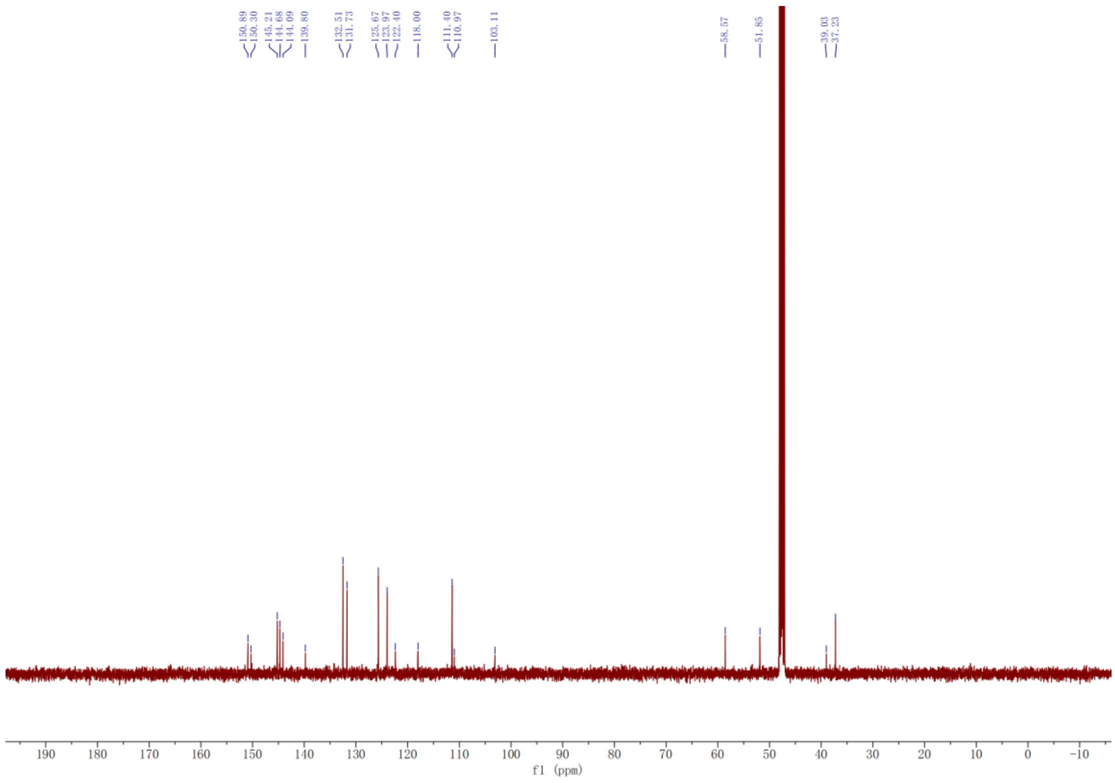
150.89
150.30
145.21
144.68
144.09
139.80
132.51
131.73
125.67
123.97
122.40
118.00
111.40
110.97
103.11
58.57
51.85
39.03
37.23
190 180 170 160 150 140 130 120 110 100 90 80 70 60 50 40 30 20 10 0 -10
f1 (ppm)

#### Synthesis of HBC-PyOx analogues (7-9):

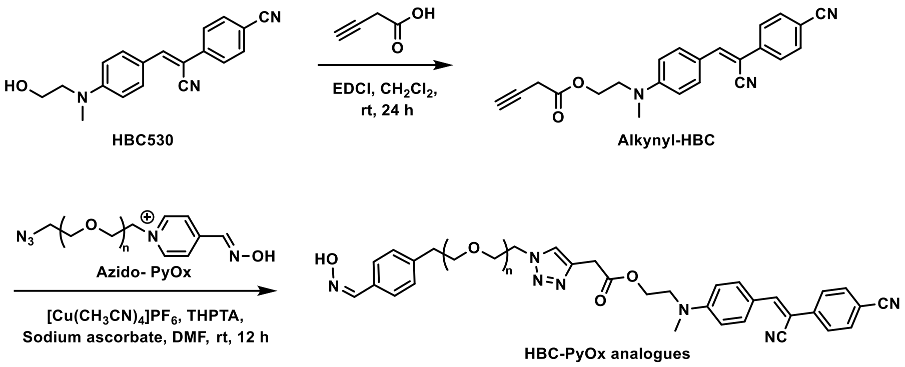


To a solution of but-3-ynoic acid (1.2 equiv.) and HBC530 (1.0 equiv.) in DCM was added EDCI (1.5 equiv.). The solution was stirred at room temperature for 24 h, then purified by preparative thin-layer chromatography (DCM:MeOH = 20:1) to afford alkynyl-HBC as a yellow solid.

Intermediate Azido-PyOx with different linkers were prepared according to previous work[8].

To a solution of alkynyl-HBC (1.0 equiv.) in dry DMF was added azido-PyOx (1.2 equiv.), $[Cu(CH_3CN)_4]PF_6$ (1.5 equiv.), THPTA (1.5 equiv.) and sodium ascorbate (5.0 equiv.). The mixture was allowed to stir at room temperature for 12 h. After removal of the solvent, the crude residue was purified by preparative reversed-phase HPLC (0% to 100% ACN in 0.1% TFA water over 35 min). The product-containing fractions were dried by lyophilisation to afford **HBC-PyOx analogues** (**7-9**) as a yellow solid.

**HBC-PyOx analogue** (7)

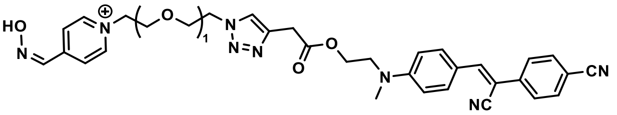


HBC-PyOx analogue (7)

Yellow solid (70%).

$^1$H NMR (500 MHz, MeOD) δ 8.67 (d, *J* = 6.9 Hz, 2H), 8.29 (s, 1H), 8.10 (d, *J* = 6.9 Hz, 2H), 7.92 (d, *J* = 9.1 Hz, 2H), 7.84 (m, 3H), 7.82 – 7.76 (m, 4H), 7.74 (s, 1H), 6.87 (d, *J* = 9.1 Hz, 2H), 4.73 – 4.68 (m, 2H), 4.57 – 4.51 (m, 2H), 4.42 (t, *J* = 5.6 Hz, 2H), 3.94 – 3.86 (m, 4H), 3.81 (t, *J* = 5.6 Hz, 2H), 3.72 (s, 2H), 3.23 (q, *J* = 7.3 Hz, 1H), 3.19 – 3.13 (m, 2H), 3.11 (s, 1H), 3.10 (s, 3H).

$^{13}$C NMR (126 MHz, MeOD) δ 170.36, 151.64, 149.95, 145.09, 144.82, 144.34, 140.06, 132.50, 131.82, 129.74, 125.74, 125.47, 123.95, 123.70, 121.20, 118.13, 111.42, 110.63, 101.46, 68.71, 68.33, 62.02, 60.49, 49.99, 49.69, 44.33, 37.52, 30.57, 22.37, 21.65.

ESI-MS $[M]^+$ calcd for $C_{33}H_{33}N_8O_4$: 605.2619, found: 605.2628.

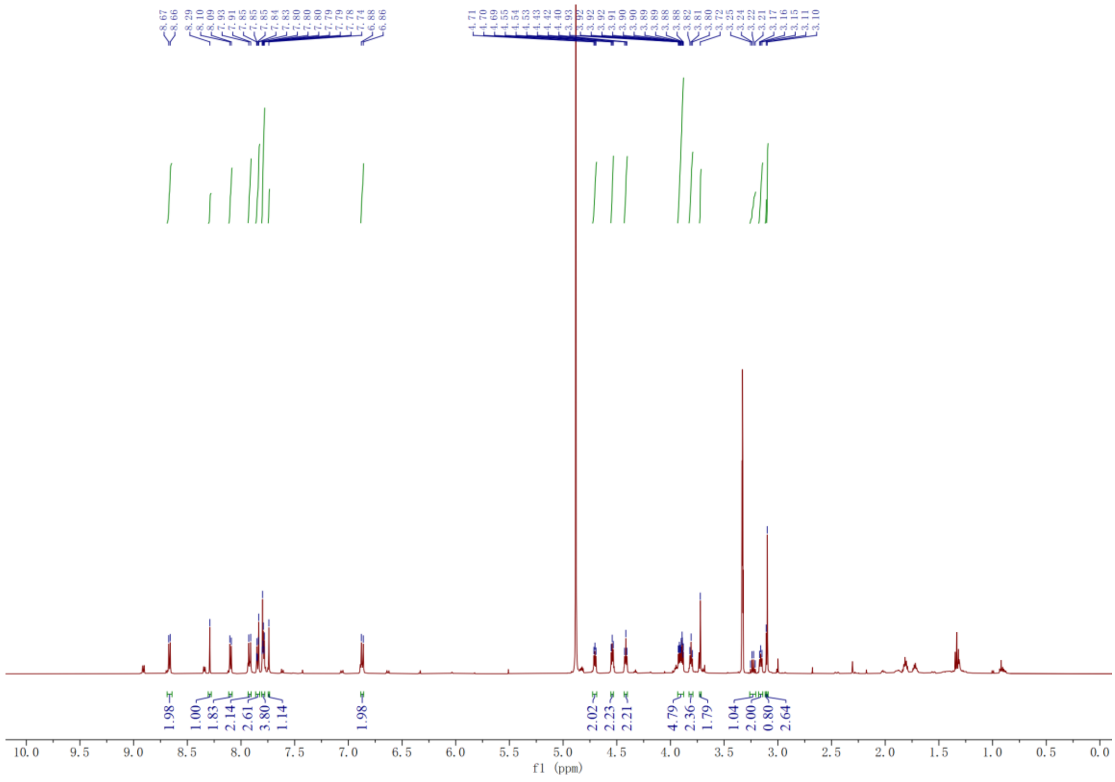
f1 (ppm)

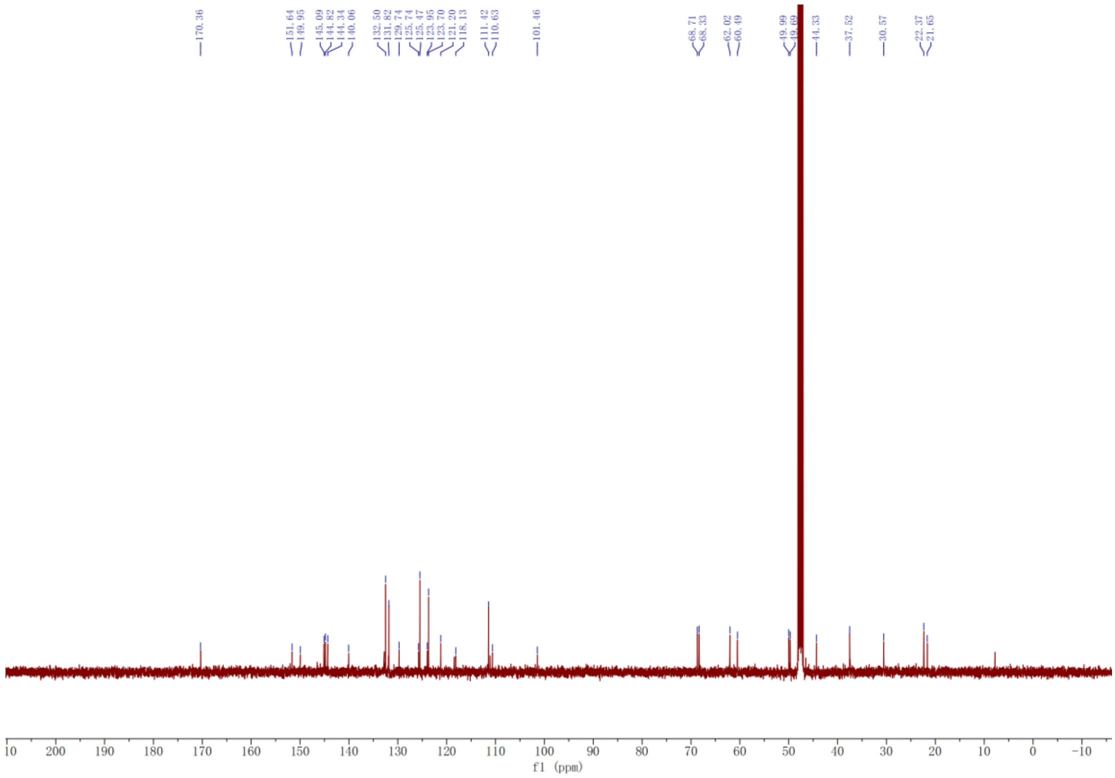
f1 (ppm)

**HBC-PyOx analogue (8)**

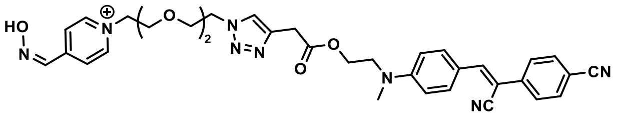


HBC-PyOx analogue (8)

Yellow solid (72%).

$^1$H NMR (500 MHz, MeOD) δ 8.76 (d, *J* = 6.9 Hz, 2H), 8.29 (s, 1H), 8.15 (d, *J* = 6.9 Hz, 2H), 7.91 (d, *J* = 9.0 Hz, 2H), 7.85 (s, 1H), 7.84 (s, 2H), 7.80 (d, *J* = 2.2 Hz, 2H), 7.78 (s, 1H), 6.86 (d, *J* = 9.1 Hz, 2H), 4.68 – 4.62 (m, 2H), 4.56 – 4.51 (m, 2H), 4.41 (t, *J* = 5.5 Hz, 2H), 3.87 – 3.79 (m, 6H), 3.77 (d, *J* = 0.6 Hz, 2H), 3.59 – 3.52 (m, 4H), 3.09 (s, 3H).

$^{13}$C NMR (126 MHz, MeOD) δ 170.51, 151.58, 149.96, 145.03, 144.22, 140.02, 132.78, 132.51, 131.82, 129.74, 125.48, 124.12, 123.65, 121.17, 118.15, 111.40, 110.66, 101.51, 70.18, 70.07, 68.85, 68.75, 62.03, 60.72, 50.04, 50.00, 48.11, 37.61, 30.63.

ESI-MS $[M]^+$ calcd for $C_{35}H_{37}N_8O_5$: 649.2881, found: 649.2883.

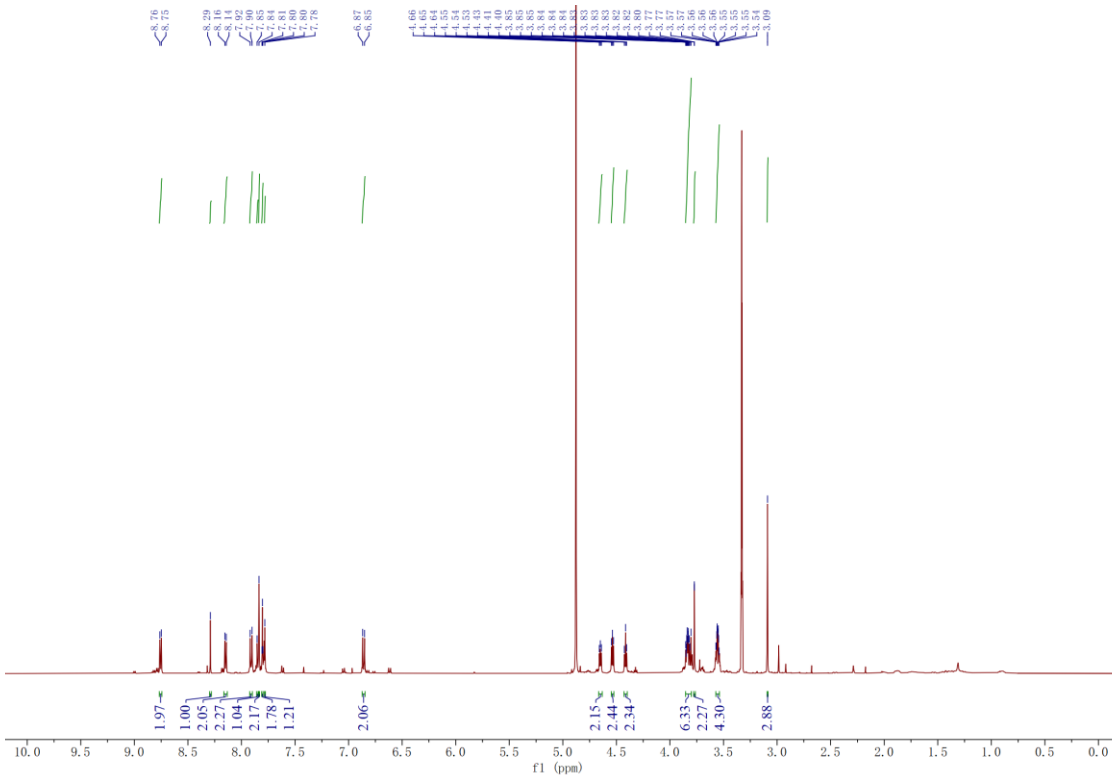

8.76 8.75 8.29 8.16 8.14 7.92 7.90 7.85 7.84 7.81 7.80 7.80 7.78 6.87 6.85 4.66 4.65 4.64 4.55 4.54 4.53 4.43 4.41 4.40 3.85 3.85 3.85 3.84 3.84 3.84 3.83 3.83 3.83 3.82 3.80 3.77 3.77 3.57 3.57 3.56 3.56 3.56 3.55 3.55 3.55 3.54 3.09
1.97 1.00 2.05 2.27 1.04 2.17 1.78 1.21 2.06 2.15 2.44 2.34 6.33 2.27 4.30 2.88
f1 (ppm)


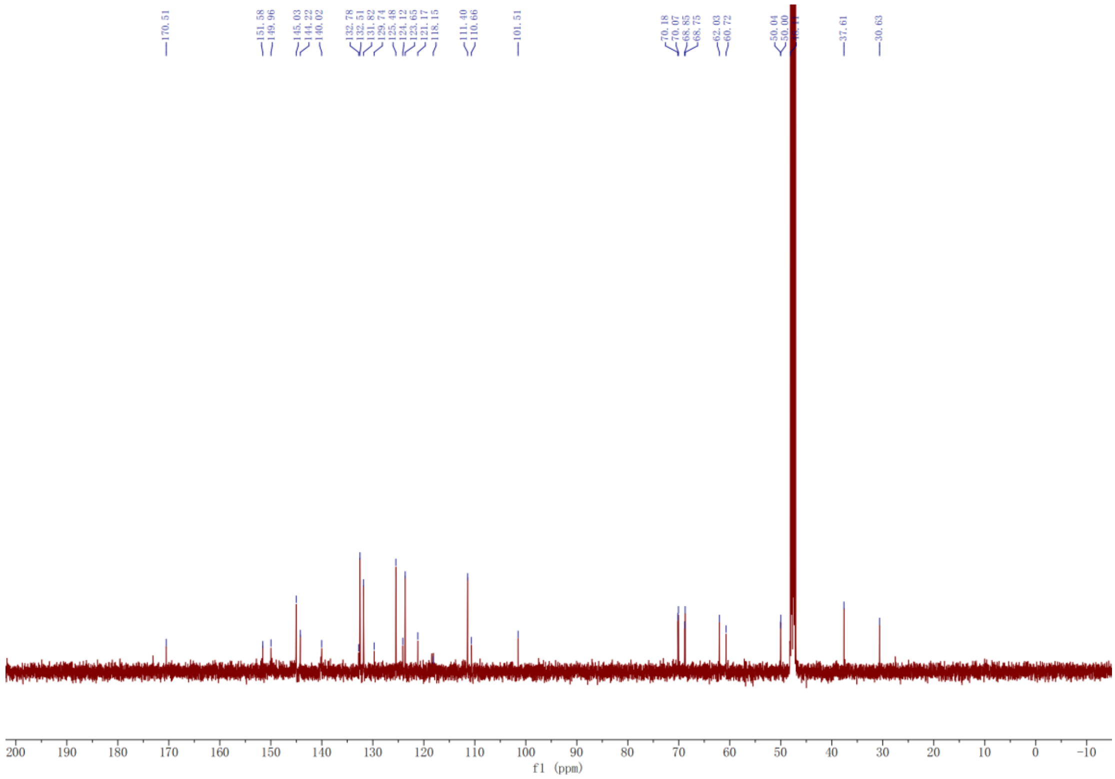

170.51 151.58 149.96 145.03 144.22 140.02 132.78 132.51 131.82 129.74 125.48 124.12 123.65 121.17 118.15 111.40 110.66 101.51 70.18 70.07 68.85 68.75 62.03 60.72 50.04 50.00 37.61 30.63
f1 (ppm)

**HBC-PyOx analogue (9)**

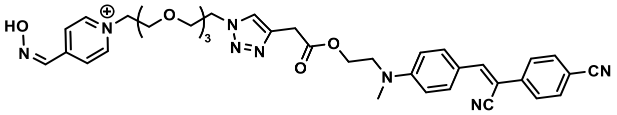


HBC-PyOx analogue (9)

Yellow solid (56%).

$^{1}$H NMR (500 MHz, MeOD) δ 8.82 (d, *J* = 6.8 Hz, 2H), 8.28 (s, 1H), 8.13 (d, *J* = 7.0 Hz, 2H), 7.90 (d, *J* = 9.1 Hz, 2H), 7.83 (s, 1H), 7.81 (d, *J* = 3.0 Hz, 3H), 7.78 (d, *J* = 2.7 Hz, 3H), 7.76 (s, 1H), 6.84 (d, *J* = 9.1 Hz, 2H), 4.72 – 4.67 (m, 2H), 4.56 – 4.50 (m, 2H), 4.37 (t, *J* = 5.5 Hz, 2H), 3.94 – 3.89 (m, 2H), 3.89 – 3.84 (m, 2H), 3.77 (t, *J* = 5.5 Hz, 2H), 3.71 (s, 2H), 3.58 – 3.52 (m, 4H), 3.51 – 3.45 (m, 4H), 3.05 (s, 3H).

$^{13}$C NMR (126 MHz, MeOD) δ 170.26, 151.63, 149.89, 145.13, 145.09, 144.26, 140.10, 140.05, 132.52, 131.84, 125.47, 124.05, 123.61, 121.18, 118.45, 118.15, 111.42, 110.61, 101.45, 70.06, 70.04, 69.89, 68.90, 68.66, 62.02, 60.67, 50.02, 49.95, 37.51, 30.71.

ESI-MS $[M]^+$ calcd for $C_{37}H_{41}N_8O_6$: 693.3144, found: 693.3158.

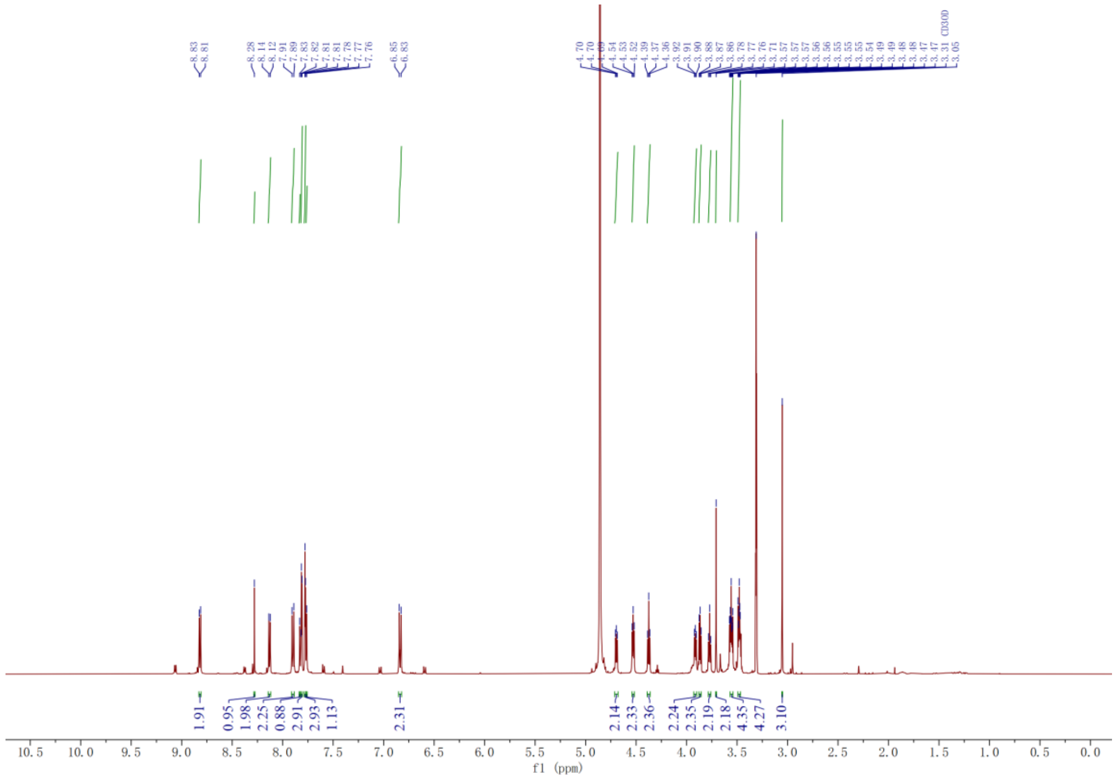
8.83
8.81
8.28
8.14
8.12
7.91
7.89
7.83
7.82
7.81
7.81
7.78
7.77
7.76
6.85
6.83
4.70
4.70
4.69
4.54
4.53
4.52
4.39
4.37
4.36
3.92
3.91
3.90
3.88
3.87
3.86
3.78
3.77
3.76
3.71
3.57
3.57
3.57
3.56
3.56
3.55
3.55
3.55
3.54
3.49
3.49
3.48
3.47
3.31
3.05
CD3OD
1.91
0.95
1.98
2.25
0.88
2.91
2.93
1.13
2.31
2.14
2.33
2.36
2.24
2.35
2.19
2.18
4.35
4.27
3.10
f1 (ppm)

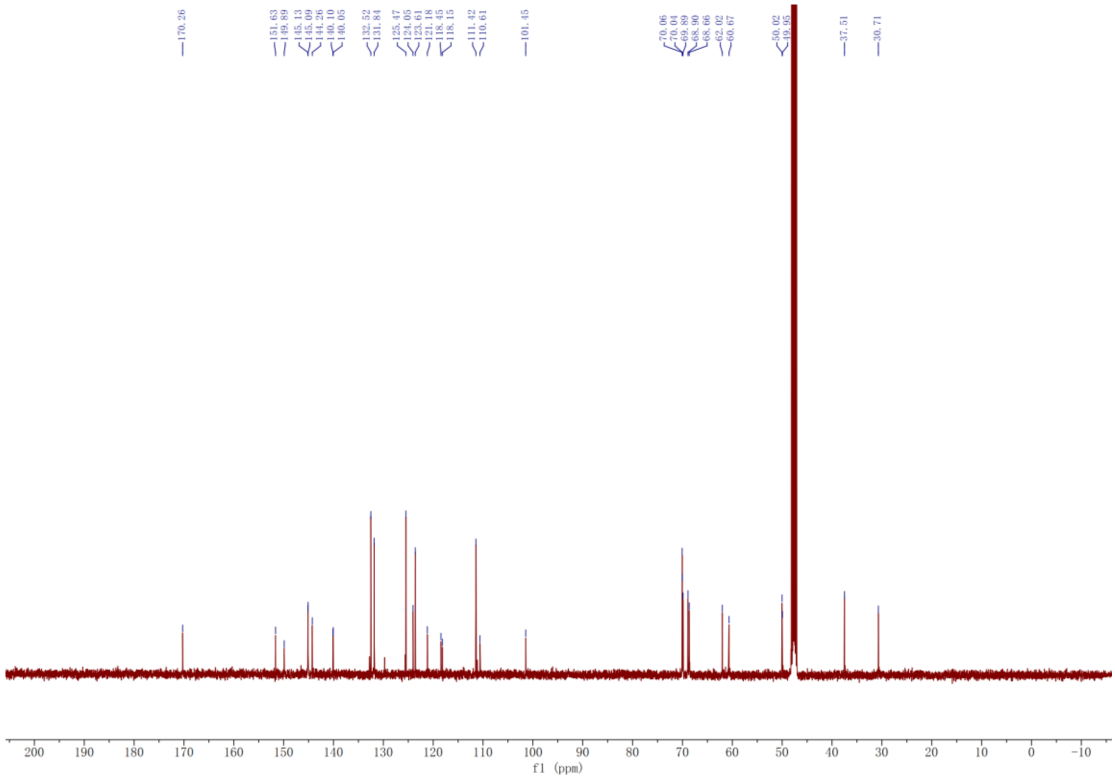
170.26
151.63
149.89
145.13
145.09
144.26
140.10
140.05
132.52
131.84
125.47
124.65
123.61
121.18
118.45
118.15
111.42
110.61
101.45
70.06
70.04
69.89
68.90
68.66
62.02
60.67
50.02
49.95
37.51
30.71
f1 (ppm)

#### Synthesis of HBC-Imidazole (10):

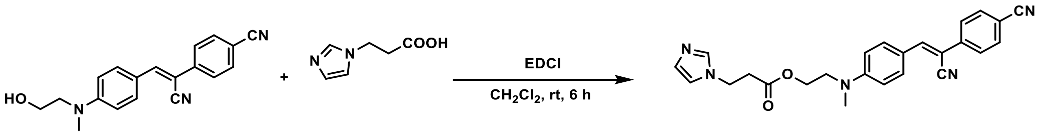


**10** was obtained by the same procedures as **1**. White solid (45%).

$^{1}$H NMR (500 MHz, DMSO) δ 9.08 (s, 1H), 8.03 (s, 1H), 7.92 (dd, *J* = 8.6, 5.7 Hz, 4H), 7.87 (d, *J* = 8.4 Hz, 2H), 7.72 (s, 1H), 7.64 (s, 1H), 6.87 (d, *J* = 9.0 Hz, 2H), 4.38 (t, *J* = 6.7 Hz, 2H), 4.25 (t, *J* = 5.6 Hz, 2H), 3.72 (t, *J* = 5.6 Hz, 2H), 3.02 (s, 3H), 2.96 (t, *J* = 6.6 Hz, 2H).

$^{13}$C NMR (126 MHz, DMSO) δ 170.73, 158.37, 151.67, 145.92, 139.98, 136.28, 133.44, 126.03, 122.43, 121.15, 120.40, 119.23, 112.14, 110.33, 101.06, 62.24, 50.26, 44.57, 38.90, 34.06.

ESI-MS $[M+H]^+$ calcd for $C_{25}H_{23}N_5O_2$: 426.1925, found: 423.1923.

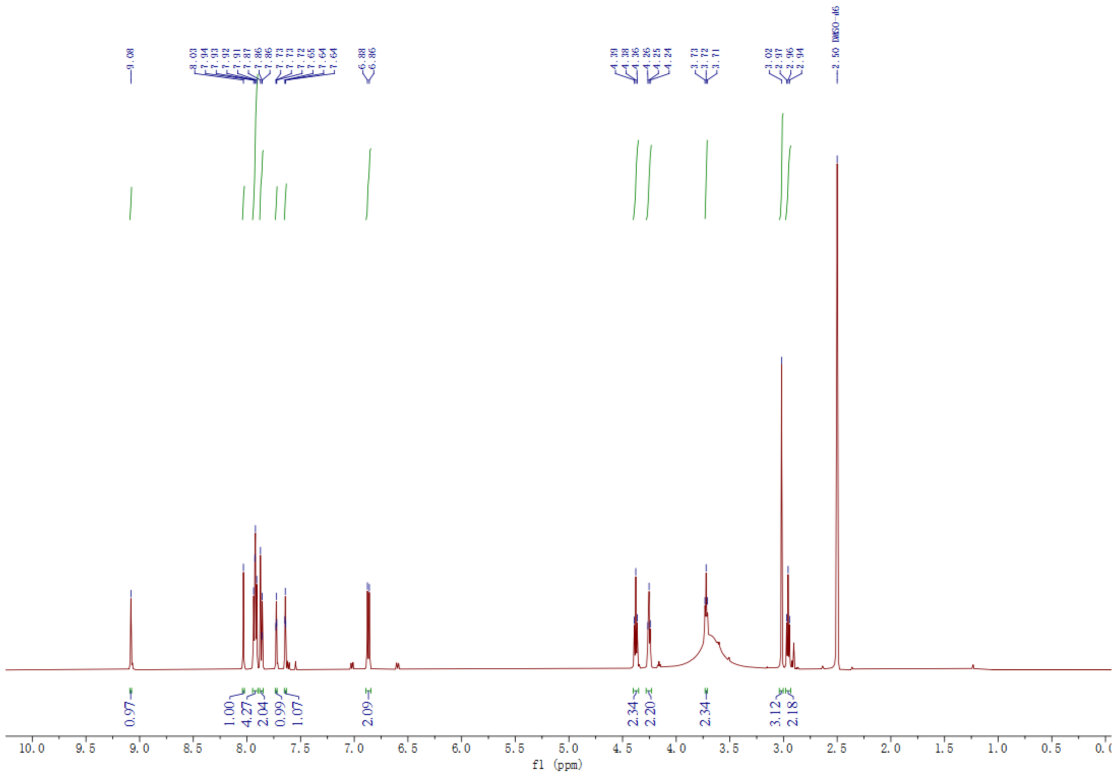
9.08
8.03
7.94
7.93
7.92
7.91
7.87
7.86
7.86
7.73
7.73
7.72
7.65
7.64
7.64
6.88
6.86
4.39
4.38
4.36
4.26
4.25
4.24
3.73
3.72
3.71
3.02
2.97
2.96
2.94
2.50 DMSO-d6
0.97
1.00
4.27
2.04
0.99
1.07
2.09
2.34
2.20
2.34
3.12
2.18
f1 (ppm)

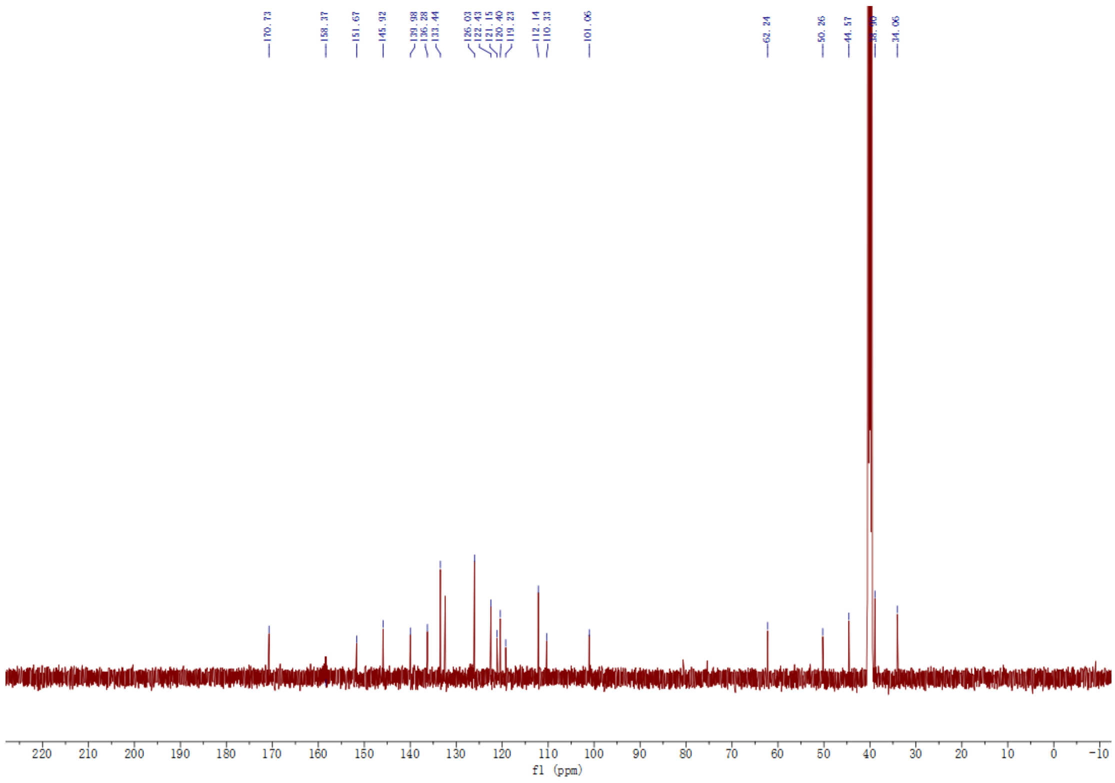
170.73
158.37
151.67
145.92
139.98
136.28
133.44
126.03
122.43
121.15
120.40
119.23
112.14
110.33
101.06
62.24
50.26
44.57
34.06
f1 (ppm)

**Synthesis of acylating reagents $N_3$-PEG-PFP (11-13):**

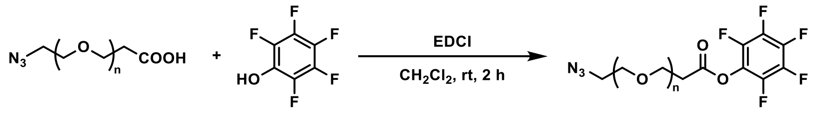


The synthetic procedure for compound **11-13** was identical to that for compound **2**, except that 2-(azidomethyl) nicotinic acid (2a) was replaced with Azido-COOH.

**$N_3$-2PEG-PFP (11)**

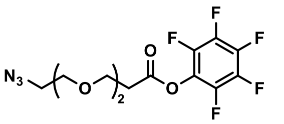
$N_3$-2PEG-PFP (11)

Colorless oil (65%).

$^1$H NMR (500 MHz, $CDCl_3$) δ 3.89 (t, $J$ = 6.2 Hz, 2H), 3.68 (m, 6H), 3.39 (t, $J$ = 5.0 Hz, 2H), 2.95 (t, $J$ = 6.2 Hz, 2H).

$^{13}$C NMR (126 MHz, $CDCl_3$) δ 167.57, 142.13, 140.14, 138.69, 136.92, 70.76, 70.65, 70.12, 66.13, 50.70, 34.45.

$^{19}$F NMR (471 MHz, $CDCl_3$) δ -152.54 – -152.63 (m), -157.97 (t, $J$ = 21.7 Hz), -162.28 – -162.43 (m).

ESI-MS $[M+Na]^+$ calcd for $C_{13}H_{12}F_5N_3O_4Na$: 392.0640, found:392.0641.

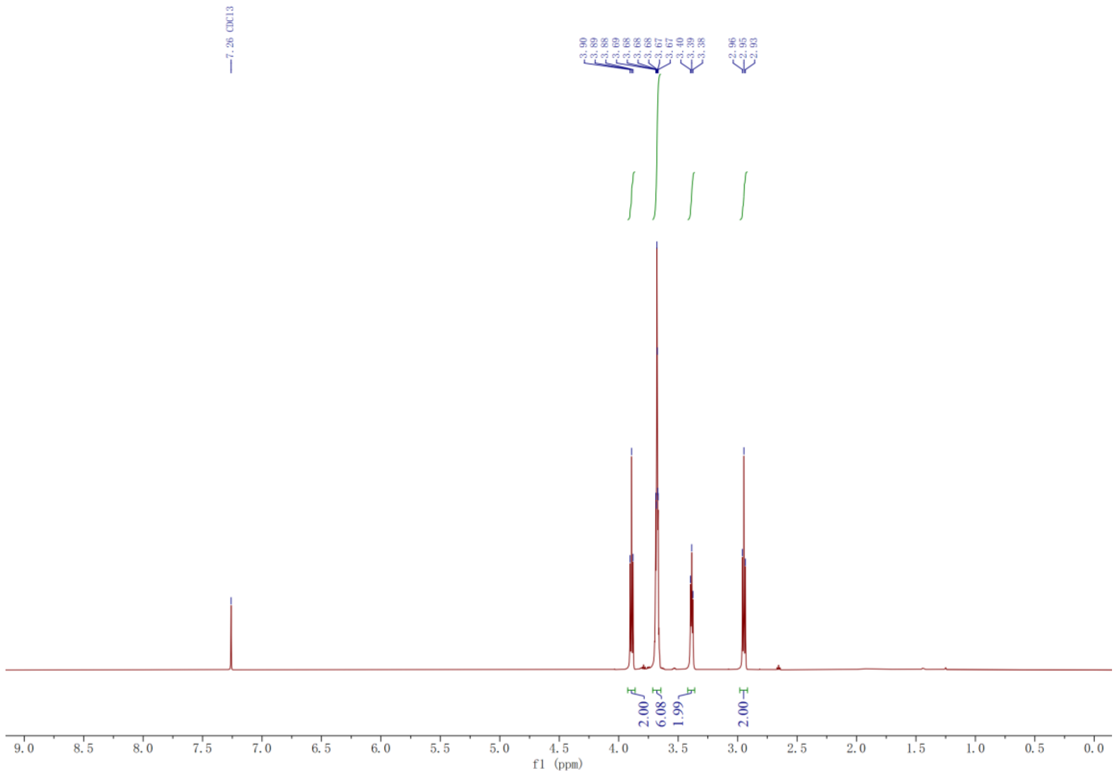
7.26 CDCl3
3.90
3.89
3.88
3.69
3.68
3.68
3.68
3.67
3.67
3.40
3.39
3.38
2.96
2.95
2.93
2.00
6.08
1.99
2.00
9.0 8.5 8.0 7.5 7.0 6.5 6.0 5.5 5.0 4.5 4.0 3.5 3.0 2.5 2.0 1.5 1.0 0.5 0.0
f1 (ppm)

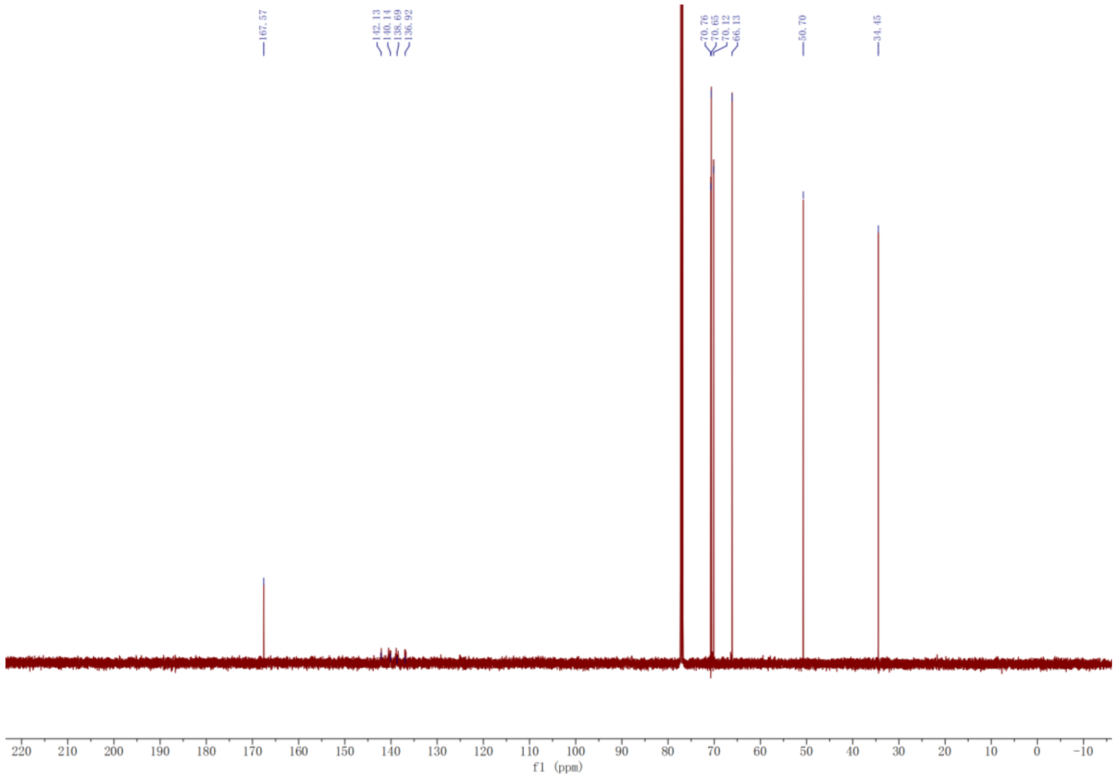
167.57
142.13
140.14
138.69
136.92
70.76
70.65
70.12
66.13
50.70
34.45
220 210 200 190 180 170 160 150 140 130 120 110 100 90 80 70 60 50 40 30 20 10 0 -10
f1 (ppm)

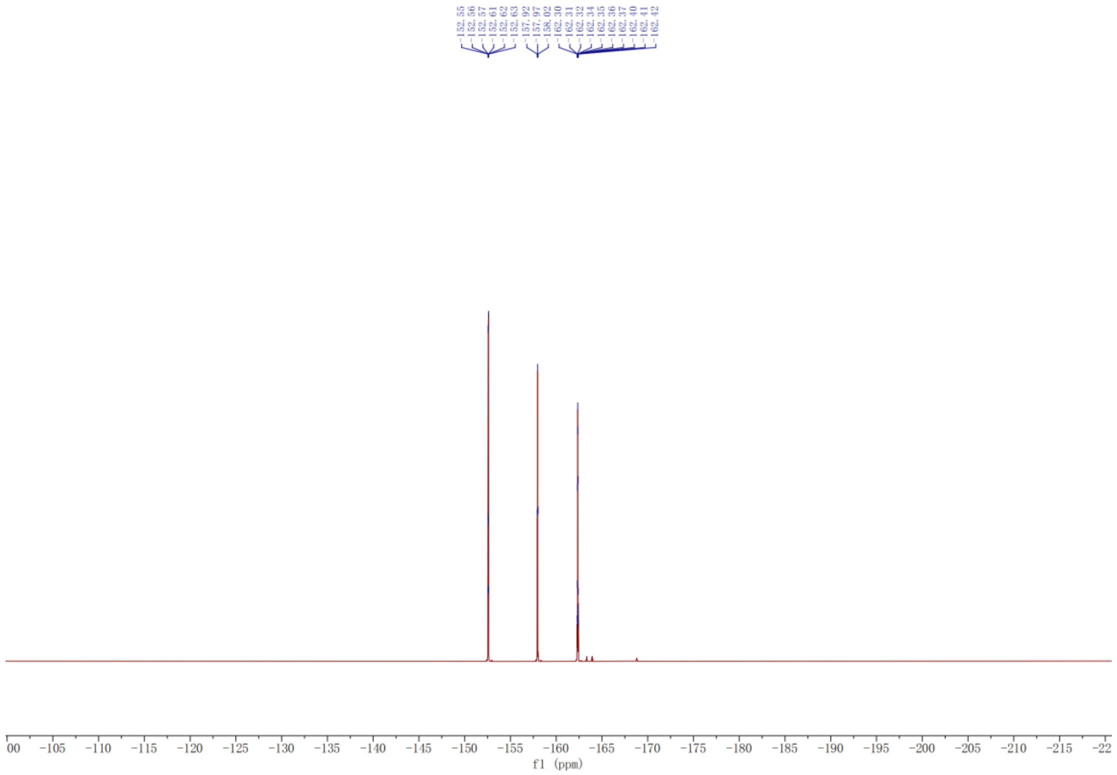
-152.55
-152.56
-152.57
-152.61
-152.62
-152.63
-157.92
-157.97
-158.02
-162.30
-162.31
-162.32
-162.34
-162.35
-162.36
-162.37
-162.40
-162.41
-162.42
f1 (ppm)

**$N_3$-3PEG-PFP (12)**

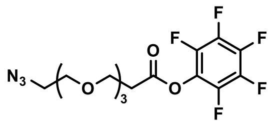

$N_3$-3PEG-PFP (12)

Colorless oil (70%).

$^1$H NMR (500 MHz, $CDCl_3$) δ 3.88 (t, $J$ = 6.2 Hz, 2H), 3.67 (m, 10H), 3.38 (t, $J$ = 5.1 Hz, 2H), 2.94 (t, $J$ = 6.2 Hz, 2H).

$^{13}$C NMR (126 MHz, $CDCl_3$) δ 167.57, 142.09, 140.09, 138.88, 136.92, 70.72, 70.70, 70.63, 70.05, 66.05, 50.69, 34.44.

$^{19}$F NMR (471 MHz, $CDCl_3$) δ -152.51 – -152.60 (m), -157.97 (t, $J$ = 21.6 Hz), -162.29 – -162.44 (m).

ESI-MS $[M+ Na]^+$ calcd for $C_{15}H_{16}F_5N_3O_5Na$: 436.0902, found: 436.0907.

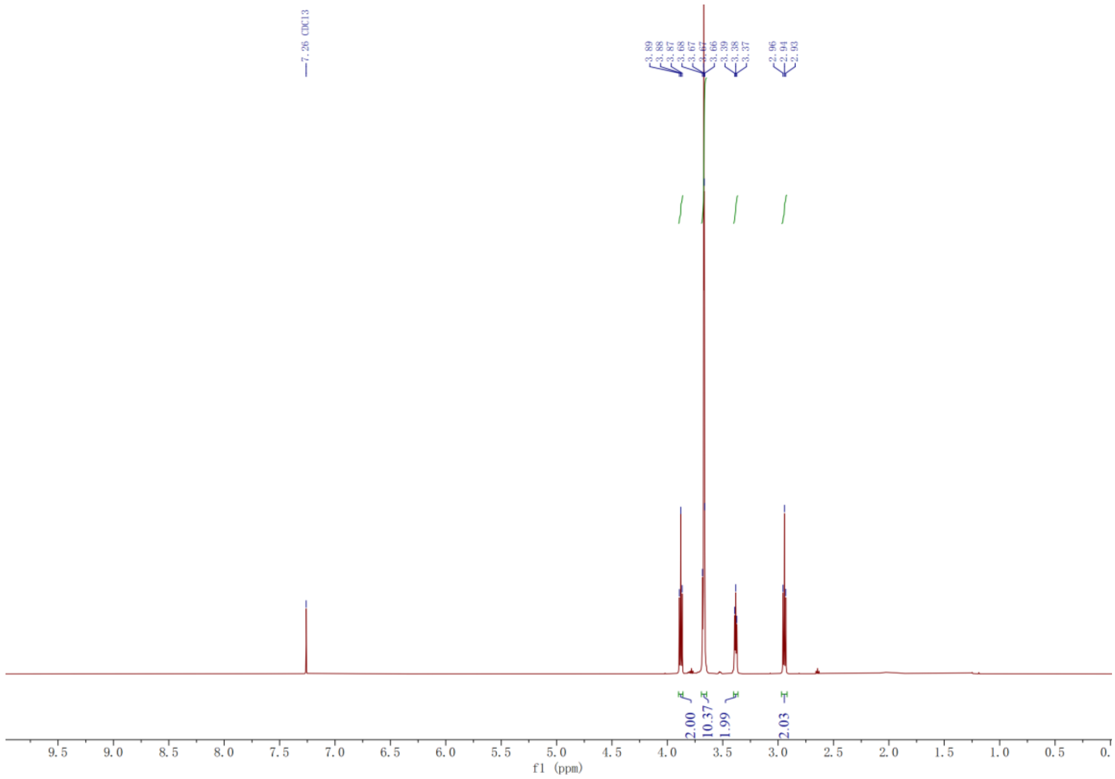
7.26 CDCl3
3.89
3.88
3.87
3.68
3.67
3.67
3.66
3.39
3.38
3.37
2.96
2.94
2.93
2.00
10.37
1.99
2.03
9.5 9.0 8.5 8.0 7.5 7.0 6.5 6.0 5.5 5.0 4.5 4.0 3.5 3.0 2.5 2.0 1.5 1.0 0.5 0.
f1 (ppm)

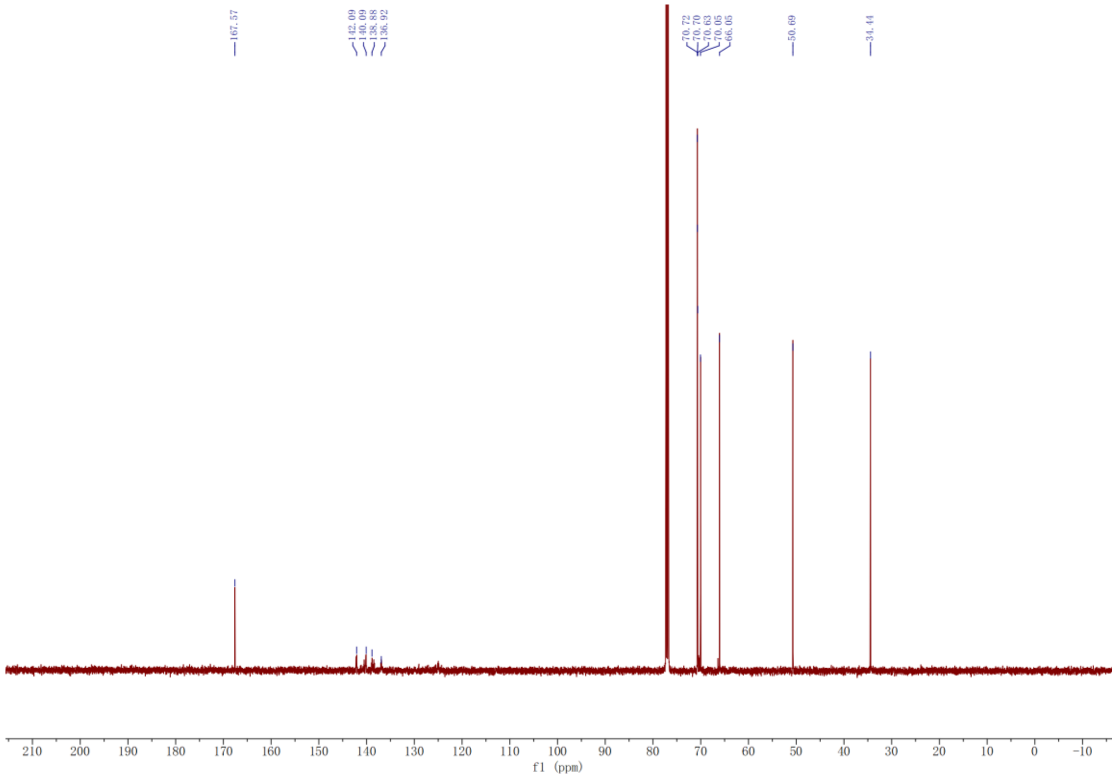
167.57
142.09
140.09
138.88
136.92
70.72
70.70
70.63
70.05
66.05
50.69
34.44
210 200 190 180 170 160 150 140 130 120 110 100 90 80 70 60 50 40 30 20 10 0 -10
f1 (ppm)

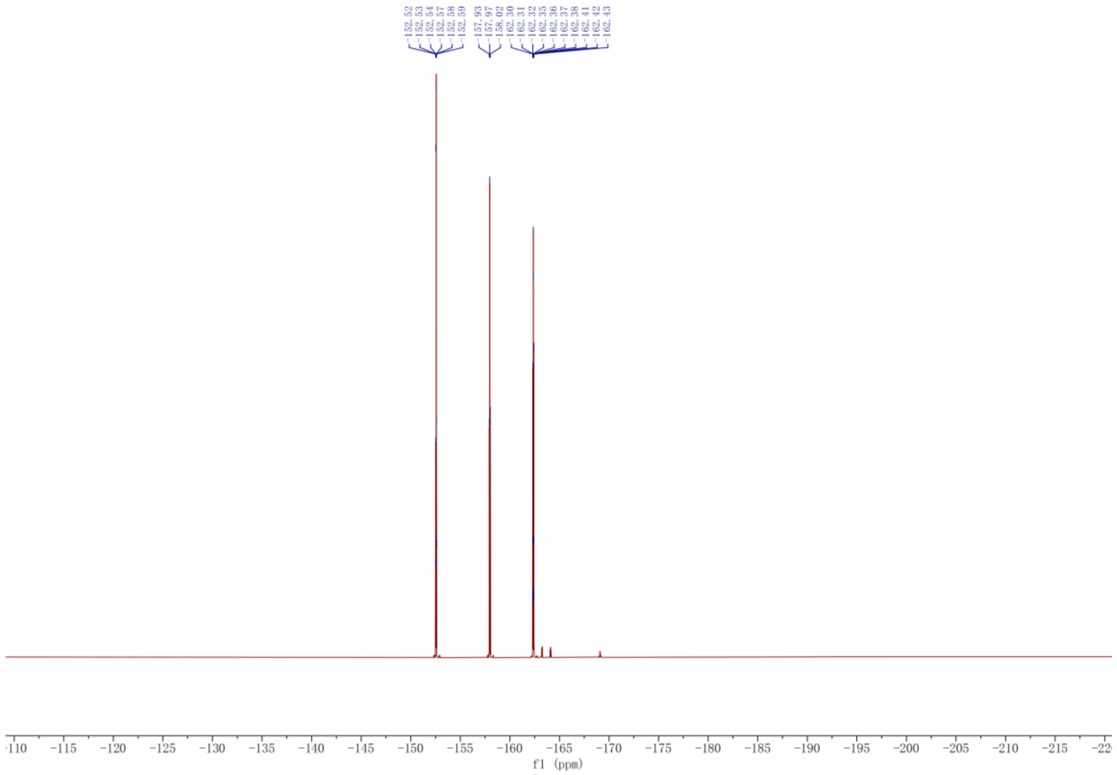
-152.52
-152.53
-152.54
-152.57
-152.58
-152.59
-157.93
-157.97
-158.02
-162.30
-162.31
-162.32
-162.35
-162.36
-162.37
-162.38
-162.41
-162.42
-162.43
-110
-115
-120
-125
-130
-135
-140
-145
-150
-155
-160
-165
-170
-175
-180
-185
-190
-195
-200
-205
-210
-215
f1 (ppm)

**$N_3$-4PEG-PFP (13)**

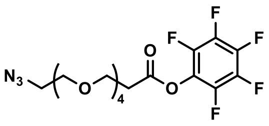

N3-4PEG-PFP (13)

Colorless oil (72%).

$^{1}$H NMR (500 MHz, $CDCl_3$) δ 3.88 (t, $J$ = 6.2 Hz, 2H), 3.67 (m, 14H), 3.38 (t, $J$ = 5.1 Hz, 2H), 2.94 (t, $J$ = 6.2 Hz, 2H).

$^{13}$C NMR (126 MHz, $CDCl_3$) δ 167.57, 141.93, 140.13, 138.58, 136.60, 70.72, 70.69, 70.67, 70.64, 70.53, 70.04, 66.05, 50.70, 34.44.

$^{19}$F NMR (471 MHz, $CDCl_3$) δ -152.50, -152.51, -152.52, -152.56, -152.57, -152.58, -157.92, -157.96, -158.01, -162.29, -162.30, -162.31, -162.34, -162.35, -162.36, -162.37, -162.40, -162.41, -162.42.

ESI-MS $[M+Na]^+$ calcd for $C_{17}H_{20}F_5N_3O_6Na$: 480.1164, found: 480.1173.

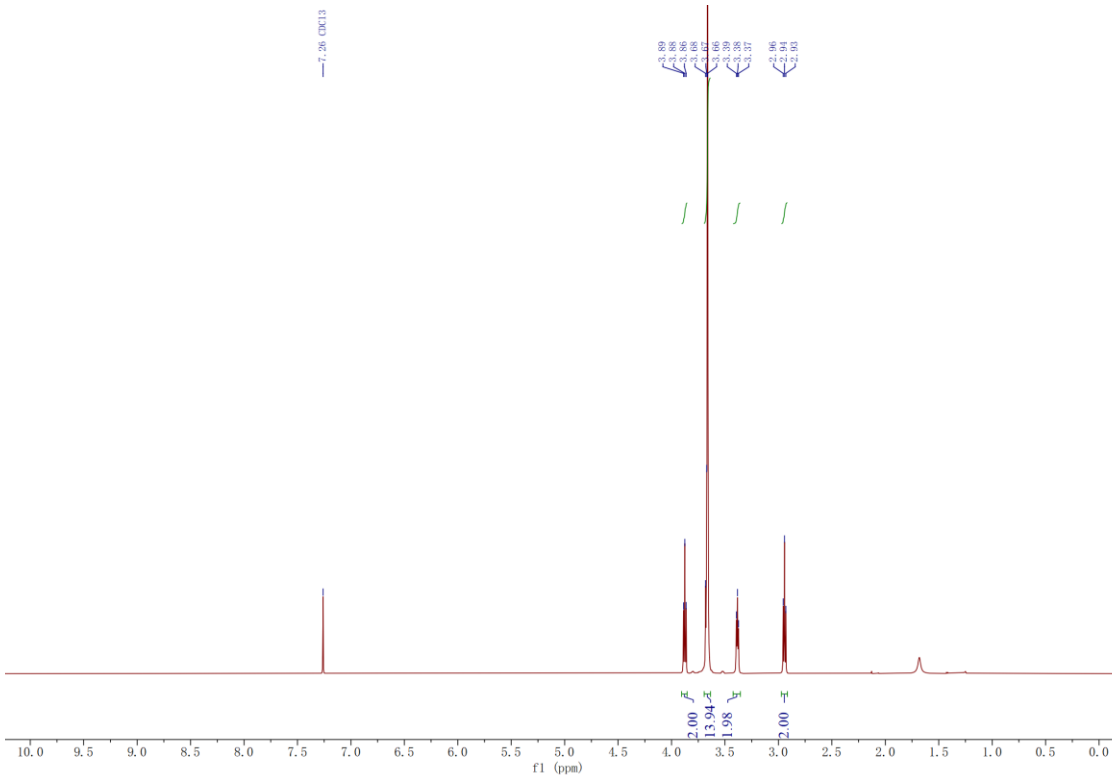

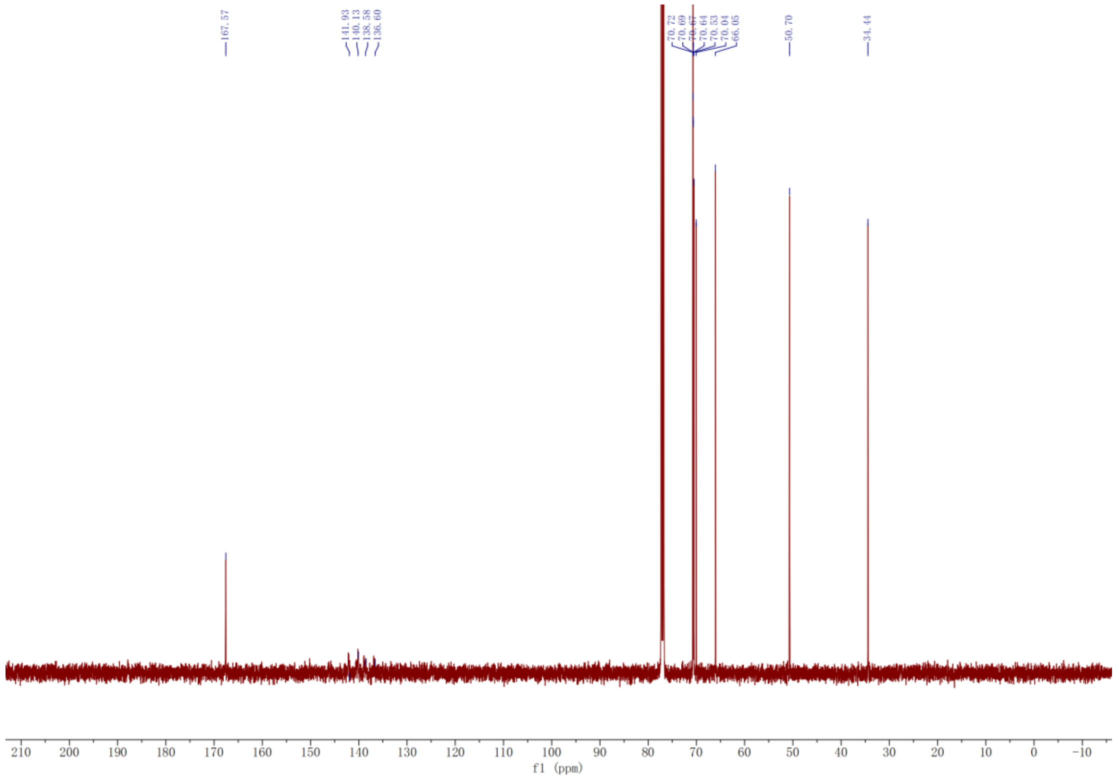
167.57
141.93
140.13
138.58
136.60
70.72
70.69
70.67
70.64
70.53
70.04
66.05
50.70
34.44
f1 (ppm)

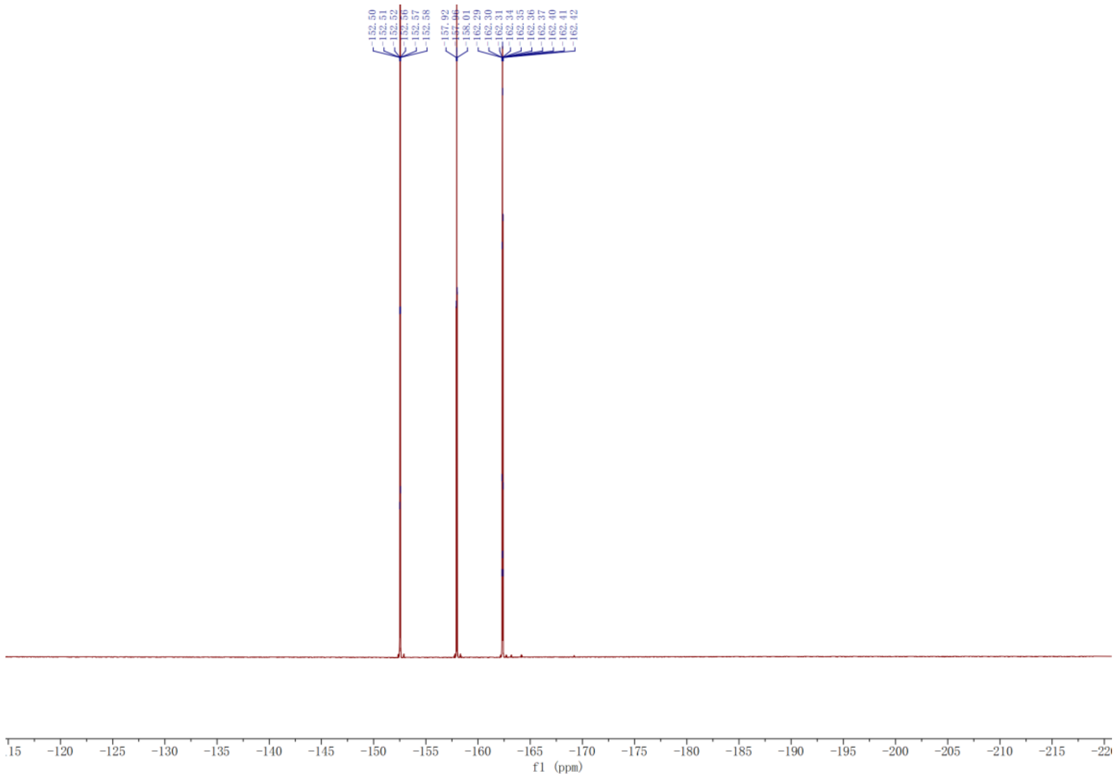
-152.50
-152.51
-152.52
-152.56
-152.57
-152.58
-157.92
-157.96
-158.01
-162.29
-162.30
-162.31
-162.34
-162.35
-162.36
-162.37
-162.40
-162.41
-162.42
f1 (ppm)

#### Synthesis of acylating reagents NAN$_3$-STP (14):

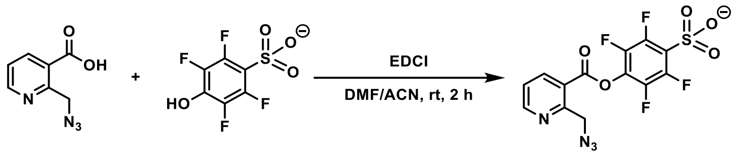


2a 14a NAN$_3$-STP (14)

To a solution of NAN$_3$-COOH (2a, 140 mg, 0.75 mmol), 2,3,5,6-tetrafluoro-4-hydroxy-benzenesulfonate[9] (14a, 200 mg, 0.75 mmol) in ACN (4.0 mL) and DMF (1.0 mL) was added EDCI (120 mg. 0.83 mmol). The solution was stirred at room temperature for 2 h, then purified by preparative reversed-phase HPLC (0% to 100% ACN in water over 50 min). The product-containing fractions were dried by lyophilization to afford **NAN$_3$-STP** (**14**) as a white solid (45 mg, 15%).

$^1$H NMR (500 MHz, DMSO) δ 8.95 (dd, $J$ = 4.8, 1.7 Hz, 1H), 8.64 (dd, $J$ = 7.9, 1.8 Hz, 1H), 7.69 (dd, $J$ = 7.9, 4.8 Hz, 1H), 4.88 (s, 2H).

$^{13}$C NMR (151 MHz, DMSO) δ 168.83, 154.37, 148.69, 145.50, 143.90, 141.34, 139.78, 138.17, 123.05, 53.45.

$^{19}$F NMR (471 MHz, DMSO) δ -141.62 – -141.75 (m), -162.19 – -162.32 (m).

ESI-MS [M]$^-$ calcd for $C_{13}H_5F_4N_4O_5S$: 404.9922, found: 404.9913.

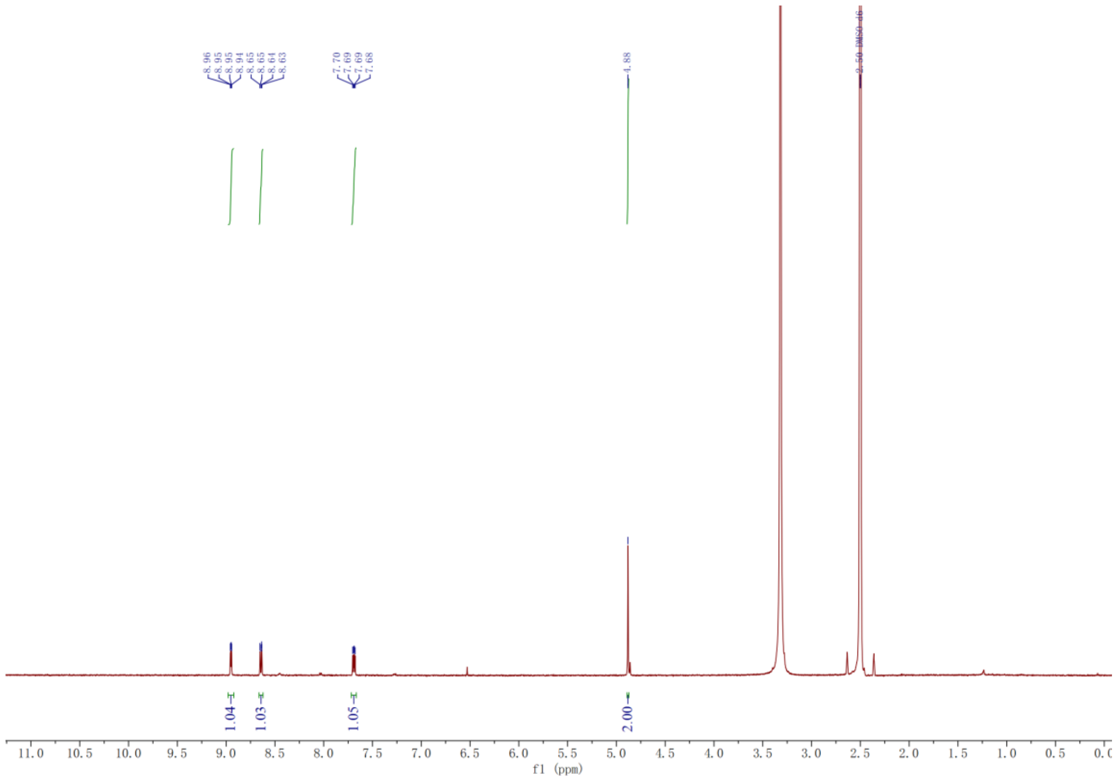
8.96
8.95
8.95
8.94
8.65
8.65
8.64
8.63
7.70
7.69
7.69
7.68
4.88
2.50 DMSO-d6
1.04
1.03
1.05
2.00
f1 (ppm)

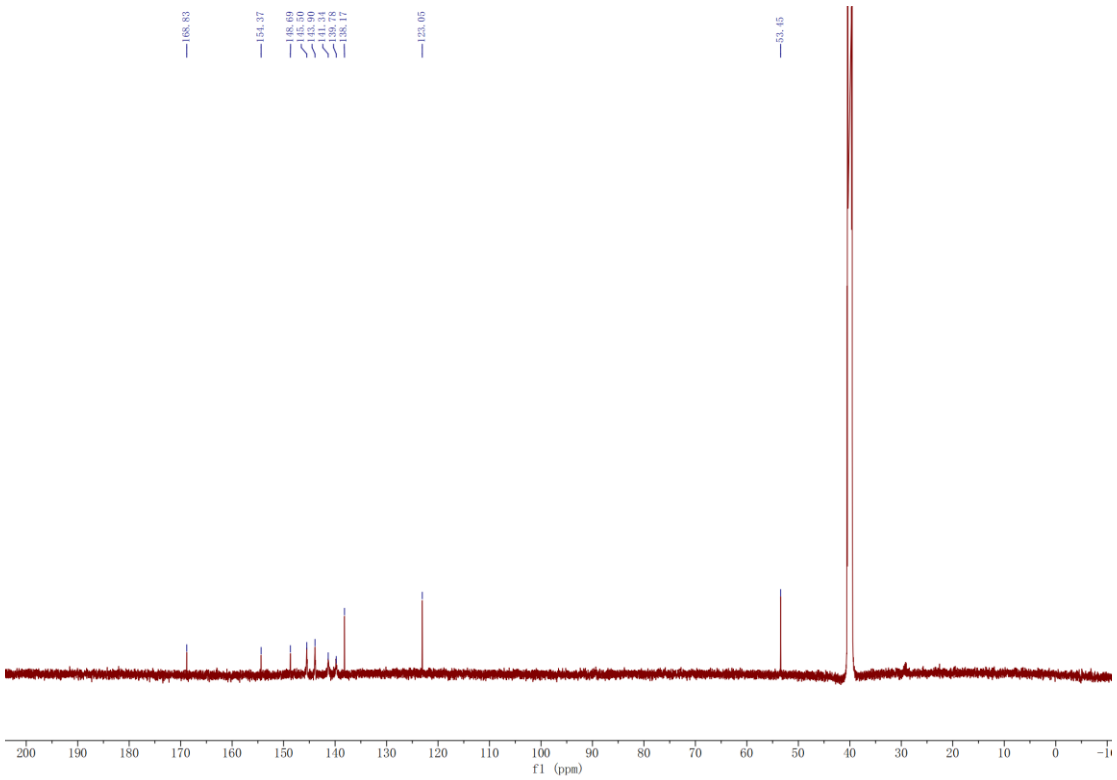
168.83
154.37
148.69
145.50
143.90
141.34
139.78
138.17
123.05
53.45
f1 (ppm)

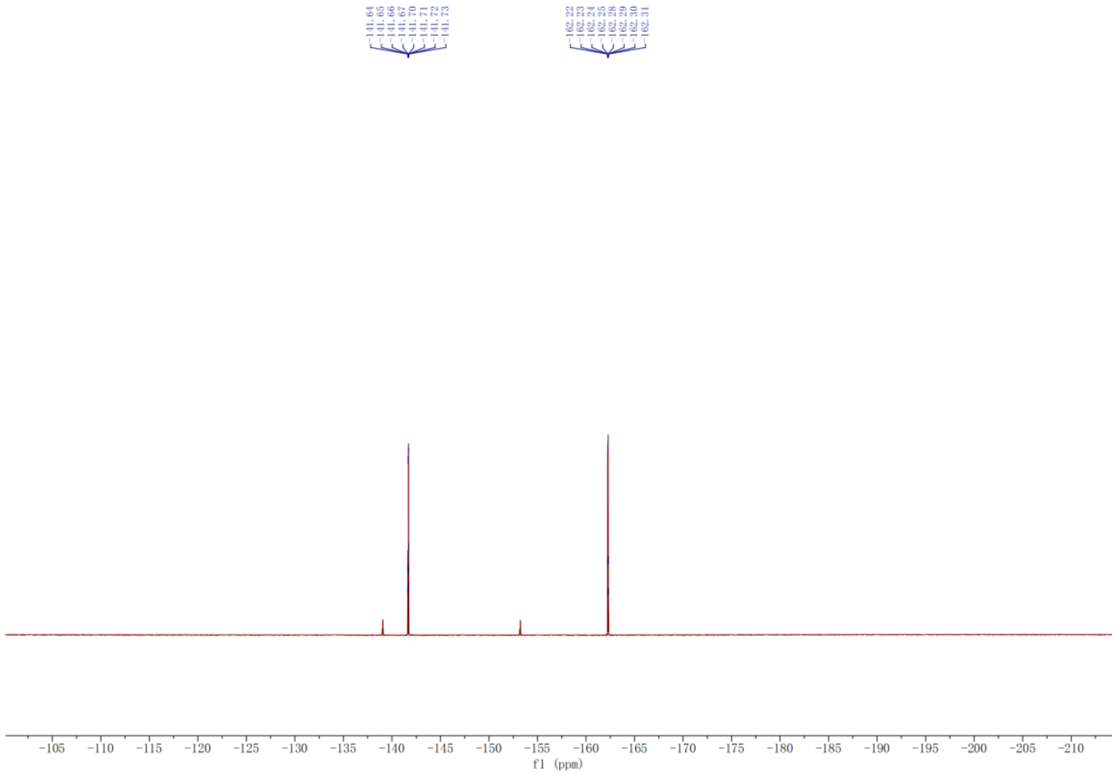
-141.64
-141.65
-141.66
-141.67
-141.70
-141.71
-141.72
-141.73
-162.22
-162.23
-162.24
-162.25
-162.28
-162.29
-162.30
-162.31
-105
-110
-115
-120
-125
-130
-135
-140
-145
-150
-155
-160
-165
-170
-175
-180
-185
-190
-195
-200
-205
-210
f1 (ppm)

**Synthesis of acylating reagents N3-2PEG-NASA (15):**

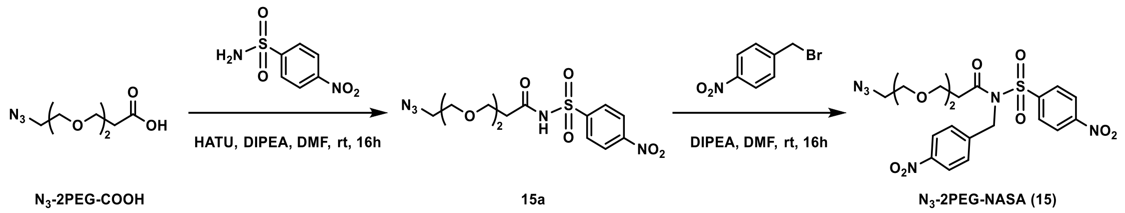


To a solution of $N_3$-2PEG-COOH (220 mg, 1.10 mmol), 4-nitrobenzenesulfonamide (200 mg, 1.00 mmol) in DMF (4.0 mL) was added HATU (570 mg. 1.5 mmol) and DIPEA (258mg, 2.0 mmol). The solution was stirred at room temperature for 16 h, then purified by preparative thin-layer chromatography (Hexane:EA = 5:1) to afford the intermediate **15a**. After dissolved in DMF (5.0 mL), 1-(bromomethyl)-4-nitrobenzene (648mg, 3.0 mmol) and DIPEA (387mg, 3.0 mmol) were added into the reaction. The solution was stirred at room temperature for 16 h, then purified by preparative thin-layer chromatography (Hexane:EA = 2:1) to afford the **$N_3$-2PEG-NASA** (**15**) as a light-yellow solid (110 mg, 22%).

$^1$H NMR (600 MHz, DMSO) δ 8.45 (d, $J$ = 8.9 Hz, 2H), 8.27 (dd, $J$ = 16.4, 8.8 Hz, 4H), 7.61 (d, $J$ = 8.7 Hz, 2H), 5.31 (s, 2H), 3.56 – 3.49 (m, 4H), 3.45 (dd, $J$ = 5.9, 3.5 Hz, 2H), 3.39 (dd, $J$ = 5.8, 3.5 Hz, 2H), 3.33 (d, $J$ = 5.1 Hz, 2H), 2.78 (t, $J$ = 6.1 Hz, 2H).

$^{13}$C NMR (151 MHz, DMSO) δ 171.98, 151.01, 147.38, 145.17, 144.35, 130.16, 128.09, 125.03, 124.37, 70.16, 69.92, 69.64, 65.85, 50.39, 49.70, 36.19.

ESI-MS [M+Na]$^+$ calcd for $C_{20}H_{22}N_6O_9SNa$: 545.1061, found: 545.1064.

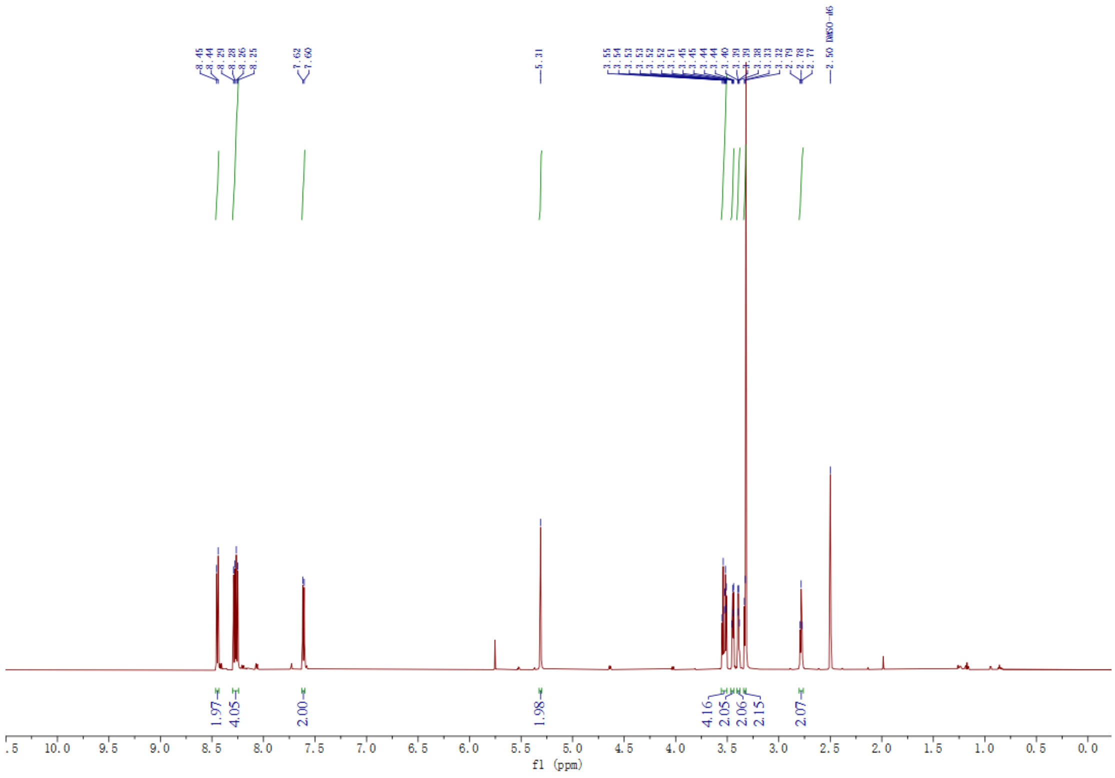

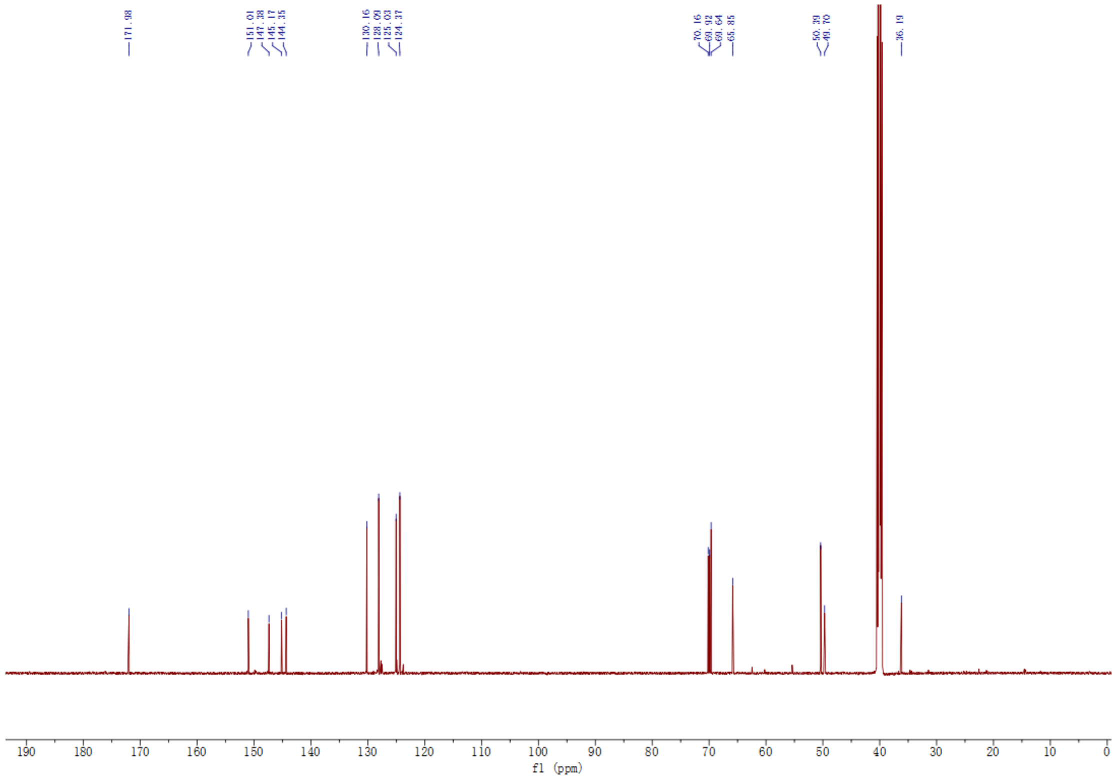

**Synthesis of NBSI-DMAP (16):**

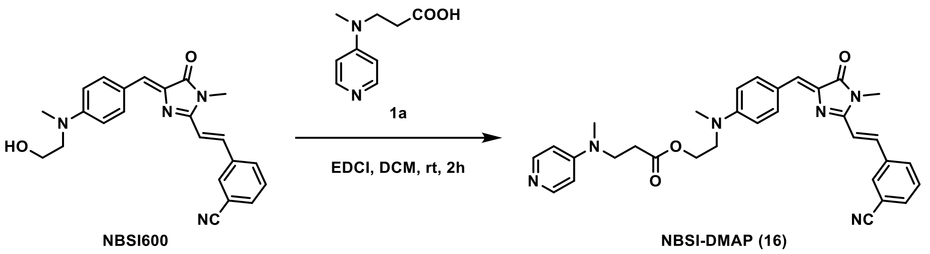


The synthetic procedure for **NBSI-DMAP** (**16)** was identical to that for **HBC-DMAP** (**1**), except HBC530 was replaced with NBSI600[3]. Red solid (50%).

$^1$H NMR (500 MHz, DMSO) δ 8.42 (s, 1H), 8.18 (d, *J* = 8.3 Hz, 2H), 8.17 – 8.11 (m, 3H), 7.93 (d, *J* = 15.9 Hz, 1H), 7.85 (d, *J* = 7.8 Hz, 1H), 7.66 (t, *J* = 7.8 Hz, 1H), 7.40 (d, *J* = 15.9 Hz, 1H), 6.98 (s, 1H), 6.87 – 6.79 (m, 4H), 4.25 (t, *J* = 5.6 Hz, 2H), 3.72 (t, *J* = 5.6 Hz, 2H), 3.68 (t, *J* = 7.1 Hz, 2H), 3.59 (q, *J* = 6.5 Hz, 2H), 3.28 (s, 3H), 3.02 (s, 3H), 3.00 (s, 3H), 2.56 (t, *J* = 7.1 Hz, 2H).

$^{13}$C NMR (126 MHz, DMSO) δ 171.46, 170.11, 165.03, 157.16, 150.94, 137.17, 136.45, 135.84, 134.95, 133.24, 133.11, 131.78, 127.95, 126.51, 122.75, 119.04, 117.34, 112.60, 112.27, 107.50, 61.99, 53.81, 47.31, 42.10, 31.34, 26.90.

ESI-MS $[M+H]^+$ calcd for $C_{32}H_{33}N_6O_3$: 549.25, found: 549.2609.

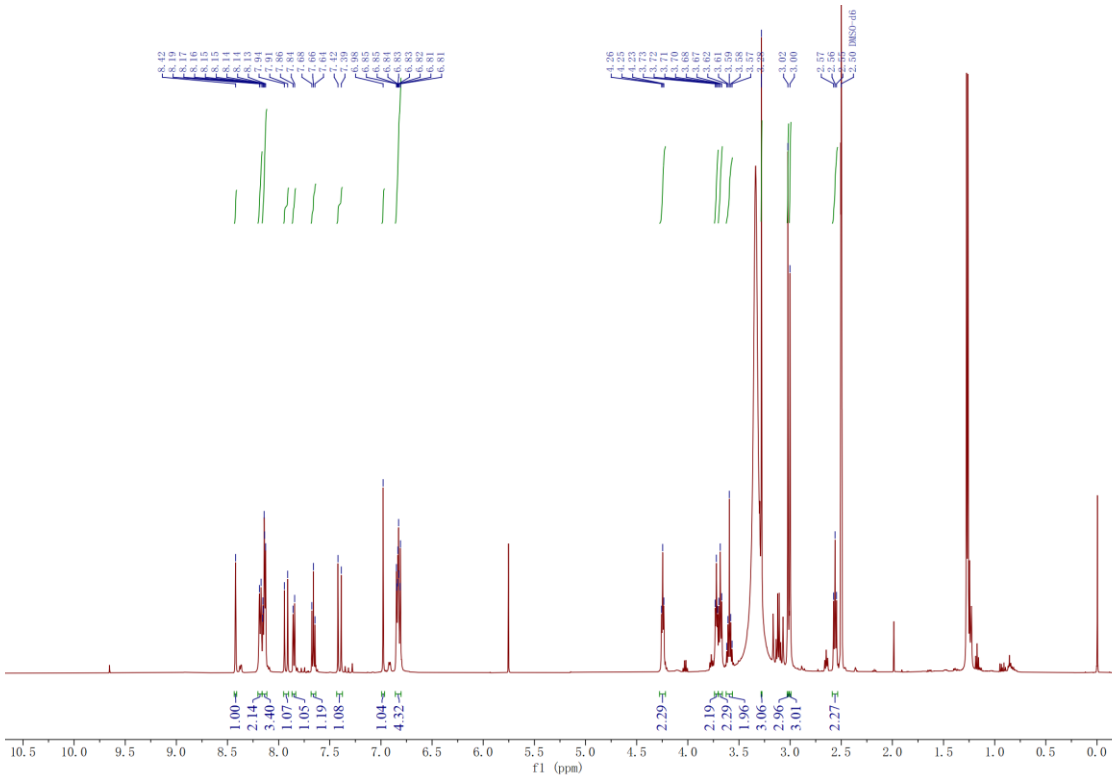

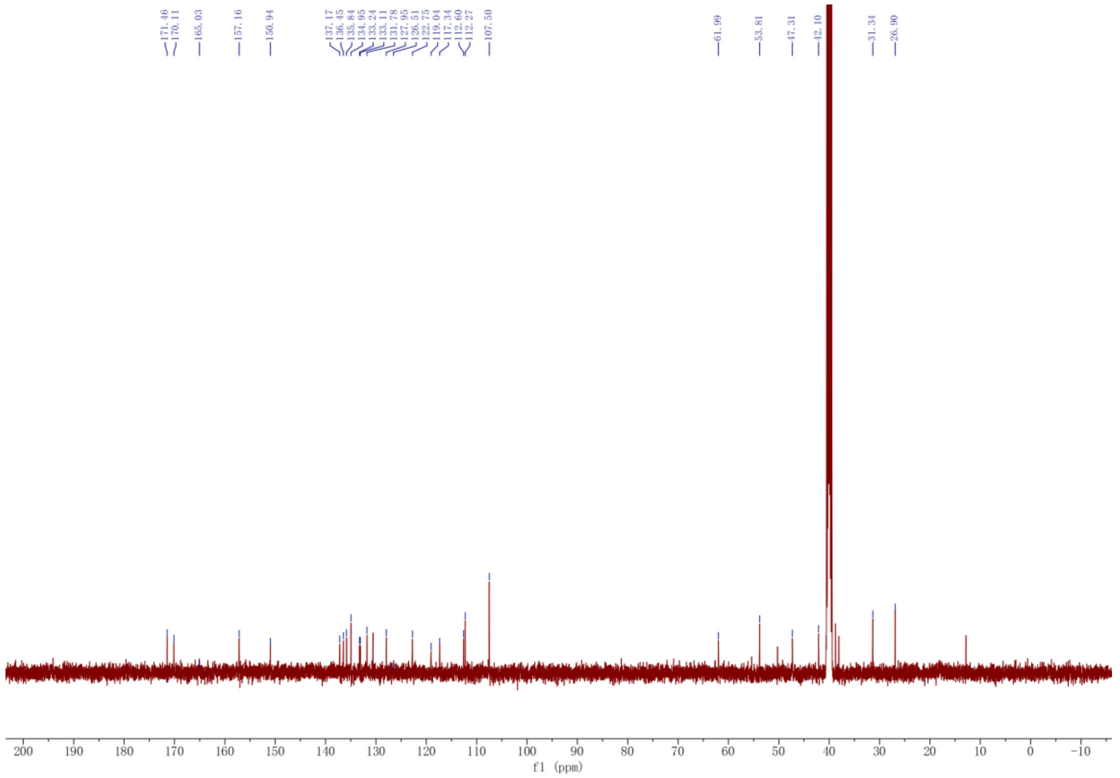

**Synthesis of DFHBI-DMAP (17):**

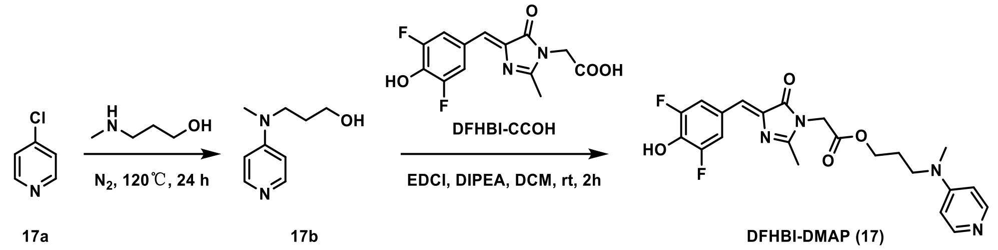


To a solution of 4-chloropyridine (17a, 113 mg, 1.0 mmol) was added 3-(methylamino) propan-1-ol (4 mL). The solution was stirred at 120 °C for 16 h, then purified by preparative reversed-phase HPLC (0% to 100% ACN in water over 35 min). The product-containing fractions were dried by lyophilization to afford intermediate **17b**. After dissolved in DCM (5.0 mL), DFHBI-COOH[10] (296 mg, 1.0 mmol), DIPEA (258 mg, 2.0 mmol) and EDCI (225 mg, 1.5 mmol) were added into the reaction. The solution was stirred at room temperature for 2 h, then purified by preparative reversed-phase HPLC (0% to 100% ACN in water over 35 min). The product-containing fractions were dried by lyophilization to afford the **DFHBI-DMAP** (**17**) as a white solid (130 mg, 30%).

$^1$H NMR (600 MHz, DMSO) δ 11.05 (s, 1H), 8.21 (d, *J* = 5.6 Hz, 1H), 7.99 (d, *J* = 9.9 Hz, 2H), 7.08 (s, 2H), 6.99 (s, 1H), 4.56 (s, 2H), 4.19 (t, *J* = 6.1 Hz, 2H), 3.65 (t, *J* = 7.2 Hz, 2H), 3.15 (s, 3H), 2.34 (s, 3H), 1.97 – 1.90 (m, 2H)..

$^{13}$C NMR (151 MHz, DMSO) δ 169.69, 168.48, 163.60, 158.43, 158.21, 157.08, 153.13, 151.53, 137.95, 136.54, 124.52, 115.92, 115.77, 63.16, 48.69, 38.58, 25.68, 15.71.

ESI-MS [M+H]$^+$ calcd for $C_{22}H_{23}F_2N_4O_4$: 445.1682, found: 445.1685.

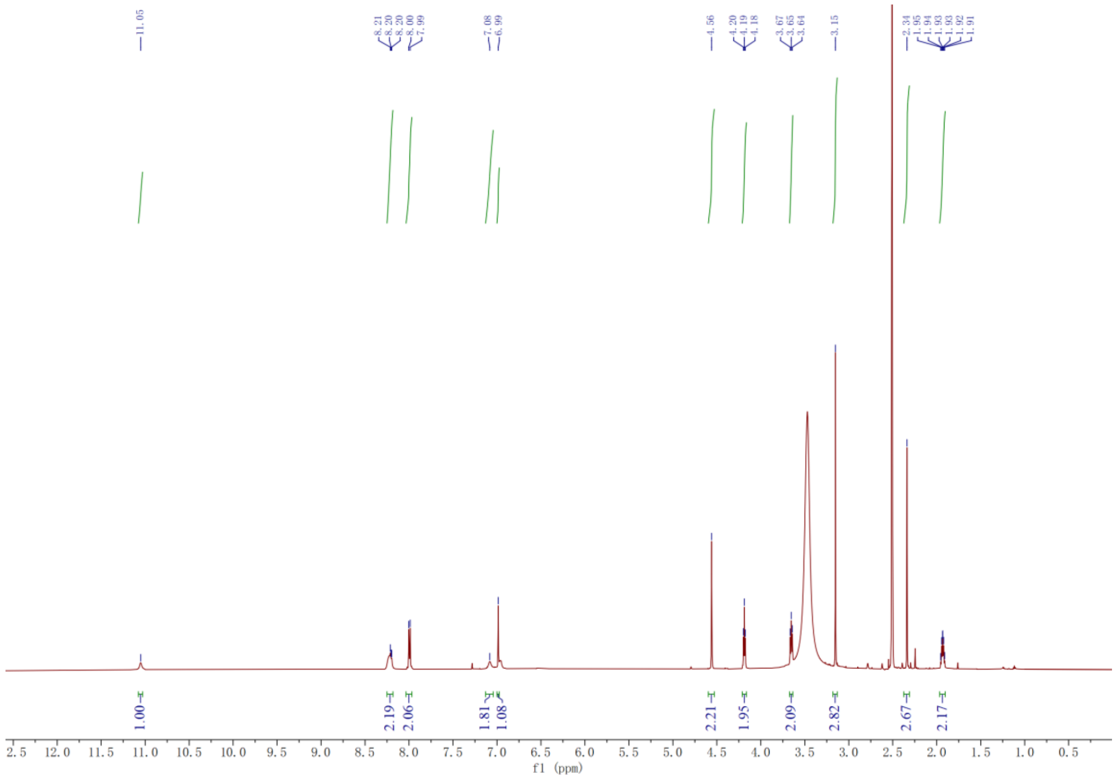
11.05
8.21 8.20 8.20 8.00 7.99
7.08 6.99
4.56
4.20 4.19 4.18
3.67 3.65 3.64
3.15
2.34 1.95 1.94 1.93 1.93 1.92 1.91
1.00 2.19 2.06 1.81 1.08 2.21 1.95 2.09 2.82 2.67 2.17
f1 (ppm)

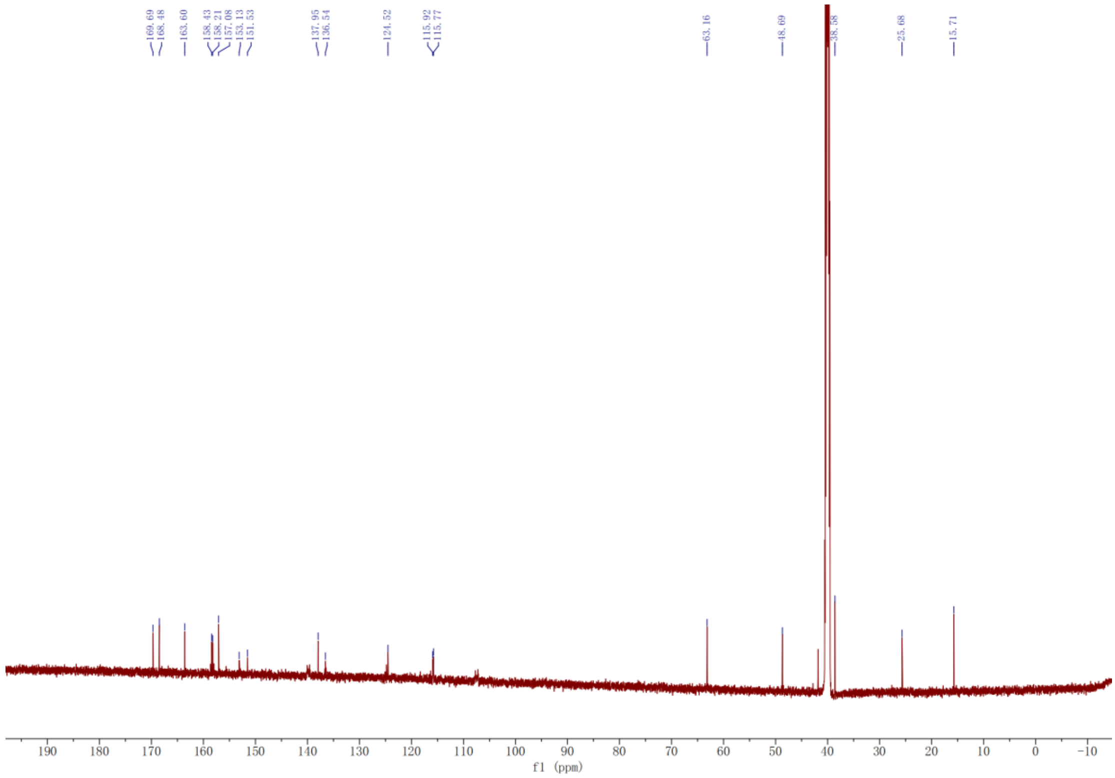
169.69 168.48
163.60
158.43 158.21 157.08 153.13 151.53
137.95 136.54
124.52
115.92 115.77
63.16
48.69
25.68
15.71
f1 (ppm)

**Synthesis of FMN-DMAP (18):**

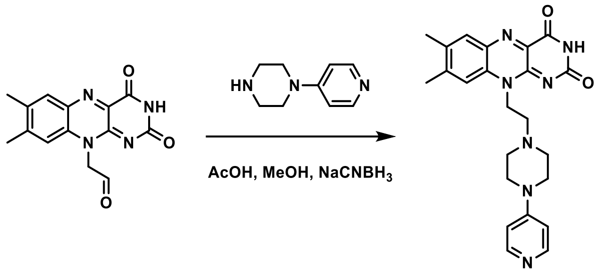


18a    FMN-DMAP (18)

To a solution of 18a[4] (200 mg, 0.70 mmol) in EtOH (10 mL) was added 1-(pyridin-4-yl)piperazine (342 mg, 2.1 mmol). The solution was stirred at 40°C for 2 h, then the $NaCNBH_3$ (90mg, 1.4 mmol) and drops of AcOH were added into the reaction to stir at 40°C for 24 h. The reaction was purified by filtration and washing by MeOH to afford **FMN-DMAP** (**18**) as a yellow solid (90mg, 20%).

$^1$H NMR (500 MHz, DMSO) δ 11.33 (s, 1H), 8.22 (d, *J* = 6.7 Hz, 2H), 7.90 (s, 1H), 7.82 (s, 1H), 7.16 (d, *J* = 6.9 Hz, 2H), 4.79 (t, *J* = 6.7 Hz, 2H), 3.58 (t, *J* = 5.0 Hz, 4H), 2.79 (t, *J* = 6.7 Hz, 2H), 2.70 (t, *J* = 5.0 Hz, 4H), 2.52 (s, 3H), 2.41 (s, 3H).

$^{13}$C NMR (126 MHz, DMSO) δ 160.46, 156.77, 156.04, 150.74, 146.88, 141.72, 137.56, 136.27, 134.22, 131.45, 116.85, 108.08, 54.31, 52.75, 46.23, 42.28, 21.15, 19.25.

ESI-MS $[M+H]^+$ calcd for $C_{23}H_{26}N_7O_2$: 432.2142, found: 432.2141

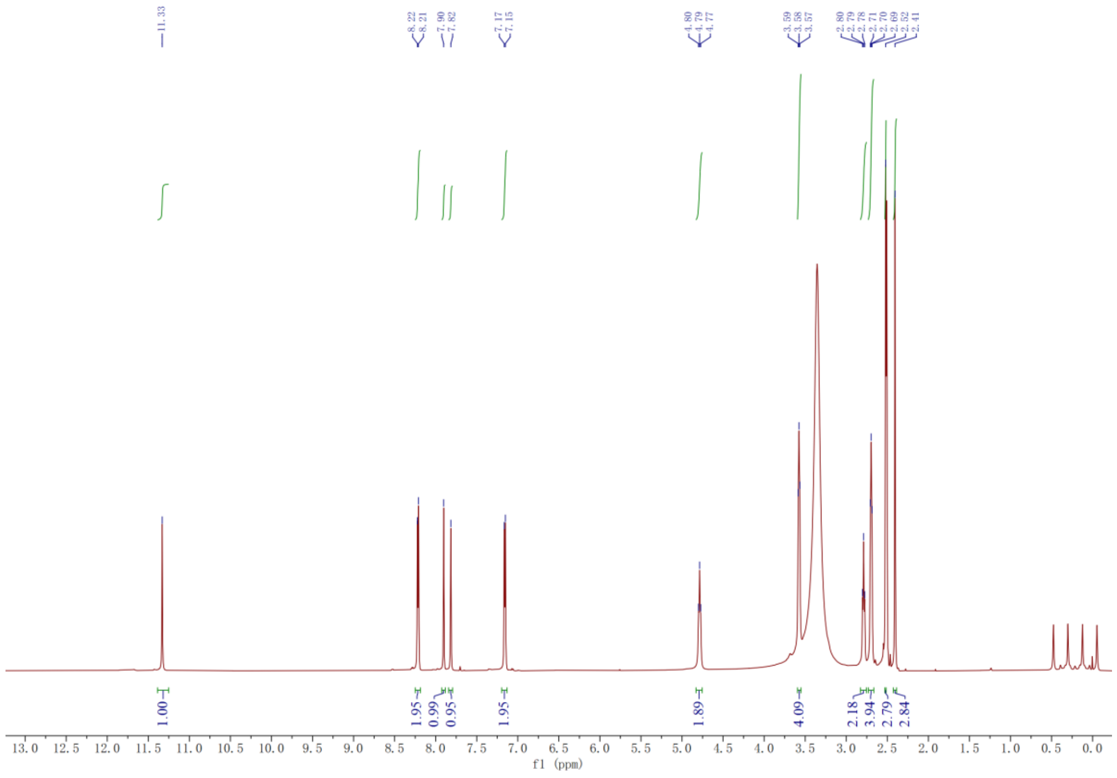
11.33
8.22
8.21
7.90
7.82
7.17
7.15
4.80
4.79
4.77
3.59
3.58
3.57
2.80
2.79
2.78
2.71
2.70
2.69
2.52
2.41
1.00
1.95
0.99
0.95
1.95
1.89
4.09
2.18
3.94
2.79
2.84
f1 (ppm)

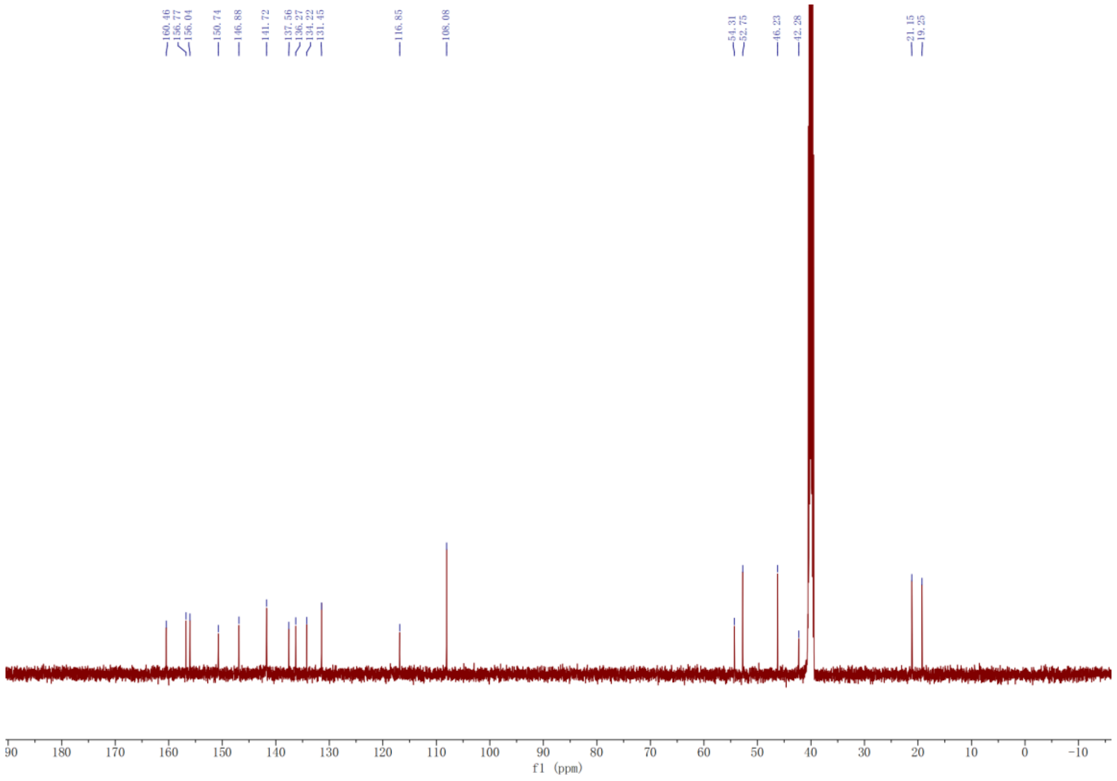
160.46
156.77
156.04
150.74
146.88
141.72
137.56
136.27
134.22
131.45
116.85
108.08
54.31
52.75
46.23
42.28
21.15
19.25
f1 (ppm)

#### MS Spectra:

#### HBC-DMAP (1)

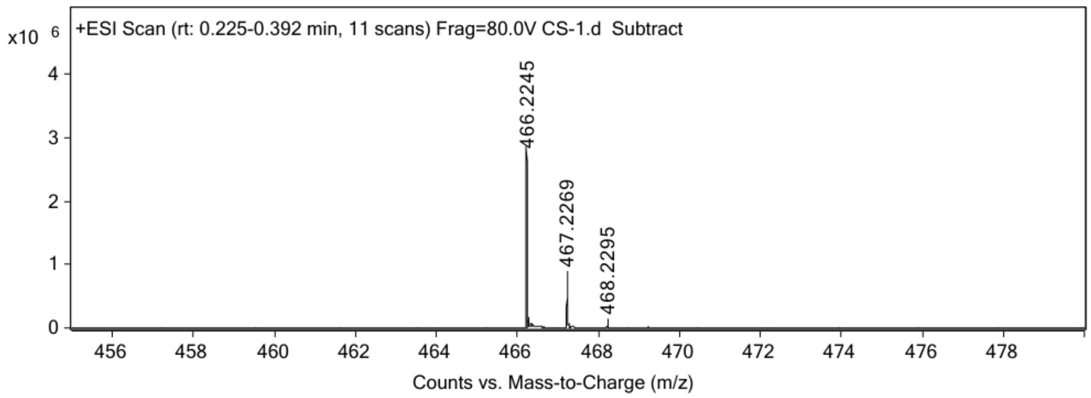


#### NAN$_3$-PFP (2)

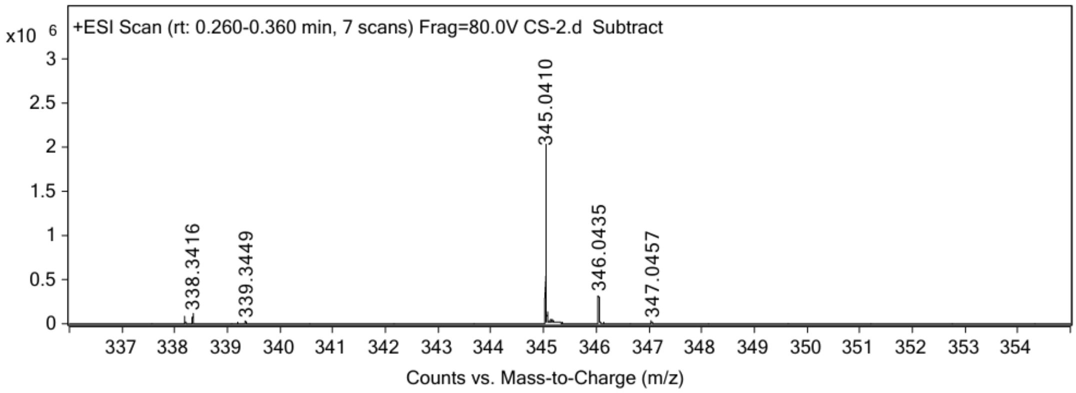


#### HBC-DMAP analogue (3)

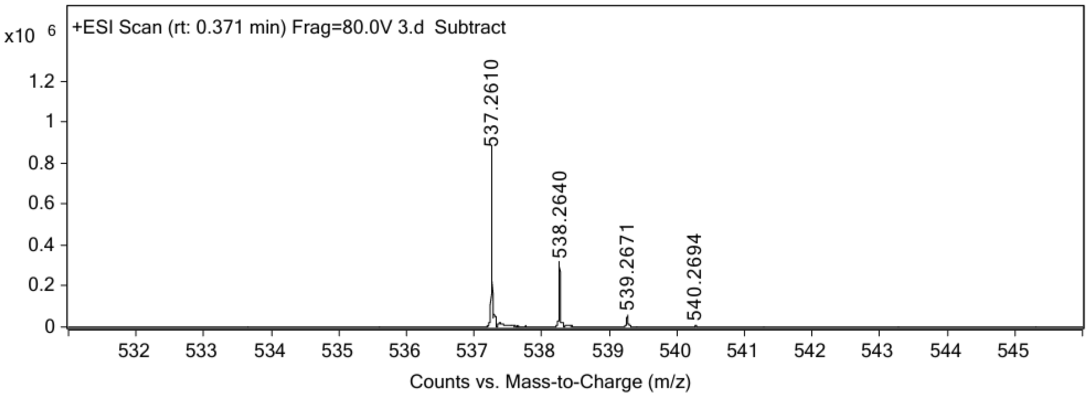

#### HBC-DMAP analogue (4)

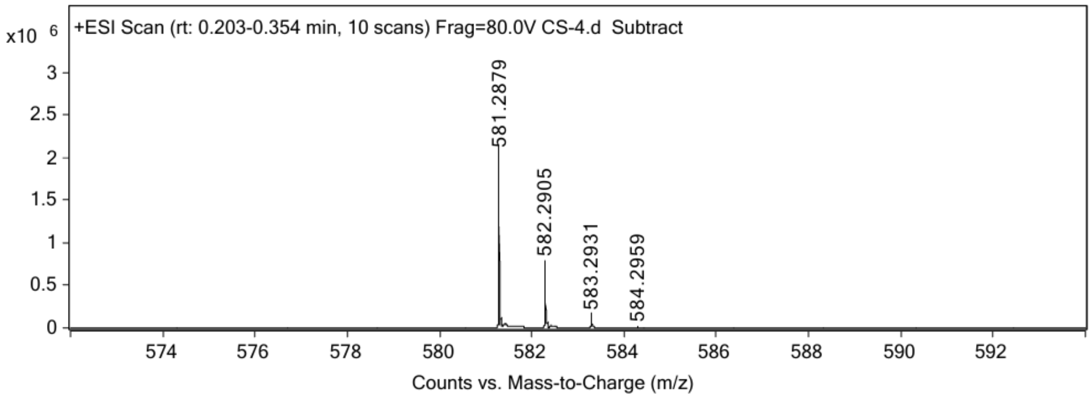


#### HBC-DMAP analogue (5)

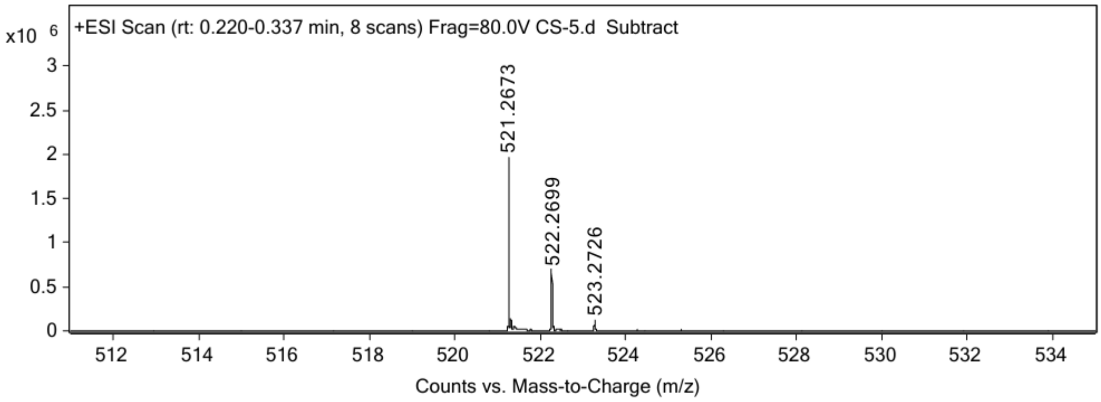


#### HBC-PyOx (6)

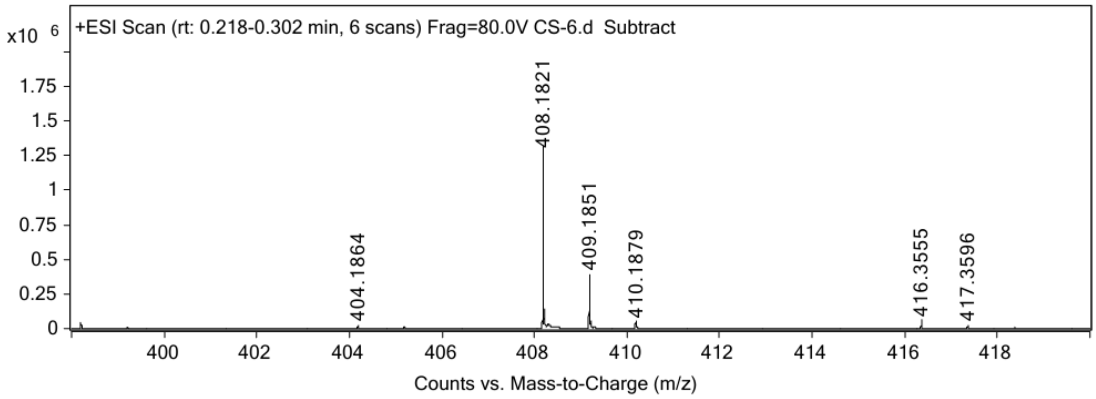

#### HBC-PyOx analogue (7)

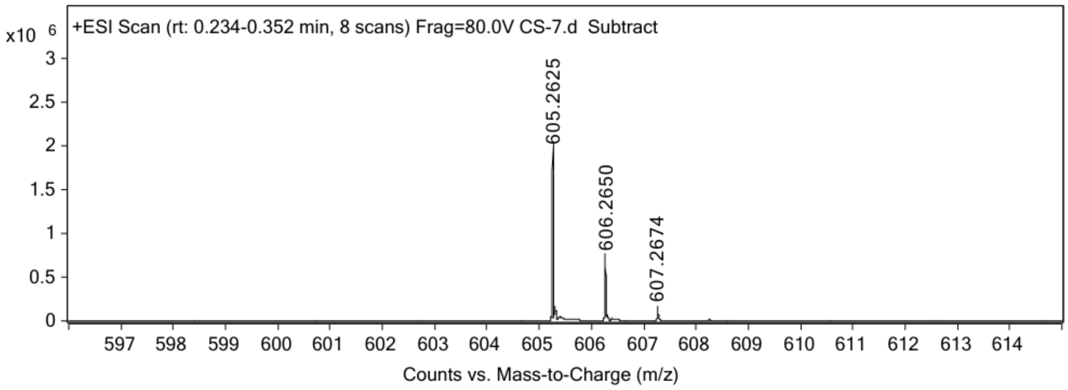


#### HBC-PyOx analogue (8)

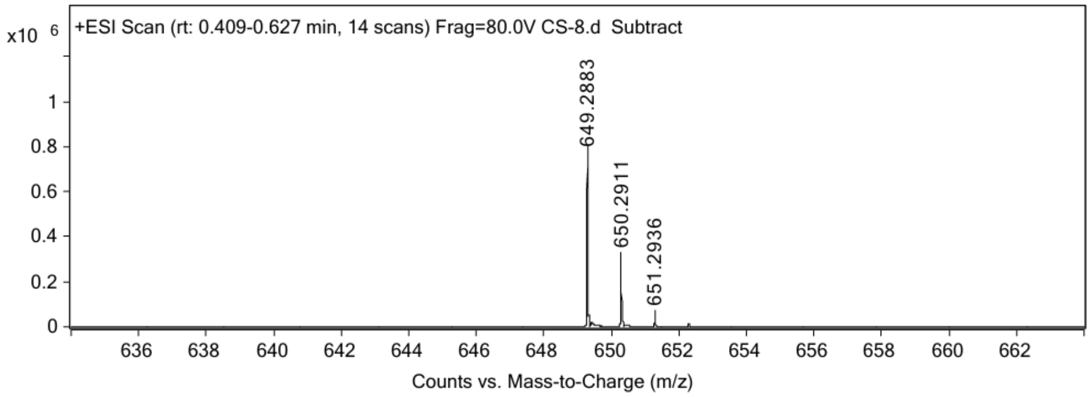


#### HBC-PyOx analogue (9)

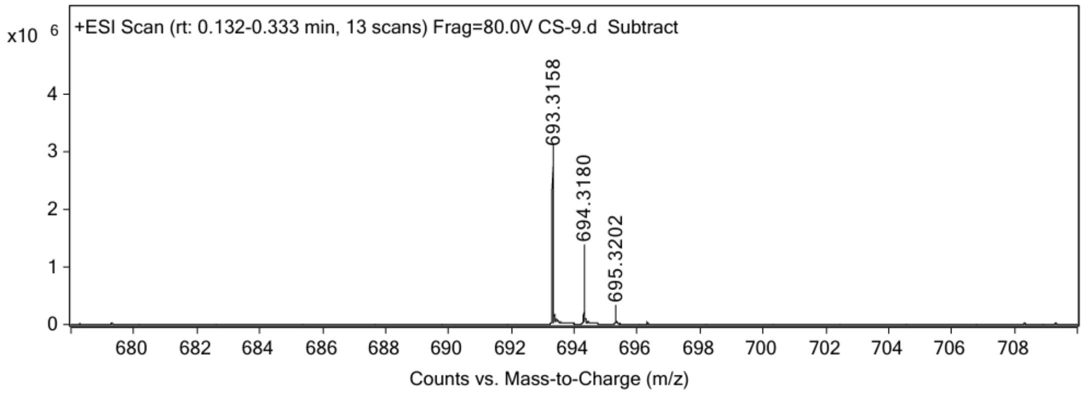

#### HBC-Imidazole (10)

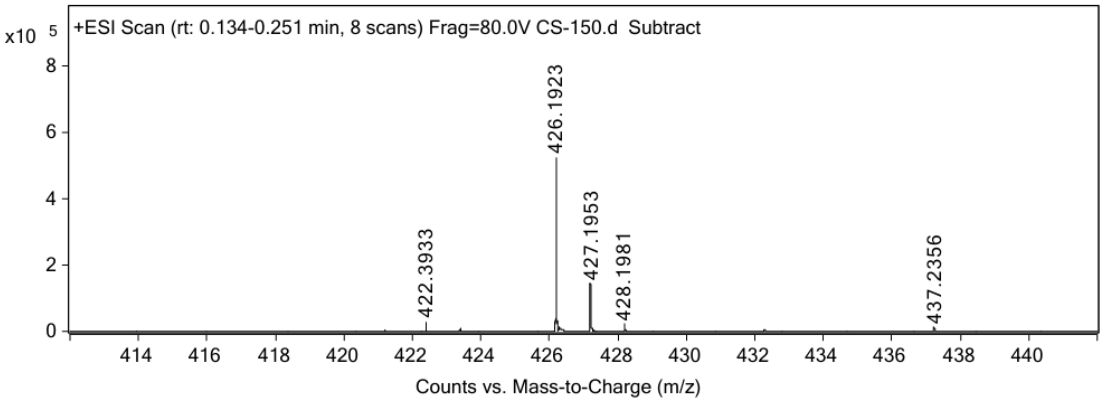


#### $N_3$-2PEG-PFP (11)

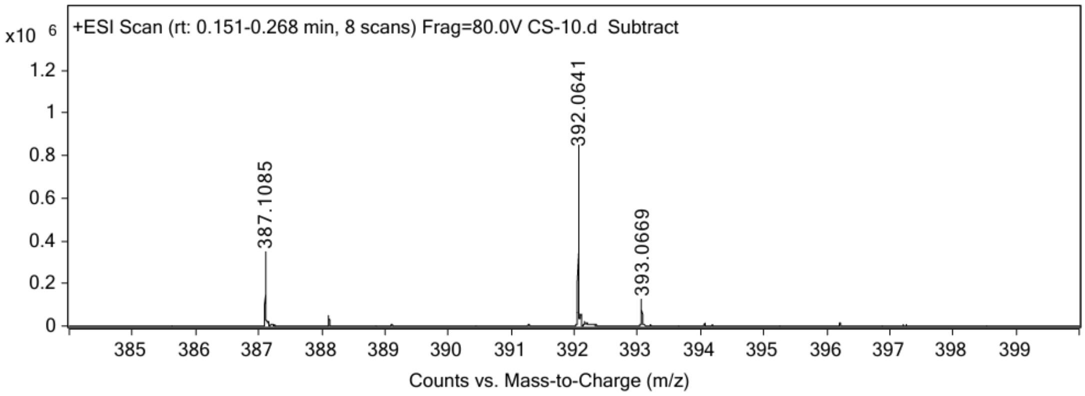


#### $N_3$-3PEG-PFP (12)

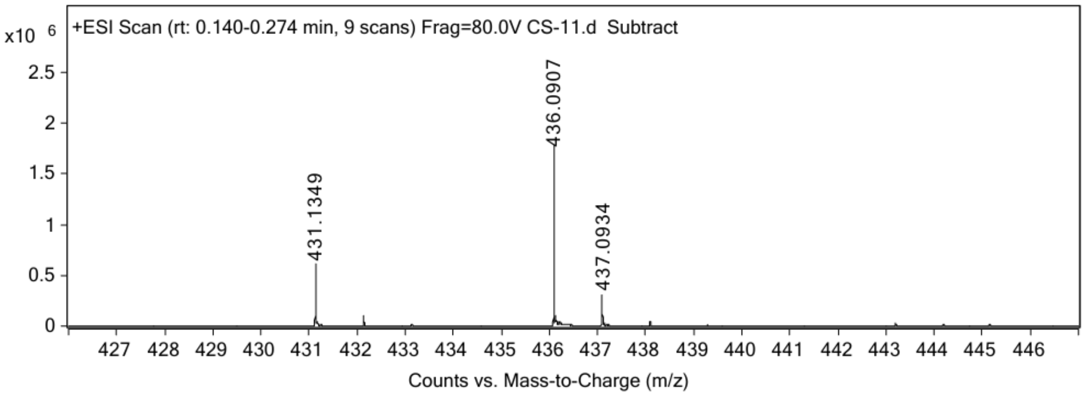

**$N_3$-4PEG-PFP (13)**

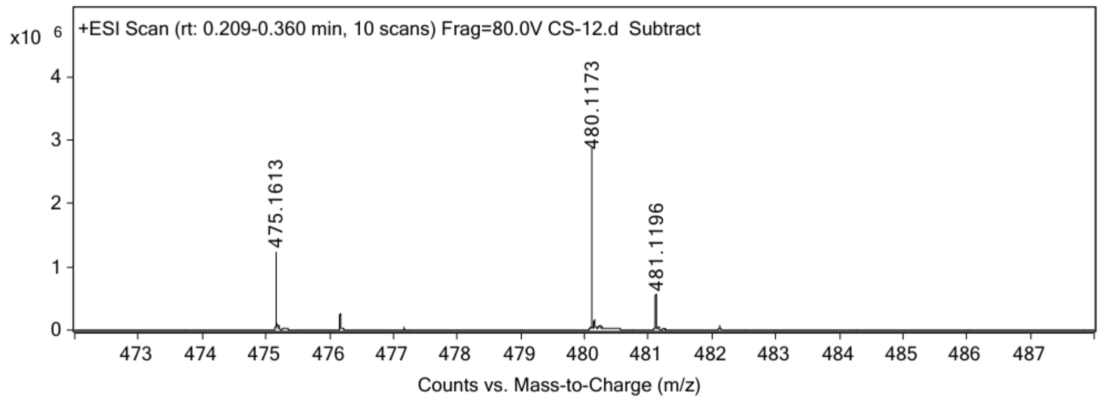


**$NAN_3$-STP (14)**

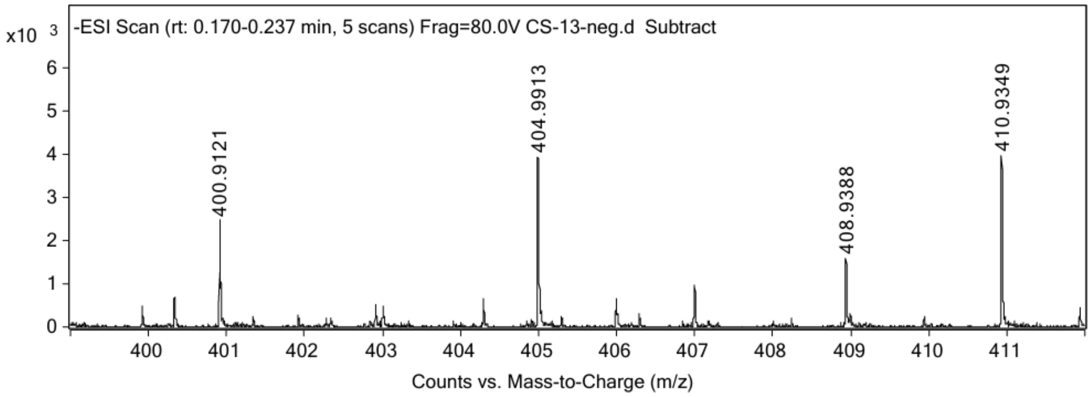


**$N_3$-2PEG-NASA (15)**

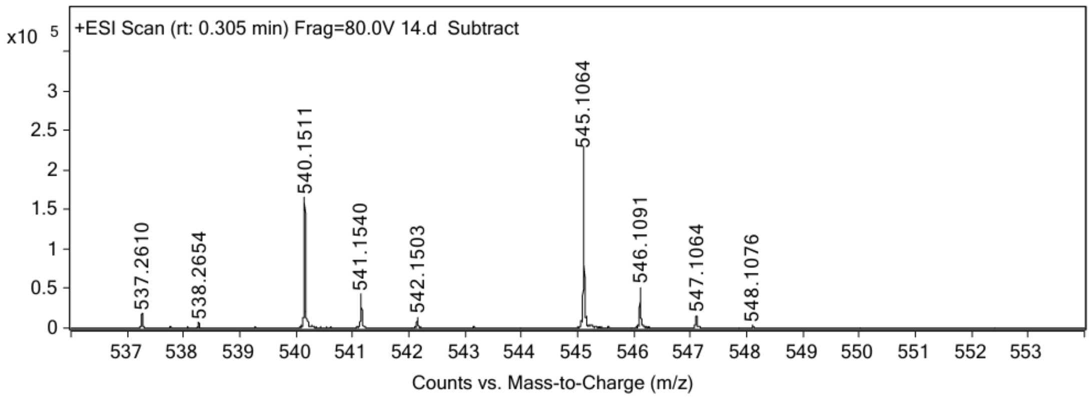

**NBSI-DMAP (16)**

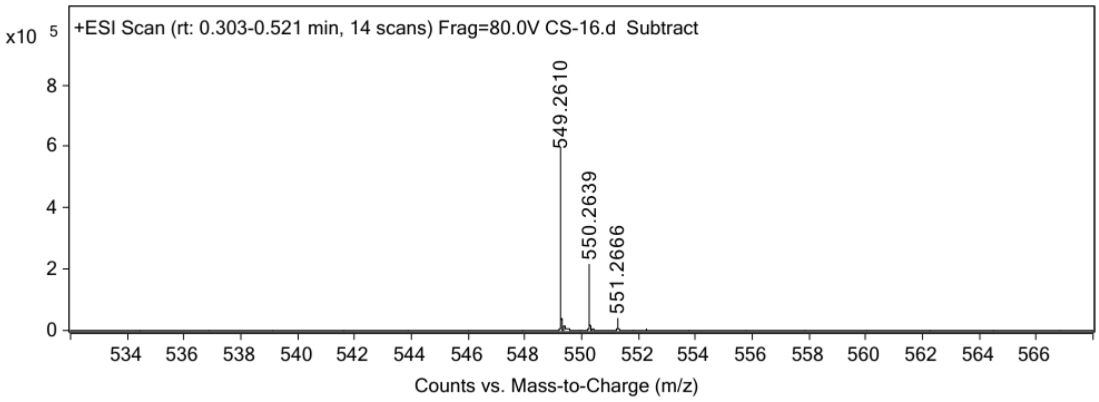


**DFHBI-DMAP (17)**

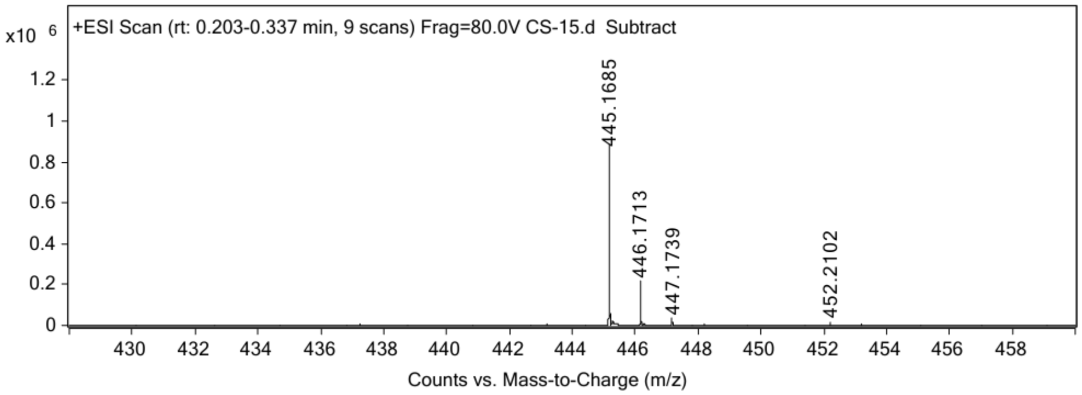


**FMN-DMAP (18)**

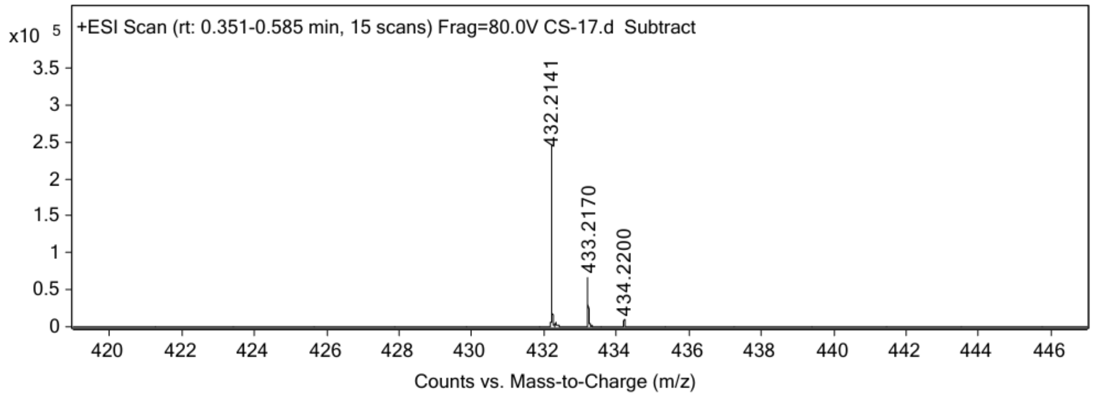

#### 4. Uncropped gels:

The channels outside white box are not discussed in this study.

Figure 2f.

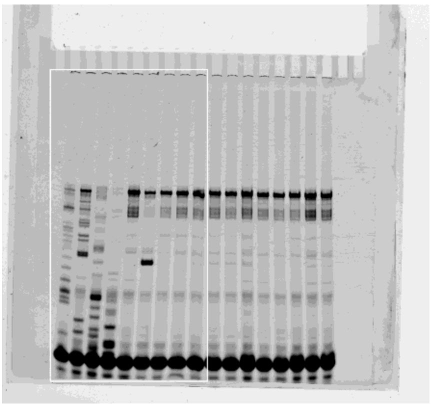

Figure 3d

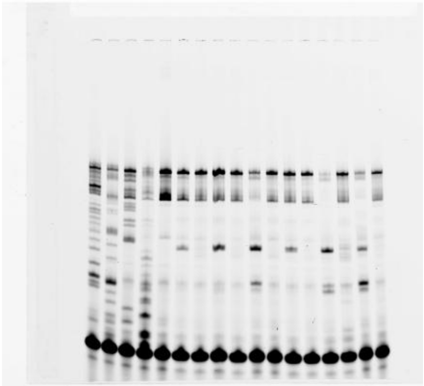

Figure 4b left.

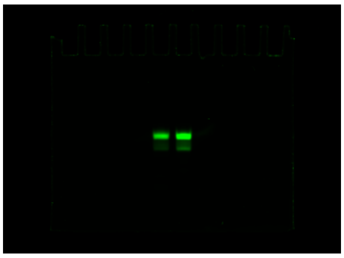

Figure 4b middle.

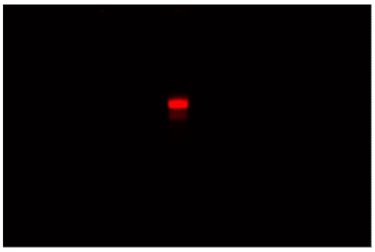

Figure 4b right.

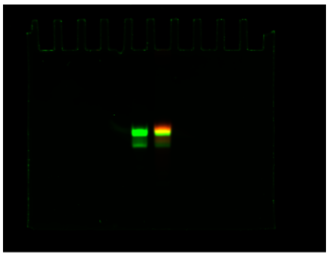

Figure 5d

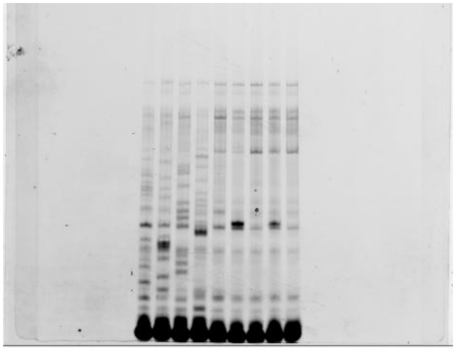

Figure 5h

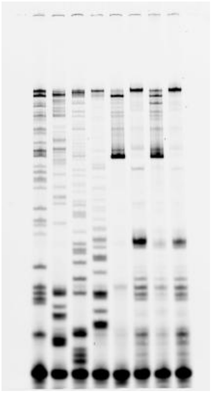

Figure 6a

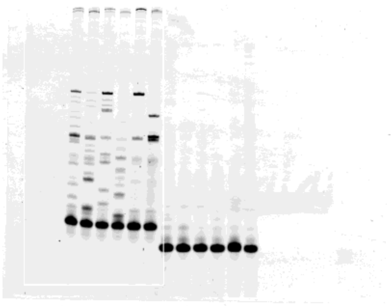

Figure 7b left.

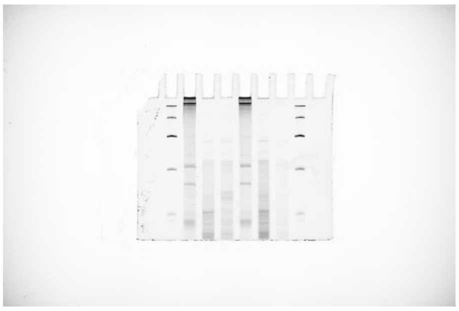

Figure 7b right.

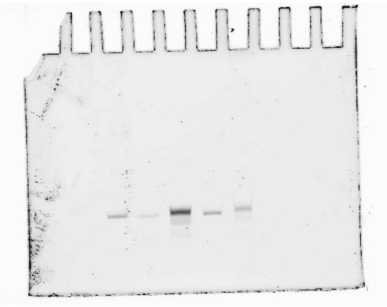

Figure S1c

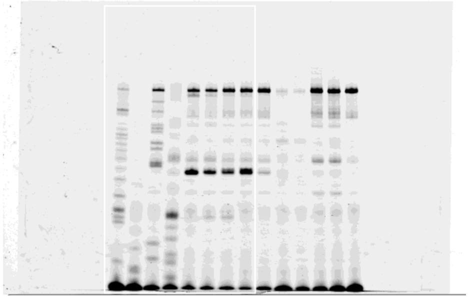

Figure S4c

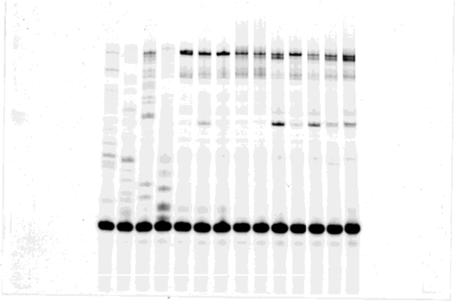

Figure S5b

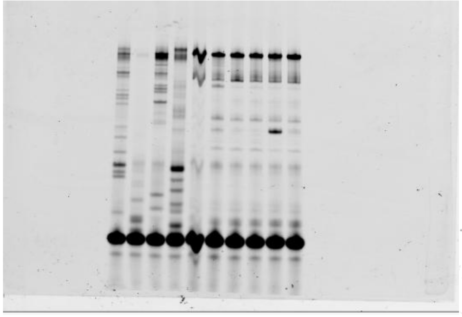

Figure S7e

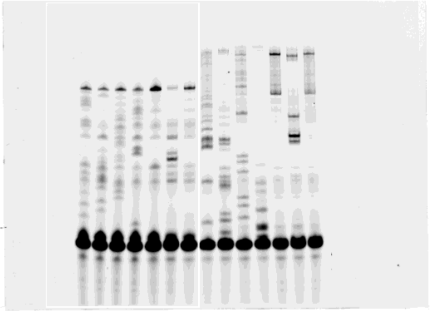

Figure S8

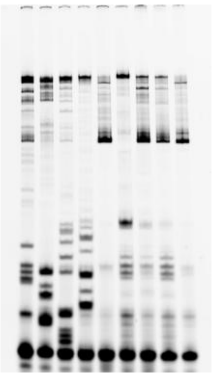

Figure S9c

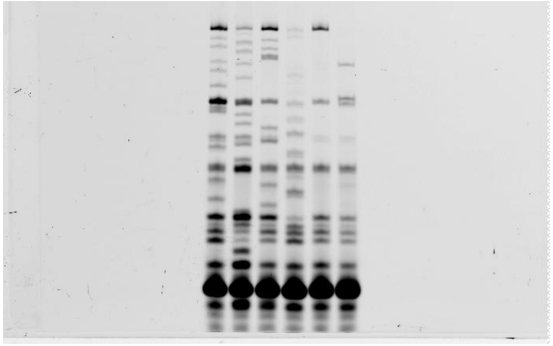

Figure 2f.

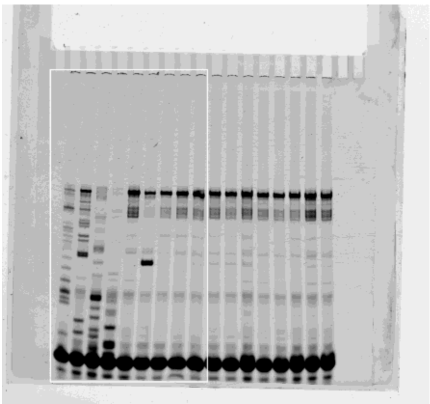

Figure S1c.

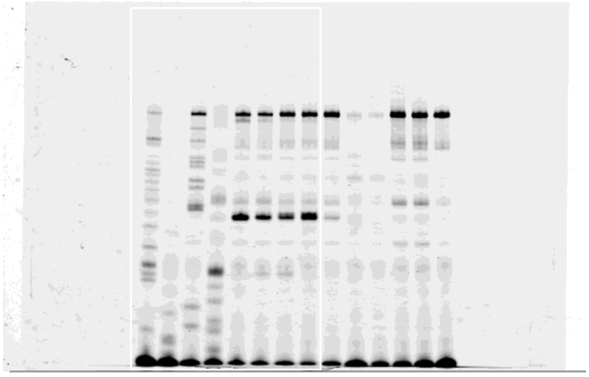

Figure 4b left.

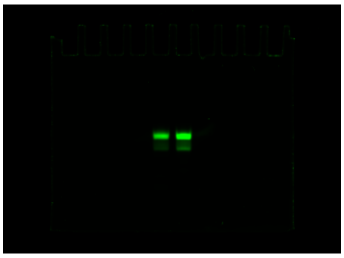

Figure 4b middle.

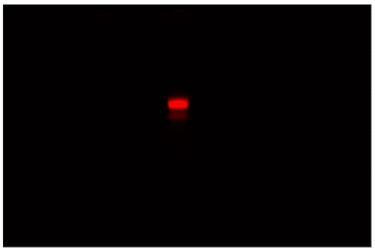

Figure 4b right.

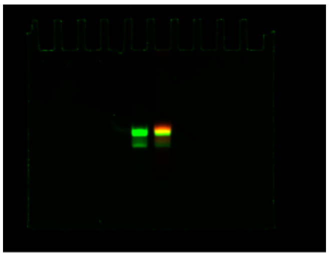

Figure 3d.

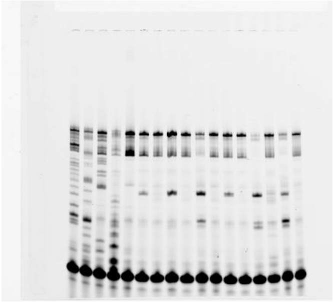

Figure 5d.

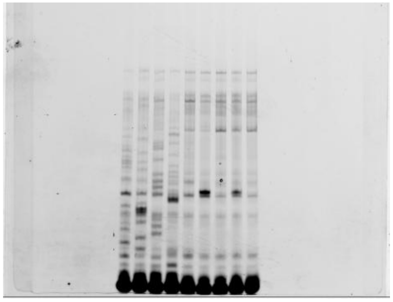

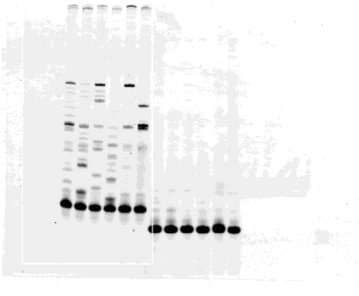

Figure 6a.

Figure 7b left.

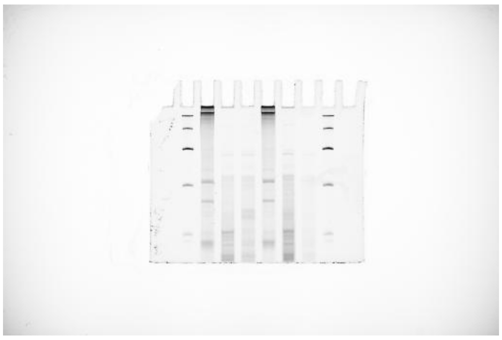

Figure 7b right.

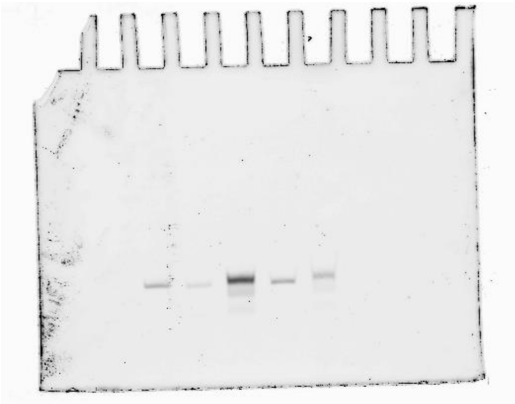

Figure s4c.

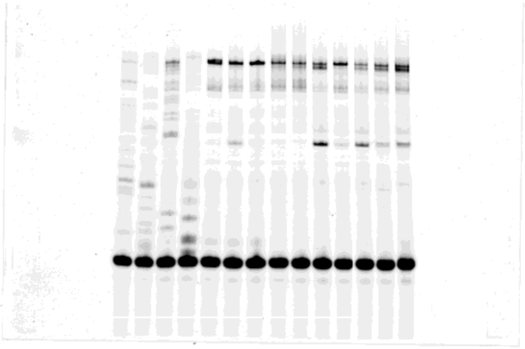

Figure s5b.

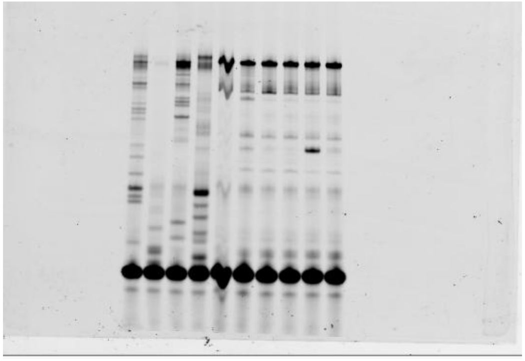

Figure s7e.

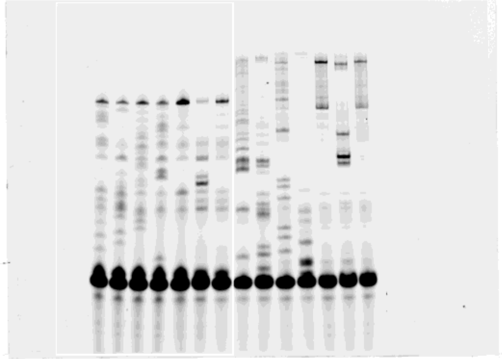

Figure s9c.

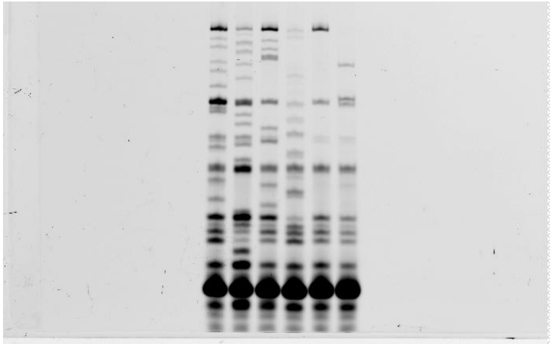

10. Hyun Lee, K.; Kimoto, M.; Kawai, G.; Okamoto, I.; Fin, A.; Hirao, I., Dye-Conjugated Spinach RNA by Genetic Alphabet Expansion. *Chemistry* **2022,** *28* (16), e202104396.
